# Disease-associated variants are enriched for altering cell-type-specific gene co-expression relationships

**DOI:** 10.1101/2025.09.06.674678

**Authors:** Dan Kaptijn, Corinna Losert, Maryna Korshevniuk, Roy Oelen, Martijn Vochteloo, Patrick Deelen, Robert Warmerdam, Anoek Kooijmans, Daniel Considine, BIOS Consortium, sc-eQTLGen Consortium, Yakov Tsepilov, Gosia Trynka, Harm-Jan Westra, Monique van der Wijst, Marc Jan Bonder, Matthias Heinig, Lude Franke

## Abstract

Genes act within complex regulatory networks, and genetic variants can perturb these networks by altering gene co-expression. Here, we performed co-expression quantitative trait locus (co-eQTL) mapping using single-cell RNA-seq from the sc-eQTLGen consortium (1,330 donors, >2 million cells), enabling sensitive detection and prioritization of informative variant–gene–gene triplets.

We identified co-eQTLs for 398 eGenes where a nearby genetic variant affected both the gene’s expression (*cis*-gene) and its co-expression with other genes, often implicating upstream regulators. For 181 genes, we inferred a likely upstream transcription factor, with motif disruption predicted for 41 genes. These upstream genes are more often loss-of-function intolerant and show more network connections, providing an explanation for why co-eQTL variants are 2.8x more strongly associated with immune diseases than classical eQTLs.

These findings position co-eQTLs as mechanistic links between genetic variation and disease, revealing how variants can rewire cell-type-specific gene networks.

## Introduction

Genome-wide association studies (GWAS) have become a key approach for identifying genetic variants linked to complex traits and diseases. To date, hundreds of thousands of genetic variants have been associated with a wide array of phenotypes through GWAS^1^. For the majority of these variants, it remains unclear which genes they affect, in which cell types they act and which pathways or other mechanisms explain their association with a particular trait^2^.

By linking genetic variants to gene expression differences via expression quantitative trait locus (eQTL) analyses, we can establish their potential downstream effects on gene expression. When these analyses are carried out at single-cell resolution, we can also identify whether effects are cell-type-specific, tissue-specific or broadly shared, as has been demonstrated before^3–7^. While many *cis*-eQTLs have now been identified for GWAS variants using this approach, few *trans*-eQTLs have been reported that inform on the downstream pathways that are affected. Furthermore, since it has been shown that *cis*-eQTLs have different characteristics than GWAS variants^8^, it is not fully clear to what extent *cis*-eQTLs help accurately annotate GWAS variants to better understand disease. Complementary strategies to determine the molecular impact of GWAS variants are therefore highly valuable.

Studying how GWAS variants affect gene co-expression networks offer such a complementary perspective. They provide a useful framework for exploring the regulatory mechanisms upstream of eQTL-associated genes (eGenes). This includes the ability to infer direct upstream regulators, e.g. transcription factors (TFs), co-expressed with the eGene, as well as to identify indirect scenarios where co-expressed genes share a common regulator or participate in a shared biological function^9,10^. Moreover, single-cell RNA-sequencing (scRNA-seq) data allow us to construct these co-expression networks for a single individual, providing a unique opportunity to investigate how genetic variation modulates gene co-expression. We call this approach co-expression QTL (co-eQTL) analysis and we infer these effects by testing whether genetic variation alters the co-expression between gene pairs.

In the past, these relationships have been studied using interaction models in bulk data^10,11^ and via small-scale co-eQTL studies in single-cell cohorts (up to 187 donors)^4,6,9^. While bulk studies have mainly been limited by their inability to distinguish signals that affect changes in cell type composition from changes in co-expression, the main limitation of single-cell studies has been insufficient power due to large multiple testing burdens and small sample sizes. The sparsity inherent in single-cell data also makes it challenging to robustly estimate correlations between features and downstream co-eQTL mapping.

To overcome these issues, in this study conducted as part of the sc-eQTLGen consortium, we have carried out the largest single-cell co-eQTL analysis to date. We analyzed five harmonized peripheral blood mononuclear cell (PBMC) single-cell datasets, comprising 1,330 individuals, representing a seven-fold increase compared to previous co-eQTL studies^9^. In addition to increasing statistical power, we systematically optimized the co-eQTL testing framework by thoroughly evaluating the factors that influence co-eQTL detection. We used simulations to test different correlation methods on scRNA-seq data and developed a filtering strategy based on gene characteristics that allowed us to limit testing to the gene pairs where co-eQTL identification is possible.

Our optimized testing framework, combined with meta-analysis and fine-mapping identified 301 unique genes where a nearby genetic variant (*cis*-eQTL variant) influences both the gene’s expression (*cis*-gene) and its co-expression with other genes (co-eGenes). This includes 13,214 credible sets and 12,474 distinct gene pairs across five cell types in PBMCs. The increased number of effects we observe compared to previous studies allowed us to carry out comprehensive downstream characterization and interpretation of these co-eQTLs. By fine-mapping our effects, colocalizing them to GWAS traits and annotating co-eQTLs with functional gene characteristics and TF binding information, we show that our approach detects cell-type-specific and disease-relevant variants that we can connect to regulatory mechanisms upstream of the eQTL effects. Our results suggest that many disease-associated variants act both locally and by altering co-expression relationships between genes, highlighting the complex regulatory consequences of those variants.

## Results

### Overview of the study

To evaluate the extent to which genetic variation influences co-expression, or correlation between gene pairs, we conducted a large-scale co-eQTL analysis using five scRNA-seq PBMC datasets, together comprising 1,330 individuals and approximately 2 million cells (Fig. 1a). We included donors from the OneK1K study^3^ (n = 1,018 donors), an unpublished single nucleus dataset we refer to as UMCG multiome (n = 101 donors), a combined processing of two studies^6,12^ merged but split by chemistry (10x v2 chemistry: UMCG v2, n = 123 donors and 10x v3 chemistry: UMCG v3, n = 47 donors) and the van der Wijst *et al* study^4^ (n = 41 donors). All samples were derived from individuals of European ancestry and consisted of unstimulated PBMCs.

**Fig. 1:**
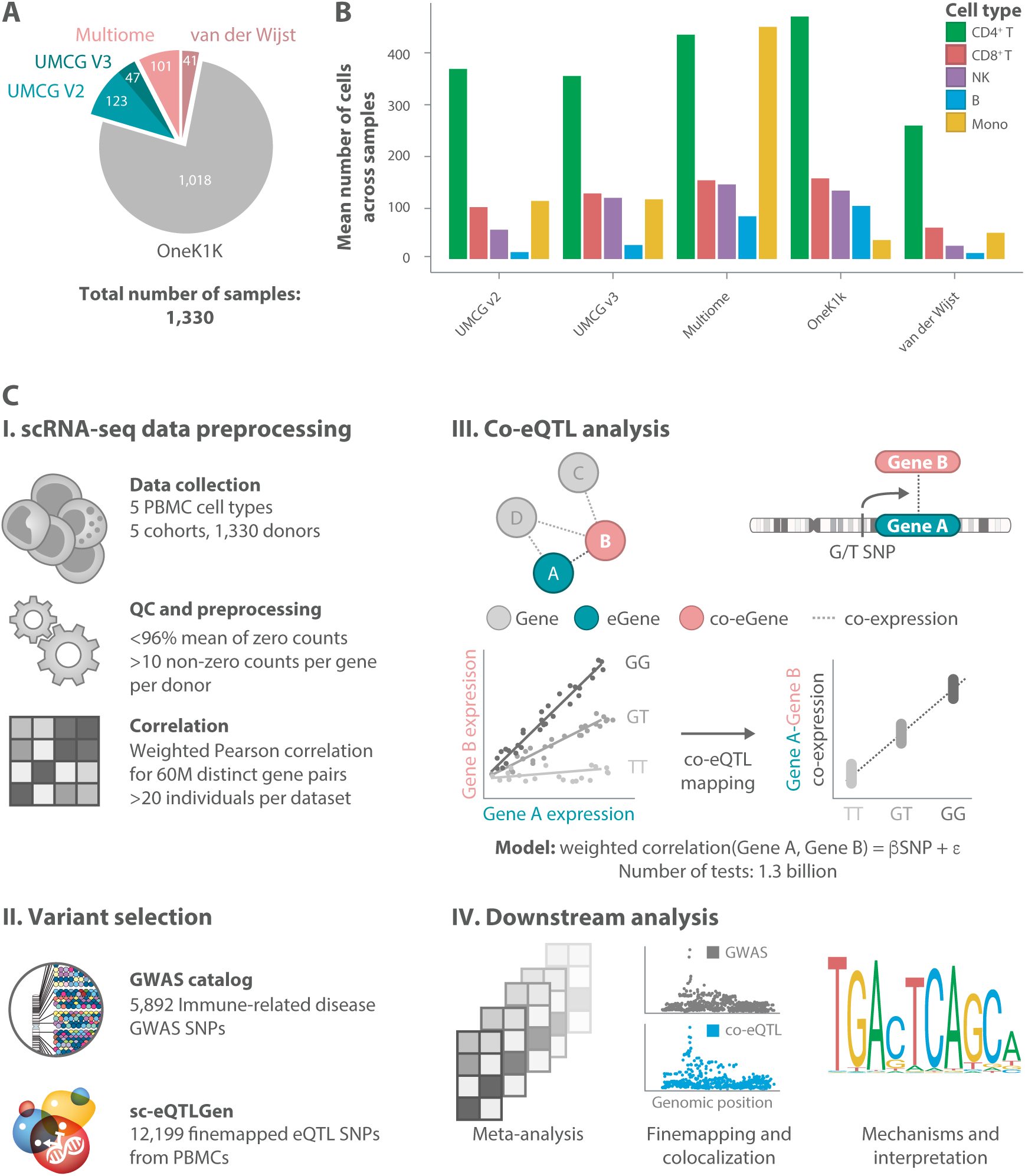
Study overview. a) Overview of the five peripheral blood mononuclear cell (PBMC) single-cell RNA-sequencing (scRNA-seq) datasets used in the co-eQTL mapping in this study, showing the number of donors per study. b) Overview of the mean number of cells per cell type across the five datasets. c) Study design. I. Processing the scRNA-seq datasets. Five pre-processed, harmonized PBMC scRNA-seq datasets from sc-eQTLGen were used. Genes were only included if they had expression in > 4% of the cells per cell type, and correlations were only calculated when there were 10 non-zero counts per gene. Weighted Pearson correlation was calculated for all gene combinations of eQTL genes (eGenes) from sc-eQTLGen and all other genes (co-eGenes) passing the filtering criteria for each individual and cell type. II. Variant selection. For the co-eQTL testing, we selected fine-mapped SNPs from sc-eQTLGen for the corresponding eGene and all immune-related disease GWAS variants in the *cis*-window of ≤ 1MB around the eGene. III. Co-eQTL mapping. Constructed variant–eGene–co-eGene triplets were tested to evaluate the impact of the variant on the co-expression (weighted Pearson correlation) of the gene pair. This identified gene pairs where the genetic variant has a significant impact on gene–gene co-expression. IV. Downstream analyses. Results from the co-eQTL mapping within the five datasets were aggregated through a meta-analysis, and significant effects were fine-mapped. Further downstream analyses included mechanistic annotation of effects.

We focused on five immune cell types (CD4+ T cells, CD8+ T cells, natural killer cells, B cells and monocytes)(Fig. 1b, Suppl. Fig. 1, Suppl. Table 1). Dendritic cells were excluded due to the low number of cells per donor in OneK1K (on average 4 cells per donor). In most of the datasets, CD4+ T cells were the most abundant cell type (average 250–500 cells per individual), followed by CD8+ T cells. In contrast, the most abundant cell type in the UMCG multiome dataset was monocytes (average 470 cells per individual), while monocytes were the least abundant in the OneK1K dataset (fewer than 50 cells per individual on average) (Fig. 1b).

To generate a large set of robust co-eQTLs across all datasets, we employed a two-step process consisting of an evaluation phase in which we optimized the key parameters for co-eQTL detection (Methods) and an application phase in which we applied our refined co-eQTL mapping strategy across all datasets and cell types (Fig. 1c).

#### Optimization of the co-eQTL mapping procedure

Optimization was required to make this project feasible. For example, simply combining all protein coding genes for each of the five assessed cell types would result in 2 billion distinct gene pairs to be tested, and then testing 100s or 1000s of variants in the *cis*-window around each eGene is computationally problematic. Our evaluation phase showed that a weighted Pearson correlation, where observations for each cell are weighted by the number of genes expressed in this cell, performed best for capturing correlations within scRNA-seq data, as determined using simulations based on the structure of OneK1K (Methods, Suppl. Note, Suppl. Fig. 2). Our evaluation also showed that filtering out genes with >96% zero counts across all individuals in a dataset allowed us to reduce the number of tests to be performed by 50% while retaining 99.7% of detected co-eQTLs in CD4+ T cells in the OneK1K dataset (Suppl. Fig. 3).

#### co-eQTL mapping results

Based on the findings from the optimization phase, we defined a set of triplets (variant– eGene–co-eGene combinations) to test for being a co-eQTL. Specifically, we selected independent eQTLs, i.e. the top variant–eGene combinations after fine-mapping in the sc-eQTLGen (manuscript in preparation) project, and filtered the eGenes using the <96% zero count filter (Fig. 1c I). To specifically allow us to study upstream regulation of disease-relevant variants we additionally included all the “immune system disease” variants from the GWAS catalog^1^ (Methods) in the *cis*-window of ≤ 1 mega base pairs (Mb) around an eGene to the *cis*-variant list (Fig. 1c II). We then constructed gene pairs by pairing eGenes with any other gene that passed the < 96% zero count filter to define potential “co-eGenes”. This was followed by computing weighted Pearson correlations for all eGene–co-eGene gene pairs for each individual and cell type separately in order to build individual-specific and cell-type-specific co-expression networks (Fig. 1c I).

This yielded approximately 1.3 billion triplets that were tested for being a co-eQTL across all cell types and datasets (Fig. 1c III). We then conducted a meta-analysis across all the datasets for each cell type, applied multiple testing correction (Methods), and performed a full window scan followed by fine-mapping for gene pairs with at least one significant co-eQTL variant (Fig. 1c IV).

### Meta-analysis identifies co-eQTL effects for 16,124 gene pairs across cell types

After the meta-analysis and multiple testing correction, we identified cell-type-specific co-eQTLs for 1,022 variants (704 unique), involving 398 eGenes (301 unique) and 16,124 gene pairs (12,474 unique), across all cell types (three-step q-value < 0.05; see Methods; Fig. 2a,b, Suppl. Table 2). We observed the highest number of co-eQTLs in CD4+ T cells, where 168 distinct eGenes were associated with co-expression effects in 8,567 gene pairs. B cells had the lowest number of co-eQTLs, with 42 eGenes involving 867 distinct gene pairs (Fig. 2a,b). These numbers of co-eQTLs positively correlated with the number of cells per individual for each cell type (Pearson r = 0.93; Suppl. Fig. 4, Suppl. Table 3). Across all cell types, ribosomal and mitochondrial genes were frequently involved in co-eQTLs, contributing to approximately 55–83% of identified gene pairs and representing 14–23% of the eGenes, depending on cell type (Fig. 2a,b).

**Fig. 2:**
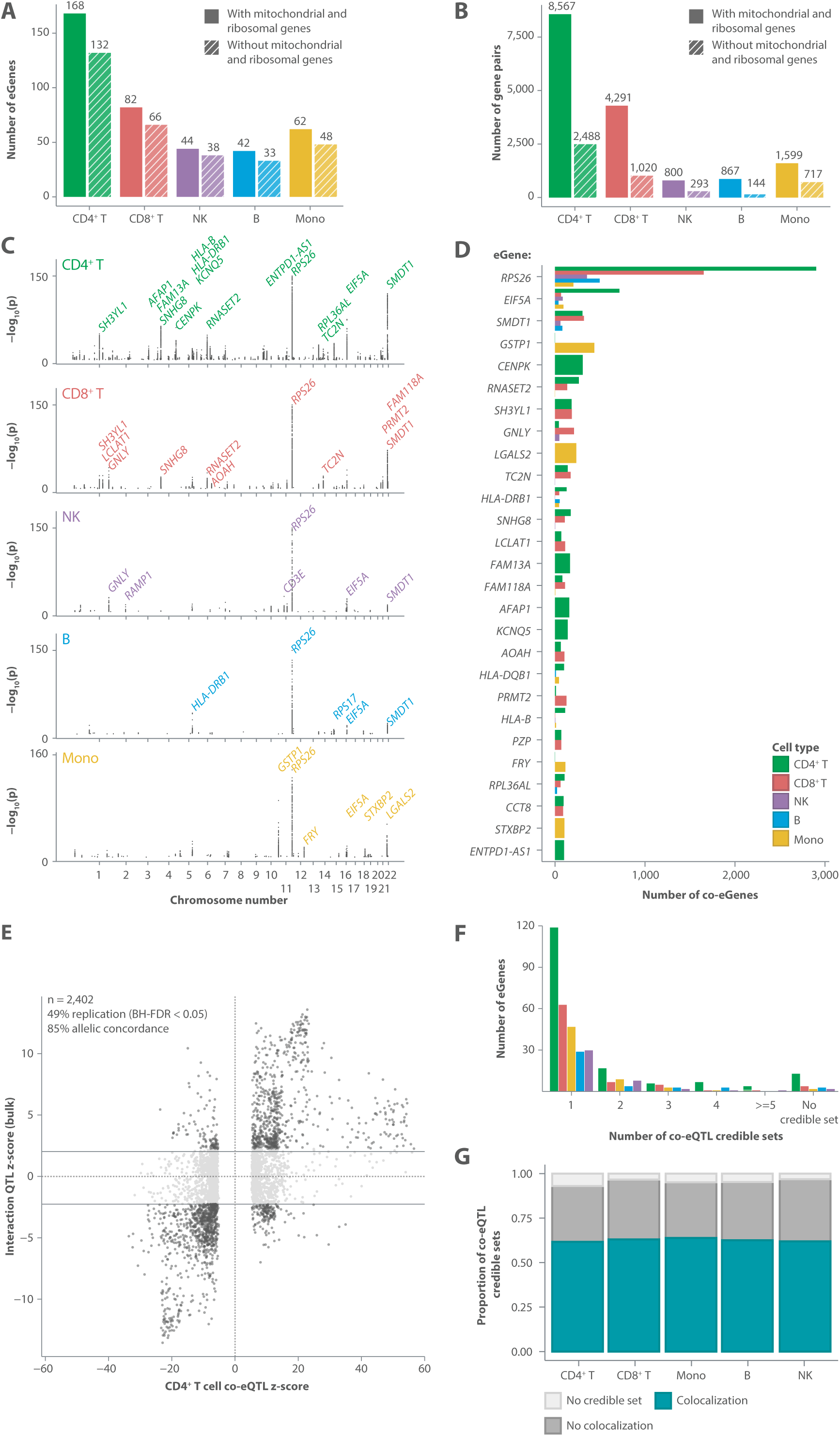
Overview of significant co-eQTL effects. a) Number of eGenes with significant co-eQTL effects within the different cell types after multiple testing correction and meta-analysis. b) Number of gene pairs with significant co-eQTL effects within the different cell types after multiple testing correction and meta-analysis. In a and b two bars are shown per cell type: the left includes all genes, the right has ribosomal and mitochondrial genes removed. c) P-values of significant effects within the different cell types and the location of the variants on the chromosome. Each point reflects a tested gene pair. If there were multiple significant SNPs per gene pair, the entry with the lowest p-value is shown. eGenes with a large number of effects are labelled. d) Number of co-eGenes per eGene and cell type. All eGenes with more than 100 co-eGenes are shown. e) Correlation of z-scores from single-cell co-eQTL mapping in CD4+ T cells and replication of effects in whole blood bulk data from the BIOS study using an interaction term. Effects were included if they were also found in another cell type. Effects were correlated on gene pair level, with the strongest effect per gene pair selected in case of multiple variants. f) Number of co-eQTL credible sets per eGene. Co-eQTL credible sets reflect the independent credible sets across all the co-eGenes of an eGene. g) Proportion of co-eQTL credible sets that colocalize with at least one eQTL credible set.

We further observed substantial variability in the number of co-eGenes associated with individual eGenes. While approximately 50% of eGenes were linked to fewer than 10 co-eGenes (Suppl. Fig. 5), a subset of eGenes exhibit a markedly higher number of associations (Fig. 2c,d, Suppl. Fig. 5). The gene *RPS26* had the most associated co-eGenes (3,511 distinct co-eGenes), which was the case in all cell types except monocytes. In monocytes, the most highly connected eGene was *GSTP1* (437 associated co-eGenes) (Fig. 2c,d). We have previously reported *RPS26*, which encodes a ribosomal protein and shows strong correlations with other genes encoding ribosomal proteins, as the top co-eGene hub^9^. We also identified 26 other eGenes with large co-eGene networks (> 100 connections), including *EIF5A*, *SMDT1*, *GSTP1*, *CENPK* and *RNASET2* (Fig. 2c,d). Similar to *RPS26*, *EIF5A* shows broad co-eQTL associations across all cell types. In contrast to these two highly connected eGenes, other eGenes often displayed more cell-type-specific patterns. *SMDT1*, although a significant eQTL in all cell types, exhibited co-eQTL effects exclusively in lymphoid lineages (between 80 and 322 co-eGenes) and exhibited none at all in monocytes. *GSTP1*, which was also a significant *cis*-eQTL in all cell types except B cells, showed the most co-eQTL effects in monocytes (437 co-eGenes) but only one co-eQTL association in CD4+ T cells. Finally, *CENPK*, a *cis*-eQTL in all lymphoid cell types, showed co-eQTL effects only in CD4+ T cells (308 associated co-eGenes). These observations indicate that co-eQTL regulatory processes of eQTL effects may be more cell type-specific than the eQTL effect itself.

#### Validation of co-eQTLs

To ensure that our co-eQTL results are robust, we performed several different analyses. First, we compared the estimated effect sizes and directions among the five datasets used in the meta-analysis. Here, we observed high correlations when comparing effect sizes (Pearson r = 0.4–0.85) across all the datasets, indicating that the same co-eQTL effects have consistent effect sizes and directions in each dataset (Suppl. Fig. 6). By subsequently applying our meta-analysis approach, we consolidated these effects across datasets. This meant we could detect co-eQTLs with cross-dataset evidence and detect more co-eQTLs than possible when only using the largest dataset (OneK1K) (1,018 individuals vs 1,330 individuals; Suppl. Fig. 7).

Second, we leveraged data from the BIOS study^10^, in which whole blood bulk RNA-seq and genotype data from 2,402 samples were used, to replicate co-eQTLs as interaction-eQTLs. Here, we used the co-eGene as the interaction term in a model to test for association between the co-eQTL variant and the eGene (Methods). We attempted replication per cell type, focussing on the effects linked to independent co-eQTLs (Methods). On average, we could assess replication for 95% of the significant co-eQTL triplets due to study differences. Of these triplets, between 32.9% and 59.7% were significantly replicated, with replication defined as any effect with a Benjamini-Hochberg (BH) false discovery rate (FDR) < 0.05. The average allelic concordances of replicated effects was 83.4% (Suppl. Table 4). As expected, the replication rate and allelic concordance increase when selecting co-eQTL gene pairs found in multiple cell types, given that whole blood is a mixture of the studied cell types. Focussing on these effects we observe replication rates between 49% and 70% and an average allelic concordance of 86% (Suppl. Table 4). For CD4+ T cells, the cell type with most effects, we see a replication rate of 49% and allelic concordance of 85% (Fig. 2e). These results show that many of the co-eQTLs we identify in our study can also be replicated in independent bulk data.

One potential confounder that might introduce false positive co-eQTLs is cell subtype composition. In these cases, co-eGenes specific to that cell subtype may be identified without there being a direct link between the variant and co-expression. To identify co-eQTLs affected by cell subtype composition, we ran a co-eQTL interaction analysis in the OneK1K cells in which we added an interaction term for the proportion of the largest cell subtype per major cell type (Methods). Overall, the results of this analysis indicated that the majority of our co-eQTLs are not likely to be caused by cell subtype effects. For instance, CD8+ T cells showed the highest number of cell subtype effects, with 323 of 4,291 gene pair associations having a significant interaction (BH p < 0.05; Suppl. Tables 2,5). In the other cell types, fewer than 30 gene pairs per cell type were likely explained by such effects (Suppl. Tables 2,5). Additionally, we annotated the co-eQTL variants with the cell count–QTL mapping results of sc-eQTLGen. All effects potentially attributable to cell count effects based on these analyses are marked by the ‘cell count QTL (ccQTL)’ and ‘cell count interaction (ccInteraction)’ columns in in our annotated co-eQTL results table (Suppl. Table 2). Furthermore, we used our simulation approach to annotate co-eQTL effects solely attributable to expression changes (‘simulation_score’ column, Suppl. Table 2, methods).

#### Fine-mapping of co-eQTLs

We conducted several downstream analyses to characterize the observed co-eQTLs. First, to identify likely causal variants and group co-eQTLs that share the same underlying causal variant, we applied joint SuSiE fine-mapping^13^ across all the co-eGenes associated with each eGene, resulting in what we refer to as “gene pair credible sets” (Methods, Suppl. Fig. 8). For most co-eQTL gene pairs, we identified a single gene pair credible set, suggesting that a single causal variant drives the observed changes in co-expression between the two genes (Suppl. Fig. 8a). For a small subset of co-eQTL gene pairs, we detected multiple credible sets, indicating independent regulatory variants. There were also co-eQTL gene pairs for which no credible set could be mapped, likely due to limited power (Suppl. Fig. 8a). Second, we colocalized these signals across co-eGenes of the same eGene. This allowed us to summarize these effects and create “co-eQTL credible sets” (Suppl. Fig. 8). We found 13,214 co-eQTL credible sets in total (Fig. 3d), counting the “independent causal genetic variants” acting on one eGene. The majority of co-eGenes whose expression correlated with the same eGene also colocalized and could be attributed to one underlying credible set (Fig. 2f). The highest number of independent effects among the co-eGenes associated with an eGene was found for *HLA-DRB5* in CD8+ T cells, with 11 distinct credible sets among the different co-eGenes. This is many more than we found for most eGenes (Fig. 2f), which can likely be explained by the highly polymorphic nature of the HLA region, where this gene is located.

**Fig. 3:**
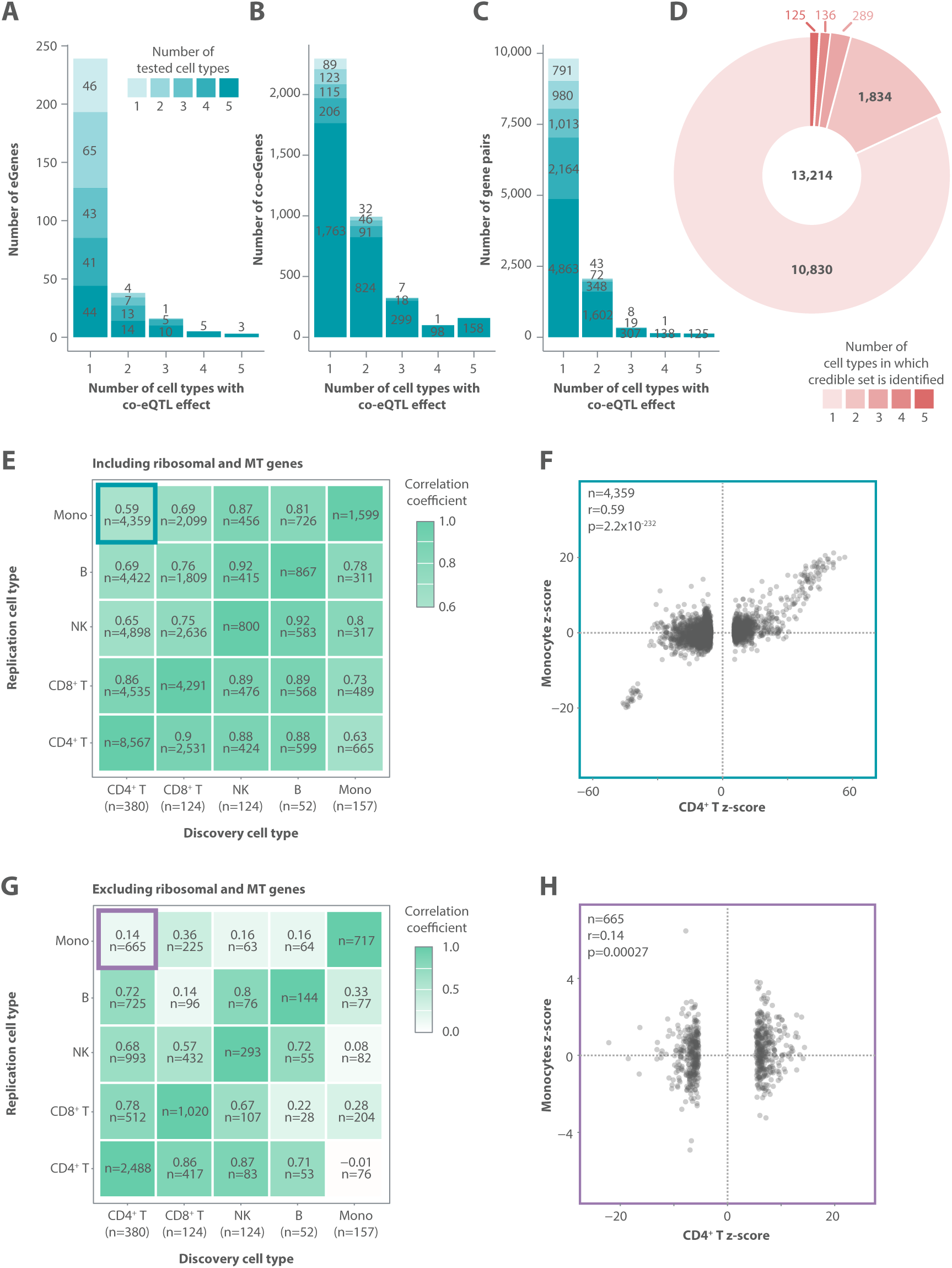
Cell-type-specificity of co-eQTLs. a,b,c) Bar plots visualizing the number of different cell types in which an eGene (a), co-eGene (b), or gene pair (c) had a significant effect (x-axis) compared to the number of different cell types in which it was tested (shades of blue). d) Number of cell types in which a credible set was identified. e, g) Correlation of z-scores across cell types for gene pairs. If there were multiple significant variants per gene pair, the entry with the lowest p-value was used. X-axis indicates the discovery cell type to which the significant gene pairs were correlated with the corresponding gene pairs in the replication cell type. On the x-axis, ‘n’ indicates the average number of cells in this cell type. In heatmap cells, ‘n’ indicates the number of gene pairs that were correlated. In (e), all gene pairs are considered. In (g), only non-ribosomal and non-mitochondrial gene pairs are shown. f, h) Scatterplots for the correlation of z-scores for co-eQTLs identified in CD4+ T cells and tested in monocytes. In (f) all gene pairs are considered. In (h) ribosomal and mitochondrial genes are excluded.

When colocalizing the co-eQTL credible sets to the eQTL credible sets from the sc-eQTLGen fine-mapping of the respective eGene, we observed that, for about 70–90% of the eGenes (depending on cell type), at least one of the co-eQTL credible sets also overlapped with one of the eQTL credible sets (Suppl. Fig. 8b). This shows that the co-eQTL signal often overlaps with the eQTL signal, indicating they likely share the same causal mechanism. Across all cell types, the co-eQTL signal of 51 eGenes did not colocalize with the eQTL signal. These included several co-eQTLs that involved a GWAS variant that also did not show an effect from the eQTL variant. Overall, about 40% of the fine-mapped co-eQTL credible sets did not colocalize with any of the eQTL credible sets (Fig. 2g). This lack of overlap with eQTL credible sets is in line with previous efforts trying to identify upstream regulators of genes via TF-QTLs, where TF-QTLs were identified without a regular eQTL^14–16^.

### Co-eQTLs reveal cell-type-specific interactions

To further characterize our co-eQTL effects, we systematically investigated their cell-type-specificity. First, we analyzed the overlap between eGenes and co-eGenes across cell types. A subset of eGenes (46 out of 301 eGenes) exhibited eQTL effects in only one cell type (Fig. 3a), and consequently showed co-eQTL effects exclusively in that cell type. However, eGenes with eQTL effects across multiple cell types rarely displayed co-eQTL effects in all of those cell types (Fig. 3a–d). In fact, only three eGenes – *RPS26*, *EIF5A* and *HLA-DRB1*– exhibited both eQTL and co-eQTL effects in all the cell types (Fig. 3a). Among the 76 eGenes with eQTL effects across all cell types, 57.8% (n = 44) showed co-eQTL effects in only one cell type (Fig. 3a). This disparity may partly reflect differences in detection power due to varying cell numbers and expression levels across cell types. However, in some cases the co-eQTL was only observed in the cell types with fewer cells, thus supporting the cell-type-specificity observations (Suppl. Fig. 9). We saw similar patterns of cell-type-specificity when analyzing co-eGenes and gene pairs rather than eGenes (Fig. 3b,c, Suppl. Fig. 9), and our co-eQTL colocalization analysis (Methods) confirms this cell-type-specificity. In total, we found 13,214 independent co-eQTL credible sets over the five cell types, with 82% found in only one cell type (Fig. 3d), while just 0.9% are shared across all five. This is much higher than similar estimates based on eQTLs from sc-eQTLGen, where 63% (4,374 of 6,925) of eQTL credible sets are present in one cell type and 5% of eQTL credible sets are shared in all five (Suppl. Table 4). These results show that co-eQTLs are more cell-type-specific than eQTLs.

In addition to assessing the overlap of co-eQTL effects across cell types, we also investigated the allelic concordance of co-eQTL effects tested in multiple cell types (Fig. 3e–h, Suppl. Fig 10). Overall, we observed high concordance, particularly among cells of the same lineage. For example, co-eQTLs identified in CD4+ T cells replicated with the highest concordance in CD8+ T cells (Pearson r = 0.9), whereas the lowest concordance (Pearson r = 0.59) was found in monocytes, a cell type from another lineage. Cell-type-specificity also increased when ribosomal and mitochondrial genes were excluded (Fig. 3 g,h). Specifically, effects identified in CD4+ T cells become more different from effects identified in monocytes which is shown by a drop in concordance from a Pearson r of 0.59 to 0.14. This suggests that co-eQTLs involving ribosomal and mitochondrial genes are more commonly shared across cell types than those involving other co-eGenes.

### Co-eQTLs show greater disease-relevance than eQTLs

Beyond examining the cell-type-specificity underlying co-eQTLs, we aimed to identify further properties of co-eQTLs. For these analyses, we focused on eQTL linked effects, i.e. removing the co-eQTL effects linked purely to GWAS variants.

First, we compared eGenes and variants as part of a co-eQTL or classical eQTL effect and found that these two groups exhibit distinct properties. In this analysis, the eGenes and variants were overlapped with GWAS signals in two ways: using GWAS annotations from the GWAS catalog^1^ (variant-based) and using colocalization results (eGene-based) (Methods). Considering all cell types together, we found that eGenes and variants with co-eQTL effects were significantly enriched for disease-related GWAS traits when compared to eGenes and variants as part of a classical eQTL effect (Fisher exact test: eGene-based all cell types: OR = 1.57, p = 0.017; variant-based all cell types: OR = 1.53, p = 0.019) (Fig. 4a). While the results were not always significant in the less abundant cell types due to limited statistical power, all the ORs exceeded 1, suggesting co-eQTL enrichment for GWAS variants in these cell types.

**Fig. 4:**
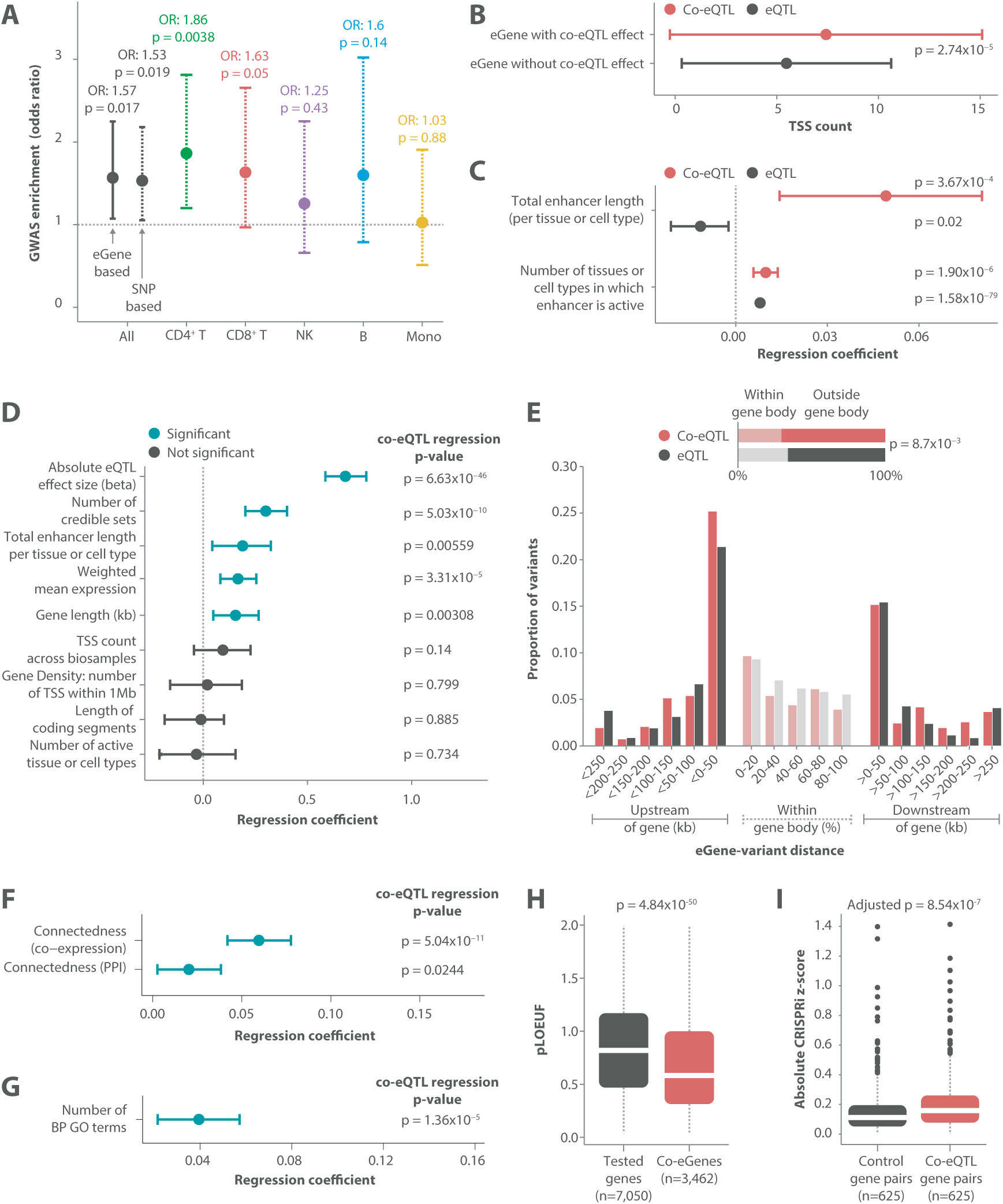
Co-eQTLs show greater disease-importance compared to eQTLs. a) Odds ratio (OR) of enrichment of disease-related GWAS terms (based on EFO classification, see Methods) for variants (variant-based) and eGenes (eGene-based) with co-eQTL effect compared to *cis*-variants and *cis*-eGenes without a co-eQTL effect. b) Transcription start site (TSS) count showing the mean count of TSS per gene across cell types in the FANTOM project for eGenes with a co-eQTL effect compared to eGenes without co-eQTL effect. c) Enhancer features: logistic regression coefficients for enhancer features derived from the Roadmap dataset (total enhancer length, count of active tissue/cell types) for predicting co-eQTL hits (red) and eQTL hits (grey) versus random variants (n = 100,000) after adjusting for confounders (Methods). d) Logistic regression coefficients in a model predicting co-eQTL hits vs. eQTL hits across all cell types. e) Comparison of the location of genetic variants between eQTLs and co-eQTLs. Variants outside the gene body are summarized into 50 kb bins. Variants inside the gene body are summarized relative to the position inside. Inset figure shows the difference in the number of variants found in the gene body. f) Regression coefficients of linear models on the residuals of predicting the connectedness of significant co-eGenes vs. tested genes after correcting for the mean expression. g) Regression coefficients of linear models on the residuals of predicting the number of BP GO terms for significant co-eGenes vs. tested genes after correcting for the mean expression. h) LOEUF scores for significant co-eGenes vs. tested genes, showing that co-eGenes are more conserved. i) Average absolute CRISPRi z-scores for co-eGenes separated into co-eQTLs where the gene pair is a co-eQTL and mismatched co-eQTLs where both genes are part of significant co-eQTLs but the gene pair is not a co-eQTL. In total, 625 co-eGenes are shown. P-value is based on 100 permutations of mismatched co-eQTLs.

We also tested several subsets of GWAS traits by filtering the traits to exclude cell count traits and focussing on disease-specific and immune-disease-specific traits (Suppl. Fig. 11). When restricting to immune-related-traits ORs increased (eGene-based all cell types: OR = 1.87, p = 0.052; variant-based all cell types: OR = 2.78, p = 0.00066) (Suppl. Fig. 11, Suppl. Tables 5–8.).

Overall, these results suggest that co-eQTL variants contribute to disease susceptibility beyond classical eQTL effects, highlighting the importance of investigating co-eQTLs as a complementary layer of genetic regulation relevant to complex traits.

#### Co-eQTL eGenes and variants show higher regulatory complexity than classical eQTLs

We next investigated if co-eQTL eGenes and related variants showed other similarities to GWAS variants when compared to classical eQTL eGenes and variants. To do so, we examined their transcriptional regulatory landscape using metrics proposed by Mostafavi *et al*^8^. Using data from the FANTOM project^17^, we observed that eGenes with a co-eQTL effect contain significantly more transcription start sites (TSS) than those without a co-eQTL effect (mean TSS = 7.4 vs 5.5, respectively; p = 2.74*10^−5^; Fig. 4b, Suppl. Fig 12a). Additionally, we applied the model established by Mostafavi *et al*^8^ to compare enhancer features (enhancer length and number of active tissues/cell types) based on enhancer–gene links derived from the Roadmap Epigenomics Consortium dataset from Liu *et al*^18^ (Methods). Both co-eQTL and classical eQTL variants showed broader enhancer activity than random variants. However, variants with co-eQTL effects were linked to significantly longer enhancer regions when compared to classical eQTLs (regression coefficient = 0.05, p = 0.0037 vs regression coefficient = −0.01, p = 0.0169); Fig. 4c, Suppl. Fig. 12b). These findings are in line with the differences observed between GWAS hits and eQTL hits by Mostafavi *et al*^8^, and they suggest that co-eQTL variants have a more complex regulatory landscape than classical eQTL variants.

When looking at eQTL fine-mapping results from sc-eQTLGen for eGenes with co-eQTL effects, we find they have more credible sets compared to eGenes that are part of classical eQTL effects (regression coefficient = 0.3, p = 5.03*10^−10^; Fig. 4d, Suppl. Fig. 13). This indicates that these co-eQTL eGenes are usually driven by multiple variants, reflecting a complex genetic architecture underlying the gene’s expression. We also observe that the number of credible sets for an eGene positively affects its colocalization with GWAS traits (regression coefficient = 0.13, p = 0.00158; Suppl. Fig. 14). This is thus one of the drivers of the observed GWAS enrichment of the co-eQTL results, as the credible set number affects both the likelihood of detecting a co-eQTL and the likelihood of GWAS colocalization (Suppl. Figs. 11 and 14). The number of credible sets for an eGene was also positively correlated with the number of co-eGenes associated with an eGene (Spearman r = 0.27, p = 9.8*10^−12^) (Suppl. Fig. 15), showing that eGenes with more potential independent causal variants also tend to have more associated co-eGenes. Taken together, these findings again suggest that co-eQTL eGenes and variants are embedded in a more complex regulatory environment than classical eQTLs, potentially facilitating broader phenotypic consequences.

When looking at the location of the variants, we observed a depletion of co-eQTLs within the gene body compared to classical eQTLs (OR = 0.813, two-sided Fisher exact test p = 0.0087) but no differences outside of the gene body (Wilcoxon rank sum test: upstream of the gene p = 0.13, downstream of the gene p = 0.10) (Fig. 4e). These observations are in line with our expectation that co-eQTLs more often capture gene regulatory effects occurring within regulatory regions that are often located outside the gene body. These findings are also concordant with previous literature showing that GWAS-implicated variants are less often found in the gene body^19^.

#### Co-eQTL co-eGenes have widespread regulatory influence

Beyond observing higher regulatory complexity for the regulatory variants and eGenes that are part of a co-eQTL effect, we also noted several functional distinctions for co-eGenes compared to the genes tested but not found significant. For this analysis, we again used several features constructed by Mostafavi *et al*^8^. First, we observed that co-eGenes tend to have higher connectivity in the co-expression networks inferred by Saha *et al* for GTEx tissues^20^ (regression coefficient = 0.06, p = 5.04*10^−11^) and in InWeb Protein Protein Interaction networks^21^ (regression coefficient = 0.02, p = 0.0244) (Fig. 4f, Suppl. Fig. 16). This suggests that co-eGenes function as regulatory hubs within gene regulatory networks, and such hubs have been reported to have higher importance for biological processes^22^. Co-eGenes are also enriched for genes under strong selective constraint, as shown by lower LOEUF scores from gnomAD^23^ (two-sided t-test p = 4.84*10^−50^) (Fig. 4h, Suppl. Fig. 16). Co-eGenes also show greater functional annotation, with significantly more Gene Ontology terms^24,25^ per gene (regression coefficient = 0.04, p = 1.36*10^−5^) (Fig. 4g, Suppl. Fig. 16). Combined with our other results, this indicates that co-eGenes are functionally important, potentially due to their widespread regulatory influence.

To further validate our co-eQTLs and their regulatory potential, we used the genome-scale Perturb-seq CRISPRi dataset from Replogle *et al*^26^. To test perturbation strength, for each perturbation of a co-eGene, we compared the perturbation impact between two equal-sized sets of eGenes: those that were and were not part of a specific co-eQTL pair (Methods). This analysis showed that eGenes belonging to a co-eQTL gene pair are more impacted by perturbation of the co-eGene when compared to gene pairs that did not show a co-eQTL effect (Wilcoxon signed-rank p = 8.54*10^−7^) (Fig. 4i, Supplementary notes).

### Co-eQTLs reveal regulatory mechanisms of disease loci

Lastly, we investigated specific upstream regulatory mechanisms underlying observed co-eQTL effects and their association to disease. We reasoned that the most likely co-eQTL mode of action would be represented by gene regulation via TFs (TF = co-eGene), where the co-eQTL variant (the variant in *cis* with the eQTL gene) disrupts the TF’s binding site, thereby modulating the TF’s regulation of the eGene. In such cases, the correlation between the eGene and co-eGene may reflect this regulatory interaction. To explore this, we annotated TFs among the co-eGenes, either directly finding a TF among the co-eGenes or finding over-representation of TF-binding sites in the promoter regions of the co-eGenes. The latter strategy was applied to include TFs with expression levels too low to be detectable in scRNA-seq data and those with activity that is regulated post-translationally. For each of the potential TF–eGene pairs we identified, we used MotifbreakR^27^ to assess whether the fine-mapped variants disrupt the known TF-binding motif for that TF (Methods).

Among the 398 eGenes we identified across all cell types, 151 (37.9%) had at least one associated co-eGene annotated as a TF (‘direct TF’), 30 (7.5%) showed enrichment of at least one TF among the co-eGenes (‘direct-enriched TF’), and 48 (12%) were supported by both lines of evidence (Fig. 5a I, Suppl. Fig. 17). In the MotifbreakR analysis, 37 direct and four direct-enriched TFs showed significant binding motif disruptions, strongly suggesting a direct regulatory role for the co-eGene TF on the corresponding eGene (Fig. 5a I, Suppl. Fig. 17). Looking from the TF side, we identified 163 (252 across cell types) TFs as potential upstream regulators of co-eQTLs; 68 of these are not a *cis*-eQTL in sc-eQTLGen. Next to TF effects, there are other regulatory mechanisms that might be represented by co-eQTL effects. One of these is regulation by microRNAs. Using the enrichment strategy outlined above, we found that for 53 of the 398 eGenes, the co-eGenes were enriched for microRNAs that regulate their expression (Suppl. Fig. 18).

**Fig. 5:**
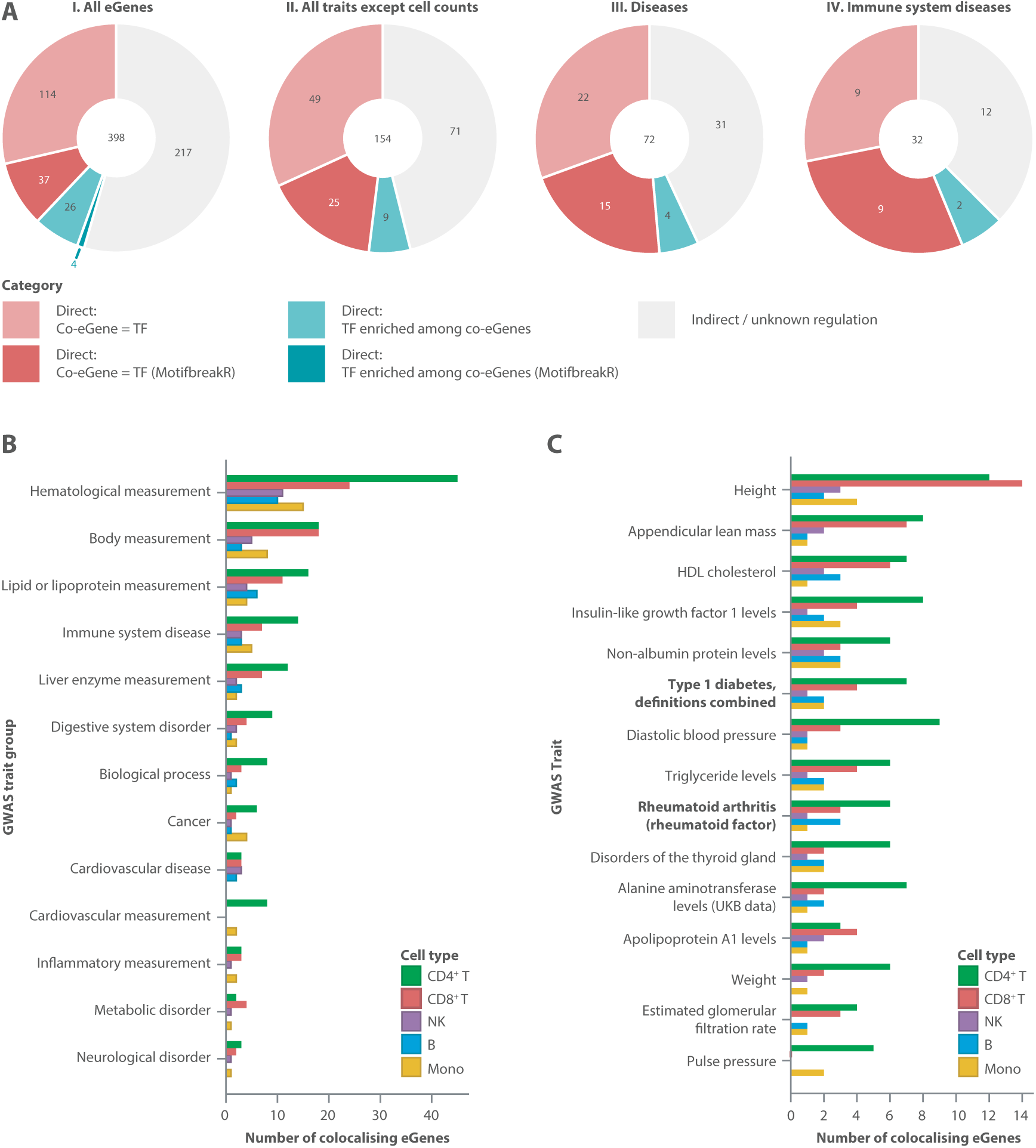
Co-eQTLs show gene networks relevant to disease interpretation. a) Number of eGenes filtered on their implication to several GWAS categories with identified likely transcription factor (TF), separated on whether the TF is part of identified co-eGenes (red) or identified through enrichment analysis of identified co-eGenes (blue). Darker colour indicates the TF has corresponding evidence from MotifbreakR. Panels from left to right: I. all 398 eGenes identified, II. eGenes associated to a non-cell count GWAS trait, III. eGenes associated to disease GWAS traits, and IV. eGenes associated to “immune system disease” GWAS traits. b) Number of eGenes colocalized versus the parent terms of GWAS traits, sorted by the number of colocalizing eGenes and numbers per cell type. c) The 15 GWAS traits with the highest number of colocalizing eGenes, including number of colocalizing eGenes per cell type.

To put the regulatory effects captured by co-eQTLs into disease context, we used the fine-mapping results of the co-eQTLs (Methods) to colocalize the co-eQTL effects with GWAS traits. In all, 46% (n = 184) of the eGenes with a co-eQTL effect colocalized with at least one GWAS trait (Suppl. Fig. 17). When filtering co-eQTLs based on whether they showed colocalization to particular GWAS categories (Fig. 5ai-aiv), the proportion of eGenes for which we have identified the likely TF is consistent with what we see over all eGenes. In fact, as the categories become more specific i.e. moving from all GWAS traits to disease-specific traits to immune-mediated traits (Fig. 5A I-IV), the proportion of eGenes that we can annotate an upstream regulator for increases, rising to 62.5% for immune-mediated traits. While the highest number of eGenes are connected to measurement traits, immune system disorder is one of the overall terms for which we see colocalizations in every cell type (Fig. 5b). When looking at specific traits, height is connected to the most genes, due at least in part to the high number of loci implicated in height GWAS. For diseases, type 1 diabetes and rheumatoid arthritis show multiple associations (Fig. 5c).

Interestingly, several colocalizations were unique to co-eQTLs, i.e. they were not identified in the eQTL colocalization results. Some of these effects could be attributed to the inclusion of GWAS variants, i.e. non-eQTL signals, in our mapping strategy (Methods, Suppl. Fig. 17). For example, the triplet *IL7R* (eGene) – *ID2* (TF) – 5:35877403:G:A (variant) (Suppl. Fig. 19) in CD4+ T cells colocalized with dermatitis only for co-eQTLs. Interestingly, altered expression of *IL7R* as a consequence of *ID2* deficiency in “Lymphoid Tissue inducer like” cells has been reported previously^28,29^. The eGene *IL7R* was also previously linked to atopic dermatitis by various GWAS^30–32^, while the co-eGene *ID2* was linked to dermatitis in a study showing that loss of *ID2* (and *ID3*) in regulatory T cells leads to dermatitis-like disease in mice^33^. Another example is the triplet *TSPAN32* (eGene) – *CCL5* (TF) – 11:2327389:T:C (variant) in CD8+ T cells, which colocalized to type 2 diabetes. There is evidence for *TSPAN32* involvement in gestational diabetes mellitus^34^, and *CCL5* has also been linked to type 2 diabetes^35^. These two examples illustrate that co-eQTLs can uncover novel trait associations that were not detected via classical eQTL mapping.

We next specifically investigated the examples with strong TF-binding support throughout our analysis. For the eGene *RNASET2*, we identified 11 different TFs among the co-eGenes, of which *LEF1* stood out. The co-eQTL effect was with the variant 6:166974681:A:G (Fig. 6a). This variant is located 17,490 bases upstream of *RNASET2* in a motif for *LEF1*, and MotifbreakR shows evidence that the A allele disrupts *LEF1*’s ability to bind to the motif (Fig. 6a,b,c). This is in line with the repressive regulation shown by the co-eQTL (Fig. 6a) of *RNASET2* by *LEF1* with the GG genotype of variant 6:166974681:A:G (r = −0.0379) and a disruption of this repression for the AA genotype in CD4+ T cells (r = −0.006). Interestingly, the G allele, which is the allele matching the motif, is the risk allele for three diseases in the colocalization analysis: Crohn’s disease (beta = 0.1477, p = 1.8*10^−20^, study: GCST004132), rheumatoid arthritis (beta = 0.0787, p = 1.1*10^−9^, study: GCST90018910) and Grave’s disease (beta = 0.1152, p = 2.5*10^−8^, study: GCST90018847) (Fig. 6d). When comparing across datasets (Fig. 6e) and cell types (Fig. 6f) we see that the relationship for this co-eQTL is consistent, and *LEF1* expression was only high enough to be able to calculate a co-eQTL effect in T cells. These results imply that the *LEF1* regulation of *RNASET2* is important in T cells within autoimmune contexts, consistent with earlier observations^36^.

**Fig. 6:**
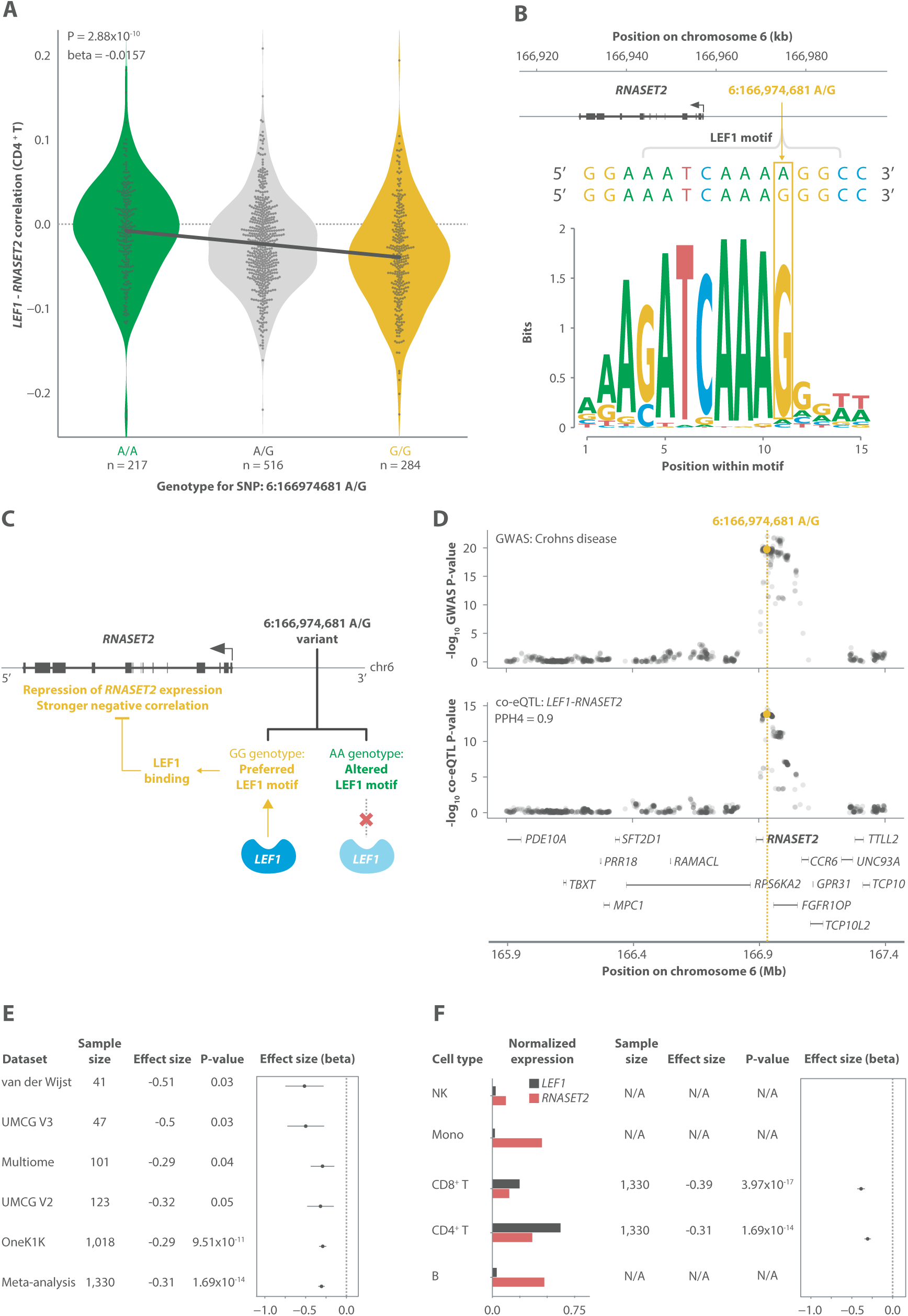
Co-eQTL effect of RNASET2–LEF1 in CD4+ T cells. a) Co-eQTL plot showing the co-eQTL effect size based on effect sizes in OneK1K. b) Motif plot from MotifbreakR. G is shown to match the preferred binding motif for LEF1, whereas A breaks the motif. c) Representation of co-eQTL effect. LEF1 acts as a repressor of *RNASET2* expression, but can only bind when the variant is the G allele and therefore is unable to repress *RNASET2* expression when the variant is the A allele. d) Locus zoom plot showing both the Crohn’s disease GWAS and co-eQTL signals in the locus. The variant 6:166974681:A:G is highlighted in yellow. A strong colocalization is observed between these signals (0.9). The protein-coding gene landscape is shown below the plots. e) Effect size of the *RNASET2*–*LEF1* co-eQTL in each dataset and in the meta-analysis. Effect sizes shown are from the main analyses not the raw correlation shown in (a). f) Normalized expression and co-eQTL effect sizes of *LEF1* and *RNASET2* in each cell type.

For the eGene *RPS26*, we identified 145 TFs among the co-eGenes. A special case here was the gene pair *NR4A3*–*RPS26* affected by variant 12:56007301:G:A (Suppl. Fig. 20), for which we observed an opposite effect in monocytes and CD4+ T cells. We detected a strong positive correlation (r = 0.6352) between *NR4A3* and *RPS26* in monocytes for the GG genotype but a negative regulation in CD4+ T cells (r = −0.1934). This indicates an activator role of *NR4A3* on *RPS26* in monocytes but a repressor function in CD4+ T cells. In both cases, the regulation is disrupted by the AA genotype (Supp. Fig. 20), aligning with the strength of the *NR4A3* motif. It was previously reported that *NR4A3*, a member of the nuclear receptor family, can play opposing roles in myeloid and lymphoid cell lines, depending on context^37–40^. This effect was found to colocalize to both rheumatoid arthritis (study: GCST90132225) and type 1 diabetes (study: GCST010681), and both *RPS26* and *NR4A3* have been associated with type 1 diabetes by GWAS^41–43^. However, the variant (12:56007301:G:A) is implicated with opposing effect signs in the traits. Both the disease-association and dual role for *NR4A3* point to complex regulation and context-dependence identified through co-eQTLs for this locus.

Other strongly supported TF–eGene regulations include the pair *SH3YL1*–*IKZF2* colocalizing to metabolic syndrome (Suppl. Fig. 21) and *AP1S3*–*BACH2* colocalizing to cystatin C production (Suppl. Fig. 22). In both cases there is a motif present for the relevant TF and, according to MotifbreakR, the allele showing no correlation is also the allele breaking this motif. We also see that these effects are consistently identified across all five datasets, indicating the robustness of these associations.

Taken together, these examples illustrate that co-eQTLs can both elucidate the upstream mechanisms behind known disease-associated eQTLs and help uncover novel trait associations and contexts.

## Discussion

This study is the largest co-eQTL analysis to date, leveraging five harmonized single-cell datasets from the sc-eQTLGen consortium. This analysis includes over 2 million immune cells from 1,330 donors, providing unprecedented resolution to map co-eQTLs across diverse cell types and contexts. Using this data, we identified co-eQTLs for 398 eGenes, constituting 16,124 gene pair interactions. These co-eQTLs capture upstream regulatory mechanisms of eQTLs by linking eGenes to co-eGenes, which together form clusters of gene regulation. Importantly, the increased power of our analysis allowed us to imply upstream regulators for 45.5% of eGenes that show a co-eQTL effect.

We used two different methods to assess the disease-relevance of these co-eQTLs: gene- and variant-based enrichment against a background set of classical eQTLs. In both cases, we found an enrichment for disease variants underlying co-eQTLs, showing that co-eQTLs are important in disease pathophysiology. We also found that co-eQTL variants share characteristics with GWAS variants that classical eQTLs do not. Similar to the observations of Mostafavi *et al*^8^, we find that classical eQTLs tend to be in locations with a low abundance of enhancers, whereas GWAS and co-eQTL variants tend to be in areas with more enhancer sites and higher activity. This can partly be explained by the number of independent effects observed for a classical eQTL, i.e. those with more independent effects have a higher chance of both being a co-eQTL and a GWAS-implicated variant. This suggests that regions with complex regulation, where there is evidence for multiple independent regulators, are more disease-relevant and that these regions are captured by co-eQTLs.

These findings are also supported by the disease colocalizations that have been identified from the co-eQTL summary statistics, which yield both genes and gene regulators. These regulators are a key mechanistic element for understanding eQTL associations, and they enable the construction of gene networks. Different strategies have previously been applied to detect the potential upstream regulators of eQTL effects, e.g. through correlation QTL analyses in scRNA-seq data^14^ or interaction QTL analyses in bulk RNA-seq data^15,16^. These strategies have revealed thousands of independent variants that can affect the co-expression between TFs and their potential target genes. Another approach would be to compare co-eQTL gene pairs to *cis*-*trans* pairs identified through *trans*-eQTL studies such as eQTLGen (manuscript in preparation). Analysis of genetic variants affecting cell state abundance in single cell data^44^ can also pinpoint larger groups of co-expressed genes and pathways associated with a variant by exploring the gene signatures that define these granular cell states. However, each of these approaches have different strengths and weaknesses, including our own. Compared to the first group of methods, our broadened (but optimized) search space allows for potential detection of regulatory events beyond TFs. Compared to *trans*-eQTLs, our co-eQTL methodology allows us to identify upstream regulators not directly under the regulation of common variants. Interestingly, we observe that co-eGenes are indeed more constrained (loss of function intolerant), a factor previously shown to limit the likelihood of identifying a gene as a *cis*-eQTL^45^, and more than a third of the co-eGene TFs we identified are not *cis*-eQTLs. Relative to both alternative approaches, co-eQTLs capture TF target genes as a proxy for TF activity, information essential to identify the upstream regulator when the TF’s expression would not reflect the TF’s activity. This is something that limits both of the alternative methods discussed above. Given these points, it is likely that using our co-eQTL approach we would identify a broader set of regulatory effects (at the cost of a higher multiple testing burden), and by leveraging single-cell information we have reduced the likelihood of cell type composition artefacts. However, co-eQTL mapping would not identify all *cis*-*trans* eQTL pairs as expression changes that do not change the co-expression relation between the two genes will not be identified through co-eQTLs. Compared to the cell state abundance approach, which identifies pathway level gene signatures, the co-eQTL results provide more detailed information about regulatory interactions and potential molecular mechanisms. Therefore, all these approaches have their own value and are complementary to discovering how genetic variants impact gene regulation.

One of the main limitations of our current setup is the validation of our co-eQTL effects. Replication via both interaction analysis on bulk RNA-seq and CRISPR are not ideal. Bulk RNA-seq suffers from cell type composition effects present in whole blood cell mixture, while the CRISPR analysis can only replicate a limited amount of effects due to incomplete gene overlap and differences between the cell types. Perhaps more importantly, the number of co-eQTLs for which we can chart “the complete” regulatory picture is limited, as this requires both TF implication and the TF motif being perturbed. However, motifs of TFs are not perfect and are likely still incomplete especially when including co-factors^46^. These co-factors often bind around the binding site of the TF^47^, and so scenarios when the variant is adjacent to the TF binding site may in fact also indicate a disruption of the regulatory mechanism. For this to be reliable we would need to also have a reliable set of co-factor binding sites. There are also limitations on the variant side: 1) we limit our analyses to variants tested within all datasets in the consortium which lowers the number of testable variants; 2) given the current sample sizes we will have imperfect fine-mapping due to discovery power. Both of these factors limit our power to pinpoint the causal variant altering the TF binding.

In summary, we have shown that co-eQTLs provide mechanistic insights into how eQTL variants function and more importantly, how disease-associated variants perturb the wiring of expression regulation. We have also shown the additional disease-relevance of co-eQTLs compared to classical eQTLs. This highlights the need to move from a single gene perspective to a gene network perspective to make progress in understanding how genetic risk variants are implicated in disease. The meta-analysis framework we have established here can form the basis for dissecting this regulatory architecture of disease as it can easily be applied to and combine data from different cohorts in order to identify disease-implicated variants that change gene expression networks at scale. We foresee that this network perspective will, in future, help to better prioritize targets to treat patients and even to prevent disease in individuals at risk.

## Author contributions

- Conceptualisation: D.K., C.L., M.J.B., M.H., L.F.
- Data curation & Analysis: D.K., C.L., M.K., R.O., M.J.B
- Analysis (co-localization & fine-mapping): D.C., Y.T.
- Analysis (replication): P.D., R.W., M.J.B
- Funding acquisition: M.H., L.F.
- Methodology and software: D.K., C.L., R.O., M.V., A.K., H.W., M.W., M.J.B
- Supervision: M.W., M.J.B., M.H., L.F.
- Writing – original draft and Visualisation: D.K., C.L., H.W., M.J.B., M.H., L.F.
- Writing – review & editing: D.K., C.L., M.K., H.W., M.W., M.J.B., M.H., L.F.

## Acknowledgements

We would like to thank all participants of the individual dataset that contributed material to do these studies, and all collaborators within the sc-eQTLGen consortium. We would like to thank Kate Mc Intyre for editorial assistance and feedback on this manuscript. We would like to thank the Center for Information Technology of the University of Groningen for their support and for providing access to the Habrok high-performance computing cluster, as well as the UMCG Genomics Coordination center, the UG Center for Information Technology, and their sponsors BBMRI-NL and TarGet for storage and compute infrastructure.

L.F., D.K., M.J.B. are supported by a grant from the Dutch Research Council (ZonMW-VICI 09150182010019 to L.F. ZonMW LongCOVID grant 10430302110002), European Union’s Horizon Europe Research and Innovation Program grant 101057553 (LongCovid) and through a Senior Investigator Grant from the Oncode Institute and a grant from Saxum Volutum (Pericode). M.H. is supported by the Chan Zuckerberg Foundation (2019-202666, 2021-237882) and the DZHK (German Center for Cardiovascular Research) projects 81Z0600106 and 81Z0600105. M.J.B. is supported by a grant from the Dutch Research Council (ZonMW-VIDI 09150172310068). M.W. is supported by a grant from the Dutch Research Council (Vidi 223.041). P.D. is supported by an NWO ZonMW-VENI Grant (no. 9150161910057) C.L. is supported by the Helmholtz Association under the joint research school ‘Munich School for Data Science – MUDS’.

## Code availability

All code is available on GitHub at:

https://github.com/sc-eQTLgen-consortium/co-eQTL-meta-analysis

## Data availability statement

Our study is based on a collection of previously published single cell eQTL studies. These datasets are available upon request through the original publication.

Suppl. Table 2 can be found on Zenodo at: https://doi.org/10.5281/zenodo.17242541

## Ethics statement

### Ethics approval and informed consent for OneK1K

The OneK1K dataset was generated under the oversight of the Tasmania Health and Medical Human Research Ethics Committee (H0012902) in compliance with local ethical guidelines and with informed consent for all participants, as described in ref. 3. These authors provided OneK1K data access for the current study before publication.

### Ethics approval and informed consent for all other studies

All other studies fall under the METC of the LifeLines DEEP study. The study was approved by the ethics committee of the University Medical Centre Groningen, document number METC UMCG LLDEEP: M12.113965. All participants signed informed consent from prior to study enrolment. All procedures performed in studies involving human participants were in accordance with the ethical standards of the institutional and/or national research committee and with the 1964 Helsinki declaration and its later amendments or comparable ethical standards.

## Methods

### Datasets

Our co-eQTL results are based on data uniformly processed within the single cell eQTLGen consortium. Specifically the following datasets were selected: OneK1K^3^ (n = 1,018), UMCG multiome (*manuscript in preparation*, n = 101), (Oelen^6^ and van Blokland^12^ split) UMCG V2 (n = 123), UMCG V3 (n = 47), van der Wijst^4^ (n = 41).

All datasets were processed through the pipelines of sc-eQTLGen (*manuscript in preparation*) ensuring that they are harmoniously processed to allow for easy integration. In short, the data were taken after alignment to the genome, as performed by the dataset contributors, they were uniformly demultiplexed, a consistent doublet detected method was applied, and empty droplets were removed. After this the cells were cell type annotated using azimuth and scPred, and cell type labels were taken based on the combination of these two tools. Per cell type outlier cells were detected by removing outliers on both the number of UMIs (maximum MAD 2), and the level of MT express (maximum MAD 3). Lastly, the past QC single-cell expression counts were normalized by applying the PFlog1PF^48^ normalization.

### Co-expression calculation

Previous co-eQTL mapping papers^9^ have used Spearman correlation to calculate the co-expression of two genes to get a non-parametric estimation of co-expression. However, multiplexed 10X single-cell data is characterised by having large variation in the amount of cells and reads per individual which affects the reliability of the calculated correlation. As there are also versions of the Spearman and Pearson correlation which take into account weights per observation, allowing to weigh the observations based on their reliability (in our case reads per cell). We evaluated these correlation measures, using simulations to identify the most accurate and robust way to measure co-expression within single-cell data.

We used a well-established Gaussian copula/ NORTA^49,50^ approach to simulate correlated gene pairs that follow a negative binomial distribution which is the distribution usually observed for single-cell RNA-seq count data. We simulated counts to reflect scenarios that we observed in the Van der Wijst^4^ dataset. More information on the concrete specification of the simulation is given in the Supplementary Methods.

After having simulated the count data for each individual of each of the scenarios we calculated the correlations on the simulated count data for each gene pair for each individual with the different evaluated correlation measurements:

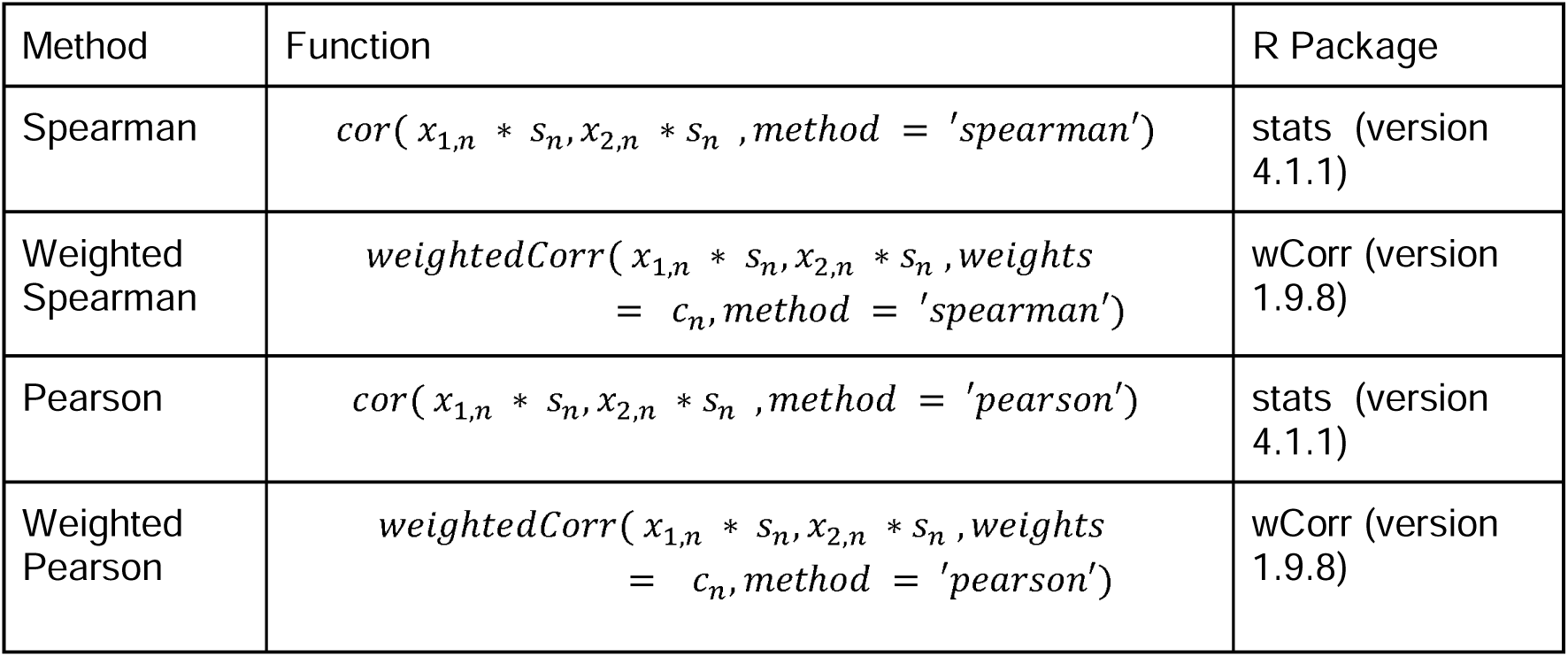

| Method | Function | R Package |
| --- | --- | --- |
| Spearman | $\text{cor}(x_{1,n} * s_n, x_{2,n} * s_n, \text{method} = 'spearman')$ | stats (version 4.1.1) |
| Weighted Spearman | $\text{weightedCorr}(x_{1,n} * s_n, x_{2,n} * s_n, \text{weights} = c_n, \text{method} = 'spearman')$ | wCorr (version 1.9.8) |
| Pearson | $\text{cor}(x_{1,n} * s_n, x_{2,n} * s_n, \text{method} = 'pearson')$ | stats (version 4.1.1) |
| Weighted Pearson | $\text{weightedCorr}(x_{1,n} * s_n, x_{2,n} * s_n, \text{weights} = c_n, \text{method} = 'pearson')$ | wCorr (version 1.9.8) |

For the weighted correlations we use the weighting factor *c_n_* which specifies the number of genes with a read in cell *n*. So observations from cells with more non-zero reads across genes are higher weighted. We then tested each gene pair within each scenario for a co-eQTL effect by specifying a linear model (lm) with the correlation value as y and the genotype encoded as numeric covariate x. We only included a gene pair of an individual in the testing when each of the genes has at least 10 non-zero counts in the simulated data.

To evaluate which of the evaluated correlation measures works best for eventual co-eQTL discovery we then compared two evaluation criteria among the four different methods. First we calculated the mean squared error between the estimated effect size (β) from the linear model and the specified difference in the correlation between the gene pair (ρ): 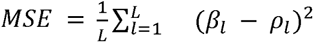. Second we used the Bonferroni adjusted p-value from the linear model to plot ROC curves (ggplot2; version: 3.5.1, geom_roc), taking the adjusted p-value as threshold to specify whether a gene pair is a true co-eQTL or not a co-eQTL based on the specified correlation difference for the simulation.

For both evaluations the weighted Pearson approach showed the best performance (Suppl. Fig. 2; therefore, we used this approach in our co-eQTL mapping.

### co-eQTL mapping

The co-eQTL mapping we performed in two steps: <u>Evaluation</u> and <u>Application</u>. In the first evaluation step we did several analyses on a subset of the data to devise the best strategy to run the co-eQTL mapping. In the second application step we then applied this strategy to run the co-eQTL mapping on all datasets and cell types.

#### Evaluation

##### Gene filtering strategy

First we selected genes for the co-eQTL mapping. We were applying a gene filtering strategy for two reasons:

1. There are very low expressed genes in single-cell data for which it is very unlikely that we robustly measured them to calculate correlations for those.
2. Testing all genes would result in an unreasonable multiple testing burden causing us to miss many true co-eQTLs. Additionally, the computational burden of testing all genes and variants is too large to be completed in a realistic time frame.

For these reasons we limited the number of gene pairs per dataset to no more than 12,000,000, basing our calculations to be reasonable when analysing the largest dataset.

We defined our filtering strategy on the CD4+ T cells from the OneK1K datasets because this was the cell type with most cells in the dataset with most individuals and as a result the setting where we expected most power to detect co-eQTLs. This is important as we expect that if we define filtering criteria in this setting these can also be applied to the lower powered settings of the other datasets and cell types. So we could identify accurate but lenient filtering thresholds.

##### Initial co-eQTL mapping in OneK1K CD4+ T cells

To identify genes for which it is unlikely to detect co-eQTLs we performed a first co-eQTL mapping on CD4+ T cells in OneK1K. As input we used the 7,310 eQTL genes that were identified in the sc-eQTLGen consortium across all cell types. We calculated a weighted Pearson correlation, as described above, for all gene pairs resulting from the combinations of these 7,310 genes for each individual if for both genes there were at least 10 non zero counts across all cells of the individual within CD4+ T cells. Then we normalised the correlations per gene pair using Gaussian inverse rank normalisation. For each gene pair where we calculated a correlation value in at least 20 individuals we then tested all variants that were found as a significant eQTL variant for at least one of the two genes using a linear mixed model (LIMIX QTL^51,52^). This resulted in a total of 31,497,753 tests for all variant-gene pair triplets.

At Bonferroni significance level we identified 2,432 significant co-eQTLs. We then removed all triplets containing ribosomal and mitochondrial genes because we were biologically more interested in finding co-eQTLs for non-ribosomal/mitochondrial genes and wanted to make sure that these were not removed by our filtering strategy. This resulted in a set of 1,029 non-ribosomal/mitochondrial significant co-eQTLs in CD4+ T cells in OneK1K.

##### Gene characteristic calculation

In the second step we defined metrics per gene pair that we hypothesized to be characteristics that influence the possibility to detect a co-eQTL. Specifically we assessed: *Mean absolute correlation, Variance of the correlation, Weighted Variance of the correlation, Number of individuals with correlation values, Percentage of significant correlations, The mean amount of cells with a zero count in at least one gene per gene pair, The mean amount of cells with a zero count in both genes per gene pair, The mean amount of cells with a non-zero count in both genes per gene pair, Minimum mean expression per gene-pair, Minimum mean variance per gene pair, Minimum of the sum of non-zero counts per gene of the gene-pair, Maximum mean percentage of zero per gene pair, which corresponds to calculating the percentage of zeros*. The calculation of these characteristics is given in the Supplementary Methods.

##### Decision tree classification

We then trained a decision tree using the sci-kit learn^53^ package (version 1.3.2) in Python (version 3.9.6) to predict significant co-eQTLs versus non-significant co-eQTLs using these metrics as features to evaluate which metric is the best to separate the significant co-eQTLs from the non-significant co-eQTLs. The list of true co-eQTLs was given as the 1,029 significant co-eQTLs identified when using a Bonferroni multiple testing correction and the remaining 30,326,351 tests as the non-significant co-eQTLs (the effects excluding ribosomal and mitochondrial genes). The data was then randomly split up into 80% for the training set and 20% for the testing set. We then trained the decision tree on the training set with depth 1 while up-weighing significant co-eQTLs with a factor of 250,000 to overcome the huge class imbalance, putting more importance on predicting significant co-eQTLs. The resulting tree led to a recall of the positive cases over 99% in both train and test (suppl. Fig 3), only classifying 4 out of the 1,029 significant co-eQTLs incorrectly. For predicting non co-eQTLs we achieved a recall of 49.96% (Suppl. Fig. 3), suggesting that we can remove about half of our tests and still identify >99% of our co-eQTLs. The feature that was selected by the tree to divide the significant co-eQTLs from the non-significant co-eQTLs was the max_mean_percentage_zero metric at a threshold of 95.89%.

Applying this threshold we could reduce the amount of tests by roughly 50% while keeping nearly all of the identified co-eQTLs at the Bonferroni threshold, allowing us to widen our search space and optimize our scan and final multiple testing burden.

#### Application

After these evaluations our final co-eQTL mapping method that we applied to all datasets and cell types consisted of the following steps:

##### Identification and filtering of gene pairs to test

Starting with the complete set of genes measured in each dataset we kept only genes that fulfilled the following criteria:

- The mean percentage of zero across all individuals of the gene was lower than 95.891 (which corresponds to the optimal threshold for filtering that was identified in the decision tree analysis)
- The gene has a measured expression value that is not zero in at least 10 cells for a given individual for which a correlation is calculated.
- The gene fulfilled the criteria above in at least 20 individuals, so that correlation values could be calculated for at least 20 individuals.

Two different lists of genes were created, one for potential eGenes and another for potential co-eGenes. The difference between these lists was that for potential co-eQTL eGenes only genes were selected that had a significant eQTL within this cell type in the sc-eQTLGen results. Further the set of genes fulfilling these criteria were evaluated separately per dataset and cell type resulting in an individual gene list per cell type and dataset. The final list of gene pairs for co-eQTL testing is then formed by combining all possible eGenes with all possible co-eGenes.

##### Calculation of weighted Pearson correlation and normalization

For each gene pair, outlined above, and for each individual we then calculated a weighted Pearson correlation across the cells of the individual, weighing for the number of genes that are expressed in each cell.

##### Identification of variants to test

In the next step we defined the set of variants to test for each gene pair. As each gene pair consisted of at least one eGene we leveraged the fine-mapping information from sc-eQTLGen to specifically test the most significant variant per credible set per gen. In case no credible set could be mapped for an eGene the variant with the highest effect size was tested.

Besides that, as we were specifically interested in how eQTLs and co-eQTLs are involved in diseases, we added additional GWAS variants to the testing. For this we downloaded all the GWAS variants from the GWAS catalog (EBI GWAS catalog version 1.0.2 with all associations) and kept only the genome wide significant variants (p < 5*10^8^). We then filtered these GWAS to those that could be assigned to “immune system diseases” (EFO term: EFO:0000540) as we were studying immune cells from blood and therefore mostly expected to see co-eQTLs linked to those diseases. We then assessed if these GWAS variants overlapped the *cis* window of a selected eGene (<=1MB from the eGene), and if the variant was within this distance added it as an additional test for the gene pair for that specific eGene.

##### co-eQTL mapping and meta-analysis

The co-eQTL mapping was carried out separately per dataset and cell type. The input for the co-eQTL mapping consisted of the weighted Pearson correlations and the imputed genotype dosages that were part of the variant sets outlined above. The gene pair correlations were normalised by applying Gaussian inverse rank normalisation before testing associations to genetics. We used the linear model implemented in LIMIX QTL^51,52^ for this. To combine the information over the datasets we leveraged a weighted z-score meta-analysis^45,54^. We weighted the co-eQTL gene pair summary statistics for the number of individuals for which we have an observation per dataset. This is done separately per cell type.

##### Multiple testing correction to identify significant co-eQTLs

Multiple testing correction was carried out separately per cell type and for the internal replication also per dataset. Before multiple testing correction per dataset we filtered out all tests that were based on less than 10% of the individuals of the dataset (OneK1K: 102; multiome: 12; UMCG_v3: 4; UMCG_v2: 10; van der Wijst: 4). For the meta-analysis the results needed to be testable in 10% of the total individuals (n = 133).

We then applied a two-step multiple testing approach. In the first step we calculated the number of tests per eGene and corrected the p-values per eGene using the Bonferroni correction, correcting for all tests of per eGene (SNPs and co-eGenes).

In the second step we selected the top effect per eGene (given by the minimum p-value across all the tests of the eGene) and corrected the adjusted p-values of the top effects across all eGenes using the q-value approach (q-value^55^; R package, version: 2.26.0). We deemed all effects with a q-value of 0.05 as significant. Lastly we went back to BF corrected p-values and determined for all tests for which the q-value corrected p-value was lower than 0.05 and took these forward as our significant effects.

##### Full window scan followed by fine-mapping and colocalization

After the discovery of the significant co-eQTLs from the main co-eQTL scan, we performed a focused full window scan for the significant co-eQTL pairs. Here we assessed the effects of all variants in a 1Mb window around each significant co-eQTL eGene. On the full summary statistics per dataset we once again applied a meta-analysis, as described above.

These summary statistics were the input for fine-mapping using Susie^13^ (susieR v = 0.12.35). We adapted a “joint” fine-mapping approach, where we first fine-mapped all co-eQTL gene pairs separately, followed by a colocalization, using coloc^56^, step to identify the shared causal variants over the co-eGenes per eGene. The colocalization was performed using the coloc package based on the log bayes factor (LBF) information from susie (leveraging the coloc function: “coloc.bf_bf”). When the gene-pairs were colocalizing we updated the LBF information to the information from the credible set that had the strongest support (highest LBF). By doing so we identified independent causal variants over eGenes (co-eQTL credible sets).

We leveraged two cut-offs for determining colocalization between co-eQTL gene pairs for different downstream analysis. For initial colocalization with gene-pairs of the same eGene as well as for the colocalization to GWAS we leveraged a conservative level where only colocalizations were taken as true when the sum of H3 and H4 was higher than 0.9, and H4/H3 was higher than 3. When we attempted to group co-eQTLs per variant to get a conservative estimate of the number of independent effects per eGene we used a less strict threshold of H3+H4 > 0.5, and H4/H3 > 1.

#### co-eQTL Validation

##### Replication in Bulk

To validate our co-eQTLs we leveraged bulk whole blood data generated by the BIOS consortium^10^. Specifically we had access to data from four datasets, LifeLines-Deep^57^, Leiden Longevity study^58^, Netherlands twin registry^59^, and the Rotterdam study^60^. The combined dataset totalled 2,402 samples. In each of the dataset we inverse rank transformed the sample expression levels and corrected the expression levels for 25 PCs derived from the expression data, as well as factor describing the sample correlation to mean gene expression levels. On the normalized data we fitted per dataset a linear model explaining expression of the eGene by the ‘variant’, the ‘co-eGene expression’ and ‘variant * co-eGene expression’. The last term, the interaction between variant and co-eGene was used for the replication of the co-eQTL effects. The per dataset interaction summary statistics were meta-analyzed by a weighted meta-analysis, weighting a dataset for the number of individuals it comprised. The meta-analyzed data was used to replicate the co-eQTL effects per cell type.

##### Annotating cell count driven effects

In order to evaluate the impact of cell subtype compositions we used two different analyses; an interaction co-eQTL test and we leveraged information from the cell type composition GWAS performed in sc-eQTLGen. For the interaction test we expanded the linear model used for co-eQTL mapping and added two terms: 1) the fractional abundance of the largest cell subtype count relative to the cell type of interest, and 2) added the interaction between the co-eQTL eVariant and the cell count fraction. We tested all significant co-eQTL effects using this interaction model within the largest single dataset (OneK1K) for effects due to cell subtype composition. When the interaction term in the model was significant after correcting for multiple testing, using the Benjamini Hochberg method, over the significant effects per cell type the effects were marked as likely cell type composition driven effects. Likewise, if the variant was linked to a cell count composition trait in the ccQTL mapping, for the relevant cell type, we annotated these effects as likely cell count composition effects.

##### Evaluation of eQTL effect with simulations

Throughout our evaluations of different correlation measures and factors influencing the correlation measurement in scRNA-seq data we observed that the expression of a gene does have a strong influence on the measured correlations. Lower expression of a gene implied larger variance of correlation measurements as well as lower average correlation measurements. This could influence co-eQTL detection as one of the genes of our gene pairs is defined as eGene whose expression varies depending on the associated eQTL variant. To determine the effect of those expression differences on our identified co-eQTLs we performed simulations for each significant co-eQTL effect where we artificially defined the same correlation for all genotypes and only varied the expression levels of the single genes according to their expression differences in the real data. We then evaluated whether those expression differences alone could lead to detecting co-eQTL effects. We used the same simulation approach as specified above, conducted 50 independent simulations for each significant co-eQTL and specified the parameters in each iteration as given below:

- number of simulated individuals: for each genotype the number of individuals as in the OneK1K data for this genotype was simulated
- number of cells: a distinct number of cells was simulated for each individual as function of max(round(rnorm(1, mean = ncells_mean_oneK1K, sd = ncells_sd_oneK1K)),10), with ncells_mean_OneK1K = the mean number of cells across all the individuals in the co-eQTL cell type in OneK1K and ncells_sd_oneK1K = the standard deviation of the number of cells across all the individuals in the co-eQTL cell type in OneK1K
- true correlation (ρ) between the gene pair: First the mean correlation of all individuals of a genotype in OneK1K for this gene pair was calculated. Then the absolute maximum of those mean correlations was set as true correlation between the gene pair for all simulations.
- mean (*µ*_1_, *µ*_2_) and dispersion parameters (*θ*_1_, *θ*_2_) of gene1 and gene2 were derived from the real data (OneK1K) estimating a negative binomial distribution on the counts (fitdistr, R)

We then computed weighted Pearson correlation for each simulated individual and each iteration and performed the co-eQTL mapping as described above with a linear model. This was followed by comparing the p-values of each of the 50 iterations of this simulated co-eQTL mapping to the p-values of the actual co-eQTL mapping. To indicate the influence of the expression on the co-eQTL mapping we added a score to the final co-eQTL results table (Suppl. Table 2) which indicates in how many of the 50 iterations in this simulated co-eQTL mapping we would get a p-value equal or lower than observed in the actual co-eQTL mapping (column: simulation_score).

#### co-eQTL interpretation and downstream analysis

##### GWAS colocalization

Fine-mapped credible sets from single-cell co-expression QTLs, were converted from susie output into StudyLocus format from gentropy (https://github.com/opentargets/gentropy; version 2.0) to be consistent with the GWAS credible data format in Open Targets. These credible sets were then overlapped with all GWAS credible sets in the Open Targets ecosystem (March 2025 data release), an overlap being defined as when two credible sets (GWAS vs eQTL/co-eQTL) share at least one variant. The LBFs for each variant were then used to calculate coloc. posterior probabilities, where we infer an LBF of 0 for variants which don’t exist in one of the two credible sets of an overlap. All analyses were performed on Google Cloud Platform via Airflow using our pipeline orchestration. The maths in the coloc. implementation is the same as the original coloc. R package^56^, and is previously benchmarked against that to ensure it produces the same output. Credible sets where filtered such that the minimal LBF was 2, or >= 0.8686 in log10BF scale, and the credible set had at least one association at p-value <= 5.0*10^−3^.

##### GWAS enrichment

We conducted two analyses to test for enrichment of GWAS effects among our co-eQTLs; one variant based and one eGene based. To assess the relation between the enrichments with different categories we also stratified these enrichments across several GWAS subsets (e.g. removing cell-count related traits). To do so we mapped the GWAS traits to EFO-Ids^61^ and corresponding parent terms from the GWAS catalog^1^ using gwasrapidd; version: 0.99.17. The different categories were defined as given below:

- All traits: no filtering; all traits in GWAS catalog
- All traits (removed cell-count traits): removed all traits assigned to EFO parent term ‘Hematological measurement’ (EFO_0004503) and these additional traits: ‘basophil count’, ‘basophil percentage of leukocytes’, ‘basophil percentage of granulocytes’
- Disease GWAS: only kept traits assigned to these EFO parent terms: ‘Cancer (EFO_0000616)’, ‘Immune system disorder (EFO_0000540)’, ‘Other disease (EFO_0000408)’, ‘Cardiovascular disease (EFO_0000319)’, ‘Neorological disorder (EFO_0000618)’, ‘Metabolic disorder (‘EFO_0000589’), ‘Digestive system disorder (EFO_0000405)’
- Immune disease GWAS: only kept traits assigned to these EFO parent terms:’Immune system disorder (EFO_0000540)’

To analyze the enrichment across all cell types (‘All’ category) we used only one cell type observation per eGene. If an eGene had an eQTL effect in multiple cell types the observation of the cell type with the lowest p-value of the eQTL effect was used and also the corresponding colocalization and co-eQTL classification within this cell type.

##### Variant based GWAS enrichment

To conduct the variant based GWAS enrichment, we took the fine-mapped eQTL variants from sc-eQTLGen (manuscript in preparation) that do show a co-eQTL effect (before full window scans) and those that do not. We annotated these variants, taking into account LD, with GWAS genetic variants with associations (v1.0.2) downloaded from the GWAS Catalog, these variants have been filtered to contain variants that are genome-wide significant (P < 5*10^−8^), and we used EFO terms^61^ to group variants into 4 different trait groups (All traits, not cell count traits, disease associated traits, immune-related traits).

LD information was used to filter the eQTL and co-eQTL sets (R^2^ >= 0.1) as well as determining whether the GWAS genetic variants are in LD (R^2^ >= 0.8) to the eQTL and co-eQTL sets. OneK1K^3^ was used as our LD reference dataset for this purpose, the pre-processing of which was done in sc-eQTLGen (manuscript in preparation). The eQTL and co-eQTL sets of variants were filtered by keeping the most significant of the variants that are in LD. eQTL and co-eQTL variants are determined to be GWAS variants either if they are present in the GWAS catalog\ set of GWAS variants or if they are in LD with any of the GWAS variants.

##### eGene based GWAS enrichment

As a secondary measure of GWAS enrichment we evaluated whether eGenes with significant co-eQTL effect were more often associated with GWAS traits than eGenes without significant co-eQTL effect. To do so we assess the relation of a gene being a co-eQTL implicated eGene and if the eGenes is significantly colocalizing to disease, by leveraging a Fisher’s Exact Test (fisher.test in R).

As throughout our analyses we observed that having a credible set, i.e. if the eQTL was fine-mapped, for an eGene was not independent of an eGene having a co-eQTL effect (Suppl. Fig. 15) we ran the enrichment analysis in two different settings. In both of these settings the set of eGenes considered for the enrichment analysis was the total number of eGenes per cell type with eQTL effects from sc-eQTLGen which passed the decision tree filter and had been used as input for the co-eQTL mapping, HLA eGenes were removed from the enrichment analysis. The settings differed by whether eGenes without mapped credible sets were included in the analysis (setting 1; Suppl. Fig. 11, Suppl. Tables 7,9) or excluded from the analysis (setting 2; Suppl. Fig. 11, Suppl. Tables 7,9). Co-eQTL effects that have only been identified by the GWAS variants, that we included for disease interpretation, were moved to the non-co-eQTL background so that an eGene was only considered to have a significant co-eQTL effect when the effect was detected based on an eQTL effect variant.

##### eGene Mostafavi analysis

To evaluate whether eQTLs with co-eQTL effect compared to eQTLs without co-eQTL effect have characteristics more similar to GWAS hits as reported previously by Mostafavi *et al*^8^, we downloaded the gene and variant annotations as provided in the publication^62^. The set of eGenes and variants considered for this analysis consisted of all eGenes and mapped variants that passed the decision tree filter and were used as input for the co-eQTL mapping. To those eGenes and variants we mapped the gene and variant annotations from the Mostafavi *et. al.* publication and classified the eGenes and variants as having a significant co-eQTL effect or not using the meta-analyzed significant results from our co-eQTL mapping.

##### TSS count comparison

To compare the TSS count as presented in the Mostafavi *et. al.* publication we used the promoter_count feature (TSS count across biosamples, processed using data from the FANTOM project^17^) from the publication and performed a t-test comparing eGenes with and without co-eQTL effect.

For the comparison across all cell types (‘All’) we considered an eGene to have a significant co-eQTL effect if it had a significant co-eQTL effect in any cell type.

##### Enhancer domain landscape comparison

To compare the enhancer domain landscape in the same way as in the Mostafavi *et. al.* publication we set up two logistic regression models predicting either variants with co-eQTL effect (y=1) or variants without co-eQTL effect (y=1) versus 100,000 variants chosen at random (y=0) from the full set of annotated variants from the Mostafavi *et. al.* publication. As in the publication we included the following covariates in the regression to account for potential confounders:

- MAF: minor allele frequency
- L2: LD score
- TSSD: gene density (defined as the number of TSSs within 1Mb window around a gene’s TSS)
- d_tss: absolute distance to closest protein-coding gene (with closest TSS)
- length: total gene length ((in Kb), |TSS - TES|)
- CDS_length: total length of gene coding sequence (in Kb)
- Roadmap_length_per_type: mean length of Roadmap Epigenomics Consortium enhancers across active biosamples, (processed using data from Liu *et al*^18^, Genome Biology 2017)
- Roadmap_count: counts of Roadmap Epigenomics Consortium^18^ enhancers across biosamples

For the comparison across all cell types (‘All’) we considered a variant to have a significant co-eQTL if it had a significant co-eQTL effect in at least cell type.

##### Credible set comparison

To evaluate which other characteristics besides the findings of the Mostafavi *et al* paper based analysis characterized eGenes with co-eQTL effect compared to eGenes without co-eQTL effect we estimated a logistic regression model (glm) to predict whether an eGene had a co-eQTL effect (1) or not (0). As covariates we used:

- Number of credible sets (SusieRss_CS): the maximum number of credible sets for the eGene resulting from the finemapping of the eQTL effect from sc-eQTLGen (eGenes without mapped credible set were excluded)
- Absolute eQTL effect size (beta): the absolute value of the beta from the eQTL-effect given from sc-eQTLGen
- Weighted mean expression: given by the mean of the mean expression of the eGene per individual in each dataset weighted by the amount of samples of the datasets
- Mapped gene-based characteristics from the Mostafavi et. al. publication also used in the Mostafavi analysis: total enhancer length per tissue/ cell type (Roadmap_length_per_type), TSS count across biosamples (promoter_count), gene density (TSSD), gene length(kb), length of the coding segments (CDS_length), count of active tissue/ cell types (Roadmap_count)

All independent features were standardized (z-score normalized) to have a mean of 0 and a standard deviation of 1.

The set of eGenes considered for this analysis consisted of all eGenes that passed the decision tree filter and were used as input for the co-eQTL mapping. Per eGene we used the eQTL effect (variant + eGene) with the lowest p-value. eGenes were classified as having a significant co-eQTL effect or not using the meta-analyzed significant results from our co-eQTL mapping.

To set up the model for across all cell types (‘All’) we used only one cell type observation per eGene. If an eGene had an eQTL effect in multiple cell types the observation of the cell type with the lowest p-value of the eQTL effect and the corresponding amount of maximum credible sets in this cell type was used.

Additionally, we estimated the same model dividing the set of eGenes into eGenes colocalizing with GWAS traits or not (strict threshold) (Suppl. Fig. 14) to evaluate the influence of those features onto GWAS colocalization. As for the GWAS enrichment analysis we ran the model filtering the GWAS traits to different categories using EFO terms.

##### co-eGene comparisons with Mostafavi features

Similarly, to identify features important for the co-eGenes contributing to co-eQTL pairs we again used gene annotations from Mostafavi *et al*. The set of genes for this analysis were all genes that passed the decision tree filters and were tested as co-eGenes in the co-eQTL mapping. As gene expression is highly influencing the capability to detect co-eQTLs we first corrected for gene expression by estimating a linear model (lm) using the weighted mean expression of a gene across datasets as covariate and predicting whether a tested gene was a co-eGene (y = 1) or not (y = 0). The residuals of this model were then used as independent variables in a second linear model to evaluate the association of other covariates with a gene being a significant co-eGene or not. We tested the connectedness (connectedness in co-expression networks inferred by Saha *et al*, connectedness score computed using InWeb PPI network), GO term (count of BP broadly unrelated GO terms per gene) and LOEUF score features extracted from Mostafavi *et al*.

##### CRISPRi validation

We used a publicly available perturb-seq dataset from Replogle *et al*^26^ from which we used the processed table that only includes perturbations where the target gene expression was reduced by at least 30%, there were at least 50 significant variable genes per target gene and at least 25 cells pass with the target perturbed pass QC. We overlapped this with our co-eQTLs, where we took the co-eGene to be the perturbed gene and looked up the observed effect on the eGene that corresponded to the co-eQTL gene pair. To see whether these perturbations were meaningful we then compared this to another set of genes, where the perturbed gene is the same co-eGene, but the other gene is a randomly selected eGene that has been shown to have a co-eQTL effect but not with this particular co-eGene. To determine whether this difference was significant we did a Wilcoxon signed-rank test, and to account for the number of random genes that could have been chosen we calculated 100 Wilcoxon p-values with different random eGenes each time and selected the largest p-value of these 100.

For calculating concordance between the co-eQTL perturbation results we used the average correlation of the gene pair as the co-eQTL direction and then took the opposite direction of the perturbation as the direction of effect of the perturbed gene. This is because after a perturbation, if the expression of a gene increases the perturbed gene was having an inhibitory effect.

##### MotifBreakR analysis

We ran MotifBreakR (motifbreakR^27^; v = 2.8.0) on the initial set of variants tested in the co-eQTL mapping and showing a significant co-eQTL effect, and all variants assigned to a credible set after the co-eQTL finemapping. We used genome build HG38 and a p-value threshold of p < 1*10^−4^ to determine motifbreakR hits. Databases checked were: HOCOMOCOv11-core-A, hPDI, jolma2013, stamlab, SwissRegulon, JASPAR_CORE, JASPAR_2014, jaspar2016, jaspar2018, cisbp_1.02, HOMER, HOCOMOCOv10. MotifbreakR hits were then overlapped with transcription factor annotations among the co-eGenes using transcription factors, defined as genes in the GO Term GO0050789.

##### co-eGene enrichment analyses

To get insight into the biological processes, transcription factors, microRNAs and other factors the co-eGenes per eGene reflect, we performed geneset enrichments using g:profiler^63^. We leveraged the R package to systematically test the groups of significant co-eGenes against all tested genes. Specifically, we ran the enrichments both per variant, testing all co-eGenes per initial variant (from the initial meta-analysis) and over all variants per eGene jointly. Only tests where more than 10 co-eGenes were identified per co-eQTL, or co-eQTL variant pair, were performed. All settings were left at defaults provided by the tool developers, other than the background sets. We set the background per eGene, where we used all tested co-eGenes as background.

Additionally, to get higher resolution on the relevance of TFs we prepared specific test information by parsing the information from the remap project. Specifically, we added three extra information sources, we added the *cis* regulatory modules (CRMs) as defined by remap, the non-redundant TF information over all cell types and the non-redundant TF information for blood relevant cell types (more details in the previous co-eQTL paper). These information sources were added by providing custom data to g:profiler.

## References

1. Cerezo, M. et al. The NHGRI-EBI GWAS Catalog: standards for reusability, sustainability and diversity. Nucleic Acids Res. 53, D998–D1005 (2025).

2. Fabo, T. & Khavari, P. Functional characterization of human genomic variation linked to polygenic diseases. Trends Genet. TIG 39, 462–490 (2023).

3. Yazar, S. et al. Single-cell eQTL mapping identifies cell type-specific genetic control of autoimmune disease. Science 376, eabf3041 (2022).

4. van der Wijst, M. G. P. et al. Single-cell RNA sequencing identifies celltype-specific cis-eQTLs and co-expression QTLs. Nat. Genet. 50, 493–497 (2018).

5. Kock, K. H. et al. Asian diversity in human immune cells. Cell 188, 2288–2306.e24 (2025).

6. Oelen, R. et al. Single-cell RNA-sequencing of peripheral blood mononuclear cells reveals widespread, context-specific gene expression regulation upon pathogenic exposure. Nat. Commun. 13, 3267 (2022).

7. van der Wijst, M. et al. The single-cell eQTLGen consortium. eLife 9, e52155 (2020).

8. Mostafavi, H., Spence, J. P., Naqvi, S. & Pritchard, J. K. Systematic differences in discovery of genetic effects on gene expression and complex traits. Nat. Genet. 55, 1866–1875 (2023).

9. Li, S. et al. Identification of genetic variants that impact gene co-expression relationships using large-scale single-cell data. Genome Biol. 24, 80 (2023).

10. Zhernakova, D. V. et al. Identification of context-dependent expression quantitative trait loci in whole blood. Nat. Genet. 49, 139–145 (2017).

11. Westra, H.-J. et al. Cell Specific eQTL Analysis without Sorting Cells. PLOS Genet. 11, e1005223 (2015).

12. van Blokland, I. V. et al. Single-Cell Dissection of the Immune Response After Acute Myocardial Infarction. Circ. Genomic Precis. Med. 17, e004374 (2024).

13. Wang, G., Sarkar, A., Carbonetto, P. & Stephens, M. A simple new approach to variable selection in regression, with application to genetic fine-mapping. 501114 Preprint at 10.1101/501114 (2020).

14. Kim, M. C. et al. Method of moments framework for differential expression analysis of single-cell RNA sequencing data. Cell 187, 6393–6410.e16 (2024).

15. Flynn, E. D. et al. Transcription factor regulation of eQTL activity across individuals and tissues. PLOS Genet. 18, e1009719 (2022).

16. Mudappathi, R. et al. reg-eQTL: Integrating transcription factor effects to unveil regulatory variants. Am. J. Hum. Genet. 112, 659–674 (2025).

17. FANTOM Consortium and the RIKEN PMI and CLST (DGT) et al. A promoter-level mammalian expression atlas. Nature 507, 462–470 (2014).

18. Liu, Y., Sarkar, A., Kheradpour, P., Ernst, J. & Kellis, M. Evidence of reduced recombination rate in human regulatory domains. Genome Biol. 18, 193 (2017).

19. Pickrell, J. K. Joint analysis of functional genomic data and genome-wide association studies of 18 human traits. Am. J. Hum. Genet. 94, 559–573 (2014).

20. Saha, A. et al. Co-expression networks reveal the tissue-specific regulation of transcription and splicing. Genome Res. 27, 1843–1858 (2017).

21. Pintacuda, G. et al. Genoppi is an open-source software for robust and standardized integration of proteomic and genetic data. Nat. Commun. 12, 2580 (2021).

22. Iacono, G., Massoni-Badosa, R. & Heyn, H. Single-cell transcriptomics unveils gene regulatory network plasticity. Genome Biol. 20, 110 (2019).

23. Karczewski, K. J. et al. The mutational constraint spectrum quantified from variation in 141,456 humans. Nature 581, 434–443 (2020).

24. Ashburner, M. et al. Gene Ontology: tool for the unification of biology. Nat. Genet. 25, 25–29 (2000).

25. The Gene Ontology Consortium et al. The Gene Ontology knowledgebase in 2023. Genetics 224, iyad031 (2023).

26. Replogle, J. M. et al. Mapping information-rich genotype-phenotype landscapes with genome-scale Perturb-seq. Cell 185, 2559–2575.e28 (2022).

27. Coetzee, S. G., Coetzee, G. A. & Hazelett, D. J. motifbreakR: an R/Bioconductor package for predicting variant effects at transcription factor binding sites. Bioinformatics 31, 3847–3849 (2015).

28. Wang, W. et al. The Interaction between Lymphoid Tissue Inducer-Like Cells and T Cells in the Mesenteric Lymph Node Restrains Intestinal Humoral Immunity. Cell Rep. 32, 107936 (2020).

29. Guo, X. et al. Innate Lymphoid Cells Control Early Colonization Resistance against Intestinal Pathogens through ID2-Dependent Regulation of the Microbiota. Immunity 42, 731–743 (2015).

30. Paternoster, L. et al. Multi-ancestry genome-wide association study of 21,000 cases and 95,000 controls identifies new risk loci for atopic dermatitis. Nat. Genet. 47, 1449–1456 (2015).

31. Tanaka, N. et al. Eight novel susceptibility loci and putative causal variants in atopic dermatitis. J. Allergy Clin. Immunol. 148, 1293–1306 (2021).

32. Budu-Aggrey, A. et al. European and multi-ancestry genome-wide association meta-analysis of atopic dermatitis highlights importance of systemic immune regulation. Nat. Commun. 14, 6172 (2023).

33. Miyazaki, M. et al. Id2 and Id3 maintain the regulatory T cell pool to suppress inflammatory disease. Nat. Immunol. 15, 767–776 (2014).

34. Li, Y., Li, D. & Cheng, X. The association between expression of lncRNAs in patients with GDM. Endocr. Connect. 10, 1080–1090 (2021).

35. Mir, M. M. et al. The Role of Pro-Inflammatory Chemokines CCL-1, 2, 4, and 5 in the Etiopathogenesis of Type 2 Diabetes Mellitus in Subjects from the Asir Region of Saudi Arabia: Correlation with Different Degrees of Obesity. J. Pers. Med. 14, 743 (2024).

36. Zhao, X., Shan, Q. & Xue, H.-H. TCF1 in T cell immunity: a broadened frontier. Nat. Rev. Immunol. 22, 147–157 (2022).

37. McEvoy, C. et al. NR4A Receptors Differentially Regulate NF-κB Signaling in Myeloid Cells. Front. Immunol. 8, 7 (2017).

38. Boulet, S. et al. The orphan nuclear receptor NR4A3 controls the differentiation of monocyte-derived dendritic cells following microbial stimulation. Proc. Natl. Acad. Sci. U. S. A. 116, 15150–15159 (2019).

39. Odagiu, L., May, J., Boulet, S., Baldwin, T. A. & Labrecque, N. Role of the Orphan Nuclear Receptor NR4A Family in T-Cell Biology. Front. Endocrinol. 11, 624122 (2021).

40. Odagiu, L. et al. Early programming of CD8+ T cell response by the orphan nuclear receptor NR4A3. Proc. Natl. Acad. Sci. U. S. A. 117, 24392–24402 (2020).

41. Onengut-Gumuscu, S. et al. Fine mapping of type 1 diabetes susceptibility loci and evidence for colocalization of causal variants with lymphoid gene enhancers. Nat. Genet. 47, 381–386 (2015).

42. Crouch, D. J. M. et al. Bayesian Effect Size Ranking to Prioritise Genetic Risk Variants in Common Diseases for Follow-Up Studies. Genet. Epidemiol. 49, e22608 (2025).

43. Robertson, C. C. et al. Fine-mapping, trans-ancestral and genomic analyses identify causal variants, cells, genes and drug targets for type 1 diabetes. Nat. Genet. 53, 962– 971 (2021).

44. Rumker, L. et al. Identifying genetic variants that influence the abundance of cell states in single-cell data. Nat. Genet. 56, 2068–2077 (2024).

45. Võsa, U. et al. Large-scale cis- and trans-eQTL analyses identify thousands of genetic loci and polygenic scores that regulate blood gene expression. Nat. Genet. 53, 1300– 1310 (2021).

46. Kribelbauer-Swietek, J. F. et al. Context transcription factors establish cooperative environments and mediate enhancer communication. Nat. Genet. 56, 2199–2212 (2024).

47. Gate, R. E. et al. Genetic determinants of co-accessible chromatin regions in activated T cells across humans. Nat. Genet. 50, 1140–1150 (2018).

48. Booeshaghi, A. S., Hallgrímsdóttir, I. B., Gálvez-Merchán, Á. & Pachter, L. Depth normalization for single-cell genomics count data. Preprint at 10.1101/2022.05.06.490859 (2022).

49. Knudson, A. D., Kozubowski, T. J., Panorska, A. K. & Schissler, A. G. A flexible multivariate model for high-dimensional correlated count data. J. Stat. Distrib. Appl. 8, 6 (2021).

50. Chen, H. Initialization for NORTA: Generation of Random Vectors with Specified Marginals and Correlations. *Inf*. J. Comput. 13, 312–331 (2001).

51. Cuomo, A. S. E. et al. Optimizing expression quantitative trait locus mapping workflows for single-cell studies. Genome Biol. 22, 188 (2021).

52. Casale, F. P., Rakitsch, B., Lippert, C. & Stegle, O. Efficient set tests for the genetic analysis of correlated traits. Nat. Methods 12, 755–758 (2015).

53. Pedregosa, F. et al. Scikit-learn: Machine Learning in Python. J. Mach. Learn. Res. 12, 2825–2830 (2011).

54. Zaykin, D. V. Optimally weighted Z-test is a powerful method for combining probabilities in meta-analysis: Optimally weighted Z-test is a powerful method. J. Evol. Biol. 24, 1836–1841 (2011).

55. Storey, J. D. & Tibshirani, R. Statistical significance for genomewide studies. Proc. Natl. Acad. Sci. 100, 9440–9445 (2003).

56. Pullin, J. M. & Wallace, C. Variant-specific priors clarify colocalisation analysis. PLOS Genet. 21, e1011697 (2025).

57. Tigchelaar, E. F. et al. Cohort profile: LifeLines DEEP, a prospective, general population cohort study in the northern Netherlands: study design and baseline characteristics. BMJ Open 5, e006772 (2015).

58. Schoenmaker, M. et al. Evidence of genetic enrichment for exceptional survival using a family approach: the Leiden Longevity Study. Eur. J. Hum. Genet. 14, 79–84 (2006).

59. Willemsen, G. et al. The Adult Netherlands Twin Register: Twenty-Five Years of Survey and Biological Data Collection. Twin Res. Hum. Genet. 16, 271–281 (2013).

60. Hofman, A. et al. The Rotterdam Study: 2014 objectives and design update. Eur. J. Epidemiol. 28, 889–926 (2013).

61. Malone, J. et al. Modeling sample variables with an Experimental Factor Ontology. Bioinformatics 26, 1112–1118 (2010).

62. Hakhamanesh Mostafavi. Supplementary Data for ‘Systematic differences in discovery of genetic effects on gene expression and complex traits’. Zenodo 10.5281/ZENODO.6618073 (2023).

63. Reimand, J., Kull, M., Peterson, H., Hansen, J. & Vilo, J. g:Profiler--a web-based toolset for functional profiling of gene lists from large-scale experiments. Nucleic Acids Res. 35, W193–200 (2007).

