## Supplementary material for "Disease-associated variants are enriched for altering cell-type-specific gene co-expression relationships"

### Supplementary Notes

#### Supplementary Results

##### CRISPRi Validation

We used the genome-scale Perturb-seq CRISPRi dataset from Replogle et al.  to compare the directions of effect and the perturbation strength of co-eQTL effects. After overlapping our co-eQTL pairs with the Perturb-seq dataset, we were left with 625 co-eGenes (16% of all co-eGenes) for which we could assess the concordance and perturbation impact on other genes. When comparing the direction of the change in expression of the CRISPRi experiment with the predicted relationship of the two genes based on the co-eQTL correlations we observed that the directional concordance is not higher than expected by chance (48% of effects are concordant, Suppl. Fig. X). The lack of concordance could be due to an overrepresentation of indirect effects among the eGenes, or a gene dosage effect. To test the perturbation strength, for each perturbation of a co-eGene, we compared the perturbation impact between two equally-sized sets of eGenes: those that were and were not part of a specific co-eQTL pair (methods). This showed that eGenes belonging to a co-eQTL gene pair are more impacted by perturbation of the co-eGene compared to gene pairs that did not show a co-eQTL effect (Wilcoxon signed-rank P=8.54x10-7, Fig. 4G). While the concordance is low, when considering the significant effect we see when assessing co-eQTL gene pairs, there does appear to be a biological relevance to these pairs. This indicates that we are often identifying relevant regulator-gene pairs.

##### Replication of Results

We replicated our results in two independent ways, by comparing our co-eQTLs to interaction eQTL results in bulk data (BIOS) and by overlapping effects to a genome-wide CRISPRi Perturb-seq study conducted in K562, a cell type derived from the same tissue (Replogle et al. 2022). In both cases we saw validation of our co-eQTLs through significant replication levels, however these replications showed some counter evidence as well (70% of the significant interaction eQTLs, and 48% of the significant CRISPRi pairs were concordant). Both of these independent sources had drawbacks due to the contexts not being aligned to the co-eQTL settings. Specifically, the interaction analysis in whole blood suffers from cell type composition effects. Most effects are not shared across cell types, given the cell mixture in blood we don't expect all effects to be visible in whole blood. On the other hand the CRISPRi replication is limited due to the limited overlap. Additionally we don't expect all effects to replicate either, as our co-eQTLs are likely a mixture of direct and indirect effects. The indirect effects, which we link to a TF via enrichment, are likely not replicable using either method, but especially for the CRISPRi method this is not possible. Moreover, the gene knockdown induced by CRISPRi perturbations is often much larger than a single genetic variant can induce. Therefore, this difference in effect size may also explain some of the discrepancies.

For further validation we also attempted to falsify our co-eQTLs where we found that some effects were potentially driven by cell sub type composition. However, this analysis indicated that the majority of our co-eQTLs are likely not due to subcell type effects, with the majority of subcell type effects being identified in CD8+ T cells constituting 7.5% of the total effects found in CD8+ T cells.

### Supplementary Discussion

We observed several credible sets that did not colocalize between eQTLs and co-eQTLs. While the strongest effects did colocalize, the lack of overlap in other cases may be due to several factors. These include technical factors such as some of the included GWAS derived disease-associated variants just not being tested in the original eQTL scans, and the noisyness of both the eQTL and co-eQTL association signals combined with the fine-mapping procedure itself. Another possible explanation is to do with biological factors which may explain some observations. Even modest regulatory shifts can have biologically relevant downstream consequences when they occur in the context of interconnected gene networks. This suggests that a variant might only weakly affect the expression of a single gene (i.e., a weak or undetectable eQTL), and if that gene is part of a co-expression network (i.e., it's highly connected to other genes), this seemingly minor regulatory effect could still be biologically meaningful. In other words, a small shift in one gene's expression may propagate through the network and affect other functionally relevant genes, making the co-expression pattern (rather than the single expression level) more reflective of disease biology. This would indicate that co-eQTL analyses such as this have captured changes in gene relationships (how genes co-vary across individuals), which might better reflect regulatory or pathway-level effects.

An important aspect of co-eQTLs is that they give insight into the mechanisms of eQTLs which allows for further insights such as the building of GRNs when there are layers of transcriptional regulation. We focussed many of our analyses on describing effects of direct TF regulation of eGenes (45.5% of our eGenes), but there are other mechanisms that can also explain why we see co-eQTL relationships between two genes. Beyond TFs these additional regulatory layers that may explain the remaining co-eQTLs might be non-coding RNA activity (many are non-polyadenylated, and therefore not captured in our assay) or alternative splicing. These alternative mechanisms merit further investigation as they may be able to explain the remaining missing regulatory logic captured by co-eQTLs.

### Supplementary Methods

#### Addition of GWAS variants to test

We broadened the genetic variants beyond the eQTL set to include variants implicated in "immune system diseases" (GWAS catalog) and also performed a full-window co-eQTL scan, testing all variants in the *cis*-window of a significant co-eQTL pair, to allow fine-mapping following our first focused co-eQTL scan. Some of the GWAS only variants did show a significant co-eQTL effect suggesting that we can find a co-eQTL effect where there is no eQTL effect. Also when we colocalized the co-eQTL credible sets to the eQTL credible sets from sc-eQTLGen, we observed several credible sets that did not colocalize. While the strongest effects did colocalize, the lack of overlap in other cases may be due to several factors.

#### Evaluation of co-expression measurement with simulations

Previous co-eQTL mapping papers have used Spearman correlation to calculate the co-expression of two genes to get a non-parametric robust estimation of co-expression. However, multiplexed 10X single-cell data is characterised by having large variation in the amount of cells and reads per individual which affects the reliability of the calculated correlation. As there are also versions of the Spearman and Pearson correlation which take into account weights per observation, allowing to weigh the observations based on their reliability (in our case reads per cell). We evaluated these correlation measures, using simulations to identify the most accurate and robust way to measure co-expression within single-cell data. We used a well-established Gaussian copula/ NORTA approach to simulate correlated gene-pairs that follow a negative binomial distribution which is the distribution usually observed for single-cell RNA-seq count data.
In a first step we specified for each simulated gene-pair (gene1 $X_1$, gene2 $X_2$) the true correlation (ρ) between the pair and the mean ($\mu_1$, $\mu_2$) and dispersion parameters ( $\theta_1$ , $\theta_2$) of the negative binomial distribution for each gene as well as the number of cells $N$.
We then used the ‘rnorm_multi’ function of the ‘faux’ package (version: 1.2.1) to generate two standard normal variables $Z_1$ (gene1) and $Z_2$ (gene2) with the specified correlation ρ and n observations (number of cells).

$$\begin{bmatrix} Z_1 \\ Z_2 \end{bmatrix} \sim \mathrm{N}\left(\begin{bmatrix} 0 \\ 0 \end{bmatrix}, \begin{bmatrix} 1 & \rho \\ \rho & 1 \end{bmatrix}\right)$$

Then we applied the normal cumulative distribution function (CDF; pnorm function in R), Φ, to each variable to transform them into uniform variables $U_1$ and $U_2$ in the range $[0, 1]$:

$$U_1 = \Phi(Z_1),\ U_2 = \Phi(Z_2)$$

Subsequently we used the inverse CDF (quantile function; qnbinom in R) of the negative binomial distribution, $F_{NB}^{-1}$, to transform the uniform variables into NB-distributed variables with the specified mean ($\mu_1$, $\mu_2$) and dispersion parameters ( $\theta_1$ , $\theta_2$) for each gene.

$$X_1 = F_{NB}^{-1}(U_1; \frac{\mu_1}{s_1}; \theta_1), X_2 = F_{NB}^{-1}(U_2; \frac{\mu_2}{s_2}; \theta_2)$$

Within the transformation the parameters $s_1, s_2$ are scaling factors to model different total counts per cell which are usually observed in single-cell RNAseq data. The parameter was sampled from scaling factors per cell calculated on the real single-cell input datasets as the mean counts across all cells and genes divided by the mean counts across all genes for one specific cell (i):

$s_n = \frac{Total\ Mean\ Counts}{Mean\ Counts_n} = \frac{\Sigma_n^N \Sigma_p^P x_{n,p}}{N\ x\ P} / \frac{\Sigma_p^P x_{n,p}}{P}$ , with $x_{n,p}$ = count for gene $p$ in cell $n$; $P$ = total number of genes ; $N$ = total number of cells.

We run different scenarios with this approach and in each scenario simulated count data for 10.000 gene-pairs where for 50% of the gene-pairs we simulated a difference in the correlation between different modeled genotypes (representing true co-eQTLs) and for others no difference in the correlation (representing true non- co-eQTLs). The selected exemplary specifications for those scenarios are based on distributions observed in the smallest dataset for different cell types (Wijst) (see Supplementary Table). The correlation difference was chosen to be rather small as this is also observed in real data :

| Scenario | Total nr. Gene-pairs per individual | Cells per individual | Simulated Genotypes | Nr. Individuals | ρ of gene-pairs |
|---|---|---|---|---|---|
| wijstB | 10000 | $\mu_s = 16$ $\sigma_s = 9$ | AA | 20 | 0 (10000) |
| | | | AB | 15 | 0 (5000); 0.05 (5000) |
| | | | BB | 10 | 0 (5000); 0.1 (5000) |
| wijstNK | 10000 | $\mu_s = 30$ $\sigma_s = 18$ | AA | 20 | 0 (10000) |
| | | | AB | 15 | 0 (5000); 0.05 (5000) |
| | | | BB | 10 | 0 (5000); 0.1 (5000) |
| wijstMono | 10000 | $\mu_s = 31$ $\sigma_s = 55$ | AA | 20 | 0 (10000) |
| | | | AB | 15 | 0 (5000); 0.05 (5000) |
| | | | BB | 10 | 0 (5000); 0.1 (5000) |
| wijstCD4_T | 10000 | $\mu_s = 262$ $\sigma_s = 126$ | AA | 20 | 0 (10000) |
| | | | AB | 15 | 0 (5000); 0.05 (5000) |
| | | | BB | 10 | 0 (5000); 0.1 (5000) |

The number of cells for each individual were sampled from a normal distribution with mean = $\mu_s$ and standard deviation = $\sigma_s$ as specified in the table so that each simulated individual had a different number of cells. The mean ($\mu_1$, $\mu_2$) parameters of the negative binomial distribution for each gene within these scenarios were sampled randomly from a range of specified $\mu$ parameters to reflect the range of observed mean counts in the real data. For gene1 of each gene-pair we also sampled the $\mu_1$ in a way that $\mu_1$differs between the 3 genotypes and by this reflects an eQTL effect on gene1. A comparison of the characteristics of the simulated counts and the real data (Wijst) are shown in Supplementary Figure XX.
After having simulated the count data for each individual of each of the scenarios we calculated the correlations on the simulated count data for each gene-pair for each individual with the different evaluated correlation measurements:

| Method | Function | R Package |
|---|---|---|
| Spearman | $cor(x_{1,n} * s_n, x_{2,n} * s_n, method = 'spearman')$ | stats (version 4.1.1) |
| Weighted Spearman | $weightedCorr(x_{1,n} * s_n, x_{2,n} * s_n, weights = c_n, method = 'spearman')$ | wCorr (version 1.9.8) |
| Pearson | $cor(x_{1,n} * s_n, x_{2,n} * s_n, method = 'pearson')$ | stats (version 4.1.1) |
| Weighted Pearson | $weightedCorr(x_{1,n} * s_n, x_{2,n} * s_n, weights = c_n, method = 'pearson')$ | wCorr (version 1.9.8) |

For the weighted correlations we use the weighting factor $c_n$ which specifies the number of genes with a read in cell $n$. So observations from cells with more non-zero reads across genes are higher weighted.
We then tested each gene-pair within each scenario for a co-eQTL effect by specifying a linear model (lm) with the correlation value as y and the genotype encoded as numeric covariate x. We only included a gene-pair of an individual in the testing when each of the genes has at least 10 non-zero counts in the simulated data.
To evaluate which of the evaluated correlation measures works best for eventual co-eQTL discovery we then compared two evaluation criteria among the four different methods. First we calculated the mean squared error between the estimated effect size (β) from the linear model and the specified difference in the correlation between the gene-pair (ρ):
$MSE = \frac{1}{L}\sum_{l=1}^{L}(\beta_l - \rho_l)^2$. Second we used the Bonferroni adjusted p-value from the linear model to plot ROC curves (ggplot2; version: 3.5.1, geom_roc), taking the adjusted p-value as threshold to specify whether a gene-pair is a true co-eQTL or not a co-eQTL based on the specified correlation difference for the simulation.
For both evaluations the weighted Pearson approach showed the best performance. Therefore, we used this approach in our co-eQTL mapping.

#### Definition of gene characteristic

Gene characteristics used in the decision tree analysis were:

- Mean absolute correlation (mean_abs_correlation) across individuals per gene-pair $l$: $\frac{1}{S}\sum_{s=1}^{S}\left|\rho_s\right|$
- Variance of the correlation (var_correlation) across individuals $s$ per gene-pair $l$: $\frac{1}{S-1}\sum_{s=1}^{S}\left(\rho_s - \overline{\rho}\right)^2$
- Weighted Variance of the correlation (weighted_variance) across individuals per gene-pair $l$: $(\sum_{s=1}^{S} N_s\left(\rho_s - \overline{\rho_w}\right)^2)/(\sum_{s=1}^{S} N_s)$ , $with\ \overline{\rho_w} = (\sum_{s=1}^{S} N_s * \rho_s)/(\sum_{s=1}^{S} N_s)$
- Number of individuals with correlation values (n_not_NA) per gene-pair l.
- Percentage of significant correlations (perc_significant; p-value < 0.05) across individuals per gene-pair $l$.
- The mean amount of cells with a zero count in at least one gene per gene pair (gene1, gene2) (mean_zero_one_cell), which corresponds to calculating for each gene-pair $l$ for each individual $s$ the sum of the amount of cells where either gene1 or gene2 or both have a zero count. Then the mean of this value per gene pair $l$ (gene1, gene2) across all individuals is calculated.
$\frac{1}{S}\sum_{s=1}^{S}\sum_{n=1}^{N_s} v_{s,n},\ with\ v_{s,n} = \begin{cases} 1\ if\ x_{s,1,n} == 0\ |\ x_{s,2,n} == 0 \\ 0\ if\ x_{s,1,n} > 0\ \&\ x_{s,2,n} > 0 \end{cases}$
- The mean amount of cells with a zero count in both genes per gene pair ( mean_paired_zero): which corresponds to calculating for each gene-pair (gene1, gene2) for each individual the sum of the amount of cells where both gene1 and gene2 have a zero count. Then the mean of this value per gene pair across all individuals is calculated.
$\frac{1}{S}\sum_{s=1}^{S}\sum_{n=1}^{N_s} v_{s,n},\ with\ v_{s,n} = \begin{cases} 1\ if\ x_{s,1,n} == 0\ \&\ x_{s,2,n} == 0 \\ 0\ if\ x_{s,1,n} > 0\ |\ x_{s,2,n} > 0 \end{cases}$
- The mean amount of cells with a non-zero count in both genes per gene pair ( mean_paired_non_zero): which corresponds to calculating for each gene-pair (gene1, gene2) for each individual the sum of the amount of cells where both gene1 and gene2 have a count value that is not zero. Then the mean of this value per gene pair across all individuals is calculated.
$\frac{1}{S}\sum_{s=1}^{S}\sum_{n=1}^{N_s} v_{s,n},\ with\ v_{s,n} = \begin{cases} 1\ if\ x_{s,1,n} > 0\ \&\ x_{s,2,n} > 0 \\ 0\ if\ x_{s,1,n} == 0\ |\ x_{s,2,n} == 0 \end{cases}$
- Minimum mean expression per gene-pair (min_mean_expression), which corresponds to calculating the mean expression for gene1 and gene2 across all cells of each individual and then taking the mean across individuals. The minimum of both genes (gene1,gene2) of the gene-pair is the minimum mean expression per gene pair.
$min\ (\frac{1}{S}\sum_{s=1}^{S}(\frac{1}{N_s}\sum_{n=1}^{N_s} x_{s,1,n})\ ;\ \frac{1}{S}\sum_{s=1}^{S}(\frac{1}{N_s}\sum_{n=1}^{N_s} x_{s,2,n}))$
- Minimum mean variance per gene pair (min_mean_variance), which corresponds to calculating the variance of the expression across cells for each individual and gene

and then calculating the mean of this variance across all individuals for each gene. Then the minimum of this value of both genes (gene1, gene2) of the gene-pair is selected per gene-pair.

$$min\,\left(\frac{1}{S}\sum_{s=1}^{S}\left(\frac{1}{N_s-1}\sum_{n=1}^{N_s}(x_{s,1,n}-\overline{x_{s,1}})^2\right)\;;\;\frac{1}{S}\sum_{s=1}^{S}\left(\frac{1}{N_s-1}\sum_{n=1}^{N_s}(x_{s,2,n}-\overline{x_{s,2}})^2\right)\right)$$

- Minimum of the sum of non-zero counts per gene of the gene-pair (min_sum_non_zero), which corresponds to calculating for both gene1 and gene2 of the gene-pair for each individual the number of cells in which the gene has a count value that is not zero. Then summing up this number across all individuals for gene1 and gene2 and taking the minimum value of both genes for the gene-pair.

$$min\,\left(\sum_{s=1}^{S}\sum_{n=1}^{N_s}v_{s,1,n}\;;\;\sum_{s=1}^{S}\sum_{n=1}^{N_s}v_{s,2,n}\right),\text{ with } v_{s,1,n}=\begin{cases}1\;if\;x_{s,1,n}>0\\0\;if\;x_{s,1,n}==0\end{cases};\;v_{s,2,n}=\begin{cases}1\;if\;x_{s,2,n}>0\\0\;if\;x_{s,2,n}==0\end{cases}$$

- Maximum mean percentage of zero per gene pair (max_mean_percentage_zero), which corresponds to calculating the percentage of zeros (meaning cells for which a 0 count was measured) across cells for each individual and gene and then calculating the mean of this percentage across all individuals for the gene. Then the maximum of this value of both genes (gene1, gene2) of the gene-pair is selected per gene-pair.

$$max\,\left(\frac{1}{S}\sum_{s=1}^{S}\frac{1}{N_s}\sum_{n=1}^{N_s}v_{s,1,n}\;;\;\frac{1}{S}\sum_{s=1}^{S}\frac{1}{N_s}\sum_{n=1}^{N_s}v_{s,2,n}\right),\text{ with } v_{s,1,n}=\begin{cases}1\;if\;x_{s,1,n}==0\\0\;if\;x_{s,1,n}>0\end{cases};\;v_{s,2,n}=\begin{cases}1\;if\;x_{s,2,n}==0\\0\;if\;x_{s,2,n}>0\end{cases}$$

With:

- S = total number of individuals
- $\rho_s$ = weighted pearson correlation of individual $s$
- $N_s$ is total number of cells of individual $s$
- $x_{s,1,n}$ = counts of gene 1 in cell $n$ of individual $s$
- $x_{s,2,p}$ = counts of gene 2 in cell $n$ of individual $s$

### Supplementary Tables

Link to Tables :
https://docs.google.com/spreadsheets/d/1eAT26HU1CbpbPArUZkRcIEyju46Fmbo1DrQAZ7k1AV8/edit?gid=0#gid=0

**Suppl. Table 1. Overview of Datasets**
Overview of number of samples and cells in each dataset.

**Suppl. Table 2. Annotated co-eQTLs**
Annotated significant co-eQTLs after meta-analysis.

**Suppl. Table 3. Average cells per individual in cell type vs number of co-eQTLs**
Per cell type the average number of cells per donor and the number of identified co-eQTLs.

**Suppl. Table 4. Replication interaction**
Table of results of replication in independent bulk (BIOS) data.

**Suppl. Table 5. Cell type composition effects**
Overview of the number of significant co-eQTLs with identified cell type composition effects.

**Suppl. Table 6. Credible sets shared across cell types**
Corresponding number of credible sets that colocalize across a number of cell types. Data from co-eQTL and sc-eQTLGen projects.

**Suppl. Table 7. eGene GWAS enrichment 2x2**
Data for calculating Fishers' exact enrichment for eGene coloc. method.

**Suppl. Table 8. Variant based GWAS enrichment 2x2**
Data for calculating Fishers' exact enrichment for variant LD method.

**Suppl. Table 9. eGene GWAS enrichment odds ratio**
Statistics overview of eGene GWAS enrichment.

**Suppl. Table 10. Variant based GWAS enrichment odds ratio**
Statistics overview of variant GWAS enrichment.

### Supplementary Figures

**Suppl. Fig. 1.** Dataset overview/cell-distributions
**Suppl. Fig. 2.** Evaluation of correlation measure: weighted Pearson
**Suppl. Fig. 3.** Decision tree
**Suppl. Fig. 4.** Cells per individual vs number of co-eQTLs found
**Suppl. Fig. 5.** Distribution of effects and locations of eGenes/ co-eGenes. Location of eGenes/ co-egenes
**Suppl. Fig. 6.** Within dataset replication: heatmap
**Suppl. Fig. 7.** Meta-analysis vs OneK1K
**Suppl. Fig. 8.** Fine-mapping Supplement
**Suppl. Fig. 9.** Cell-type-specificity: upset plots
**Suppl. Fig. 10.** Cell-type-specificity: scatter plots
**Suppl. Fig. 11.** GWAS enrichment with different filters
**Suppl. Fig. 12.** Mostafavi eGene plots for different cell types
**Suppl. Fig. 13.** Effect of the number of credible sets on co-eQTL
**Suppl. Fig. 14.** Effect of the number of credible sets on GWAS coloc.
**Suppl. Fig. 15.** Relationship of the number of CS and number of co-eGenes
**Suppl. Fig. 16.** Mostafavi co-eGene plots for the different cell types
**Suppl. Fig. 17.** GWAS colocalization co-eQTL overview + TF mechanisms
**Suppl. Fig. 18.** micro-RNA enrichment
**Suppl. Fig. 19.** Figure for ID2/ IL7R example
**Suppl. Fig. 20.** RPS26 - NR4A3 example
**Suppl. Fig. 21.** IKZF2-SH3YL1 example
**Suppl. Fig. 22.** BACH2- AP1S3 example

Suppl. Fig. 1

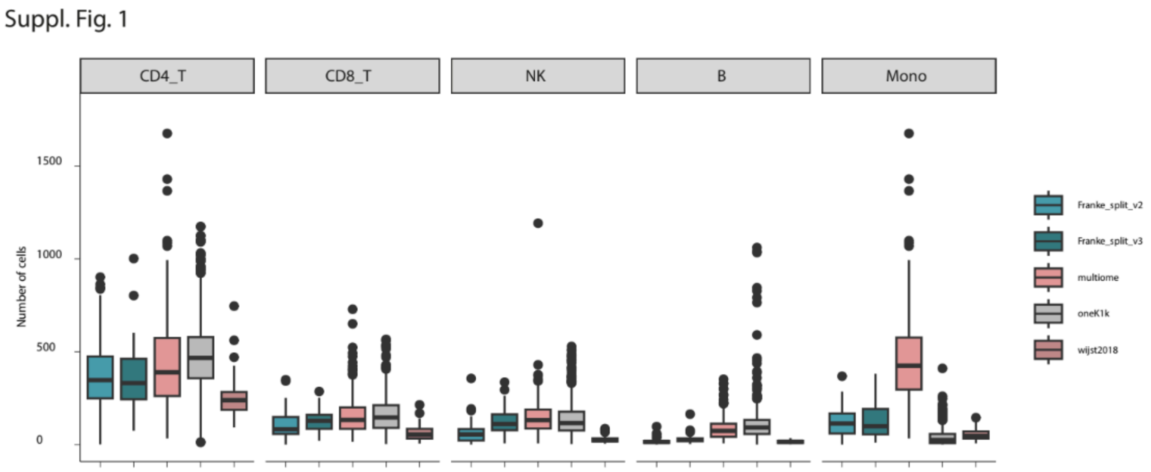


**Suppl. Fig. 1:** Distribution of the number of cells in the different datasets and cell types.

### Suppl. Fig. 2

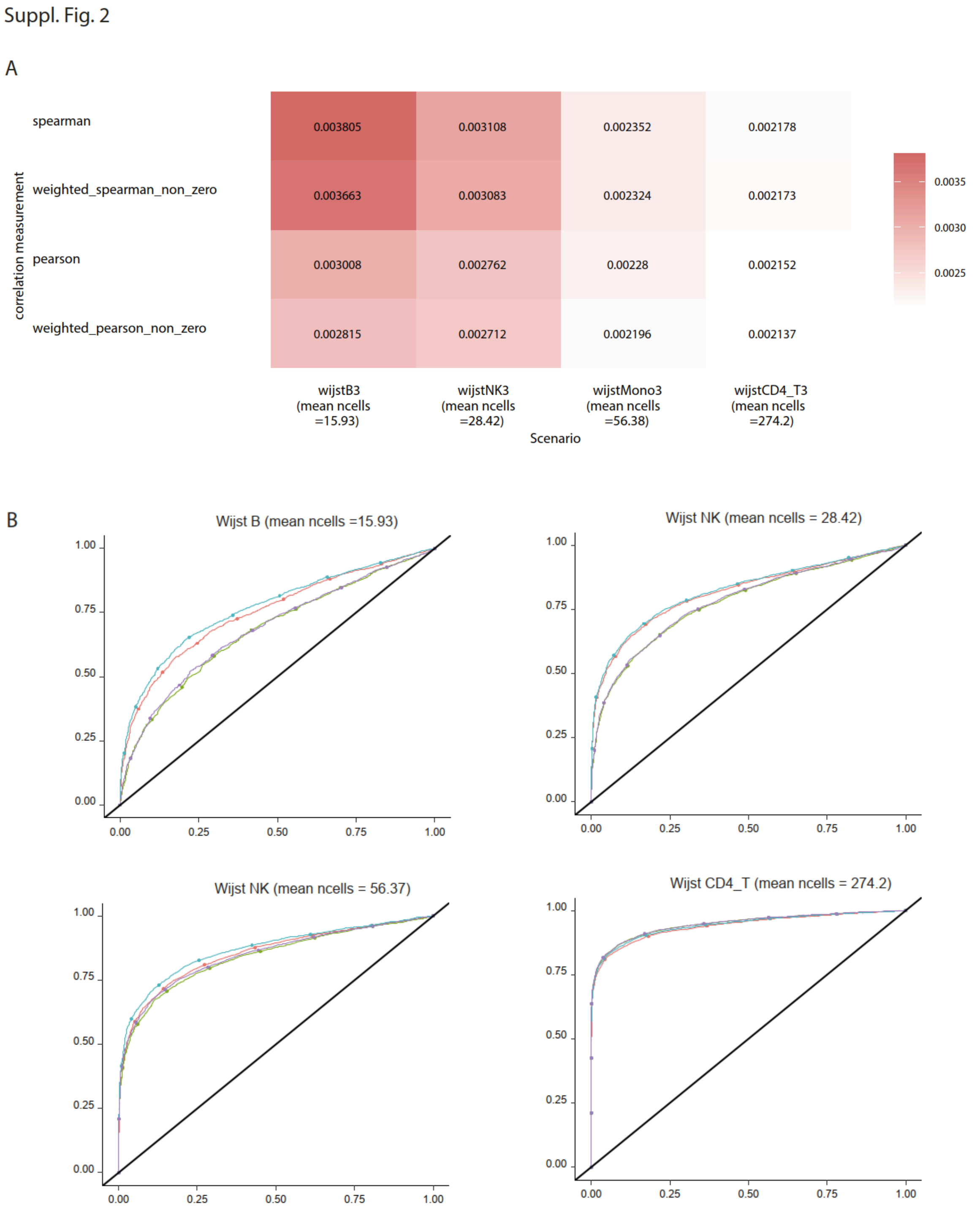


**Suppl. Fig. 2:** Comparison of different correlation measures on simulated data. **a)** MSE comparing true simulated co-eQTL effect to estimated co-eQTL effect using the different correlation measures. **b)** ROC curves using the p-value from co-eQTL mapping on simulated data with different correlation measures.

### Suppl. Fig. 3

Suppl. Fig. 3

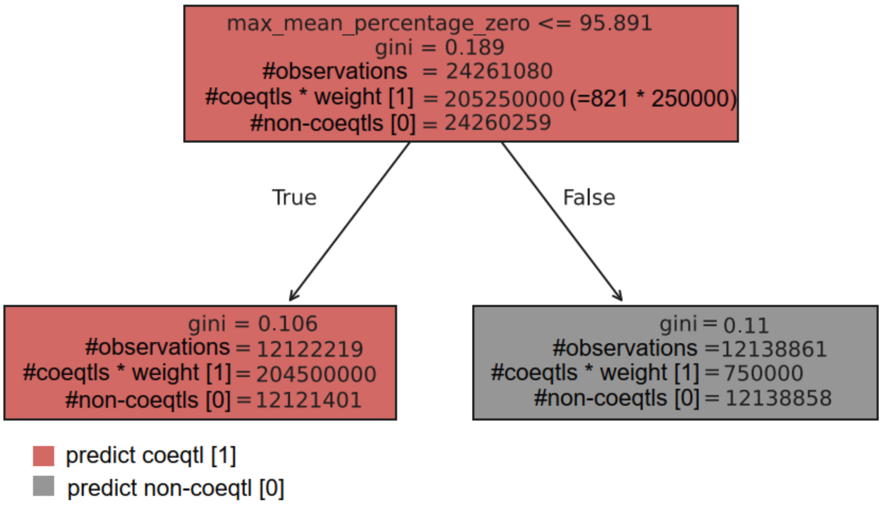


| | Training | Test |
|---|---|---|
| Accuracy on significant co-eQTLs | (1- (3/821)) * 100 = 99.63% | (1-1/208) * 100 = 99.52% |
| Accuracy on non co-eQTLs | (1 - (12,138,858/24,260,259)) * 100 = 49.96% | (1- (3,034,503/(6,065,063))) * 100 = 49.97 % |

**Suppl. Fig. 3:** Decision tree result on the training data. **a)** Showing the number of tested triplets that are predicted to be either a significant co-eQTL or not when using the feature “maximum mean percentage zero” and the threshold chosen by training the tree. Co-eQTL occurrences have been upweighted by the factor 250.000. **b)** Overview of decision tree classification performance on training and test data predicting significant co-eQTLs (y=1) vs. non-significant co-eQTLs (y = 0).

Suppl. Fig. 4

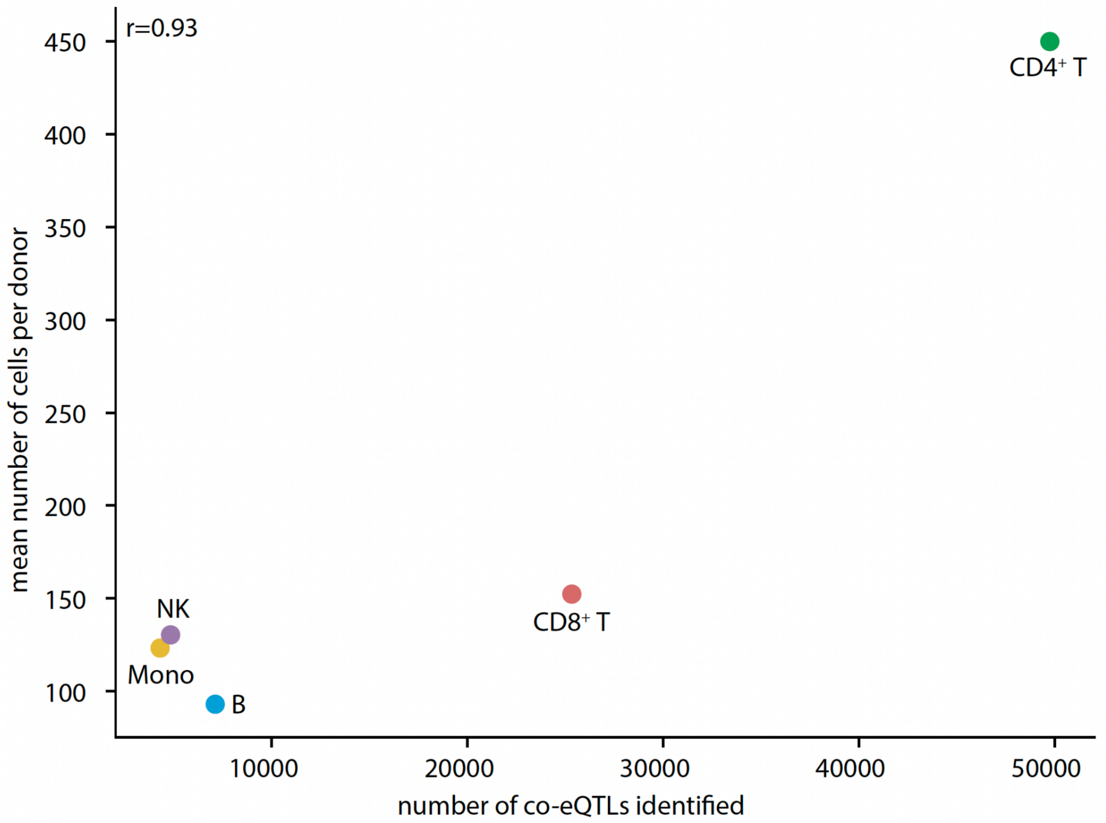


**Suppl. Fig. 4:** The number of co-eQTLs identified is strongly correlated to the mean number of cells per individual.

Suppl. Fig. 5

A

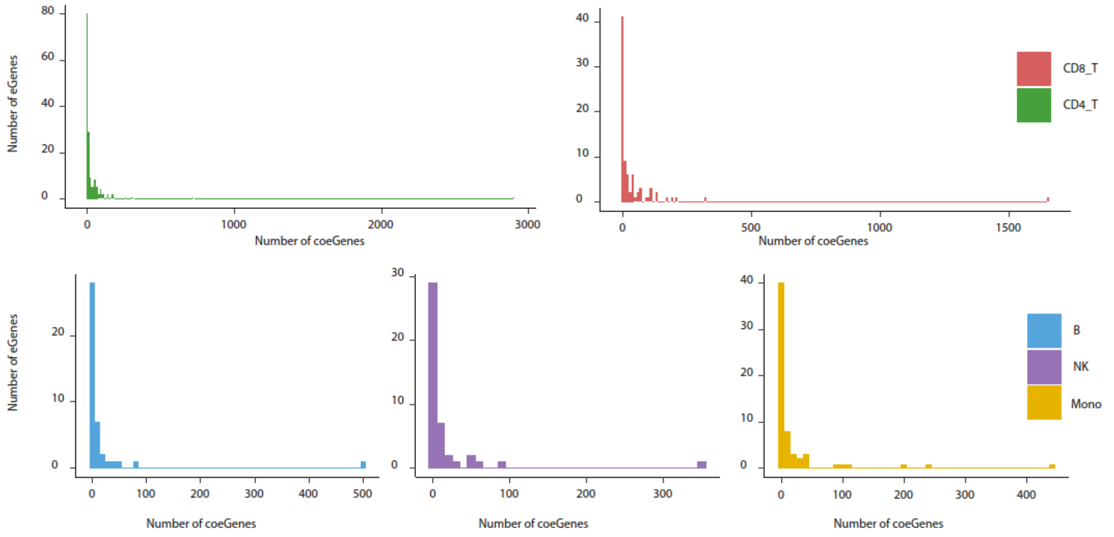


B CD4_T

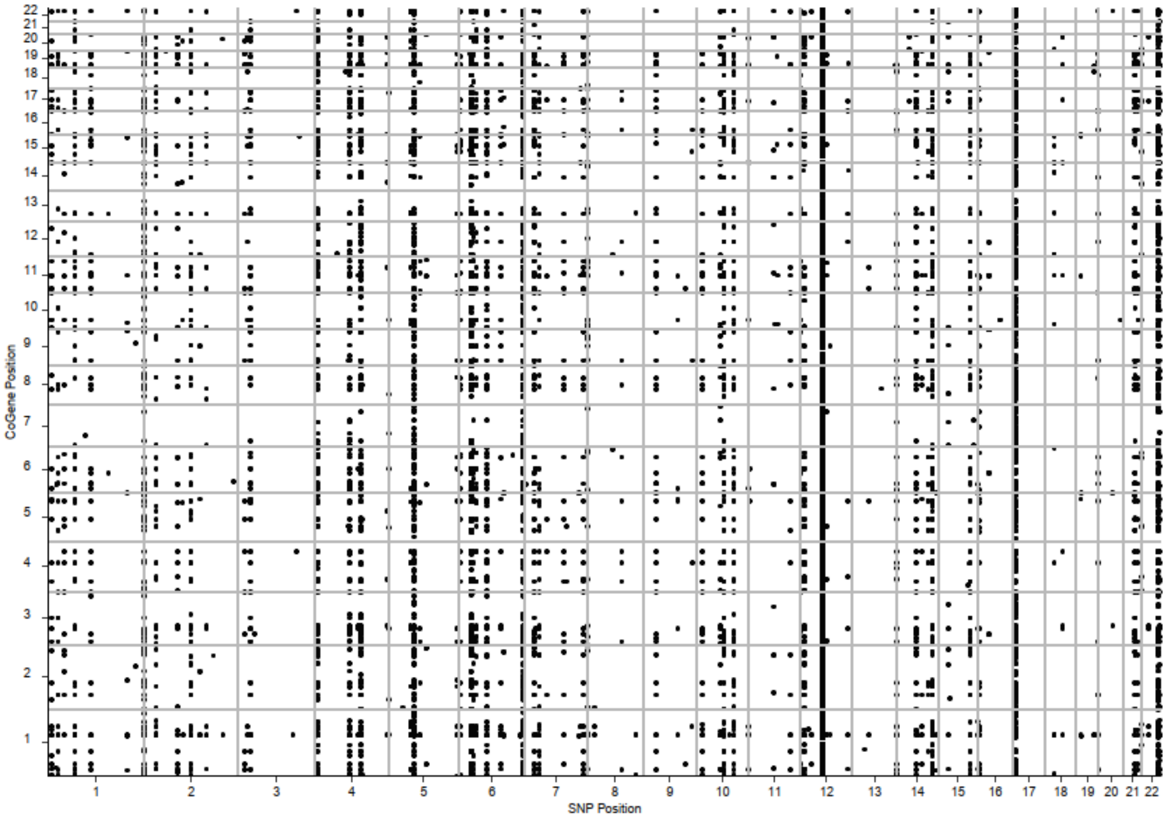

CD8_T

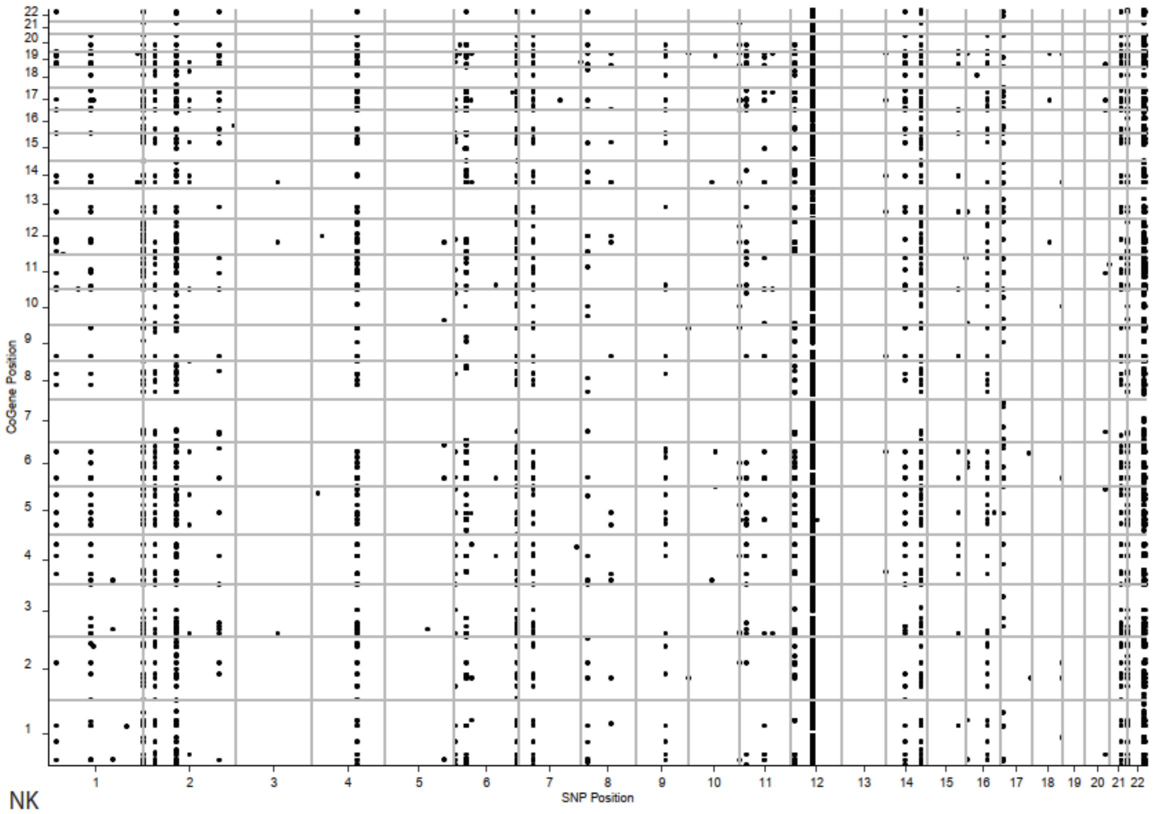
CoGene Position
SNP Position

NK

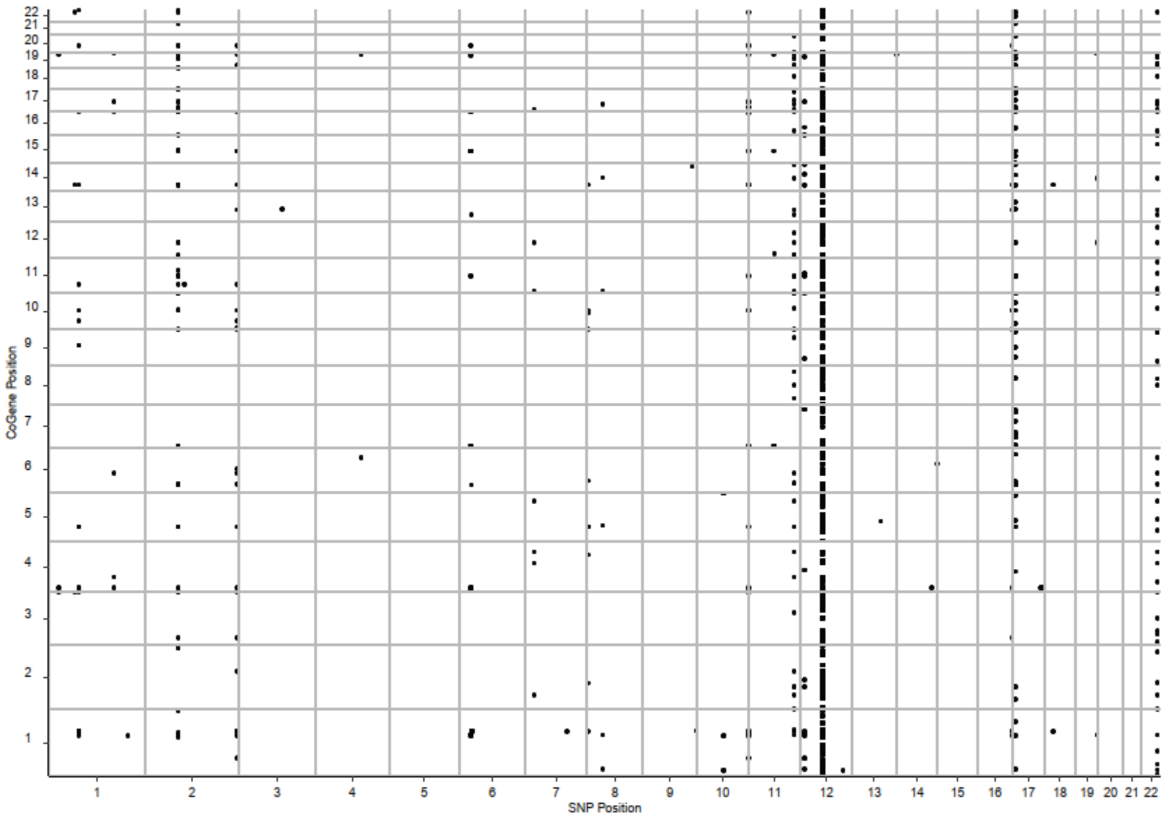
CoGene Position
SNP Position

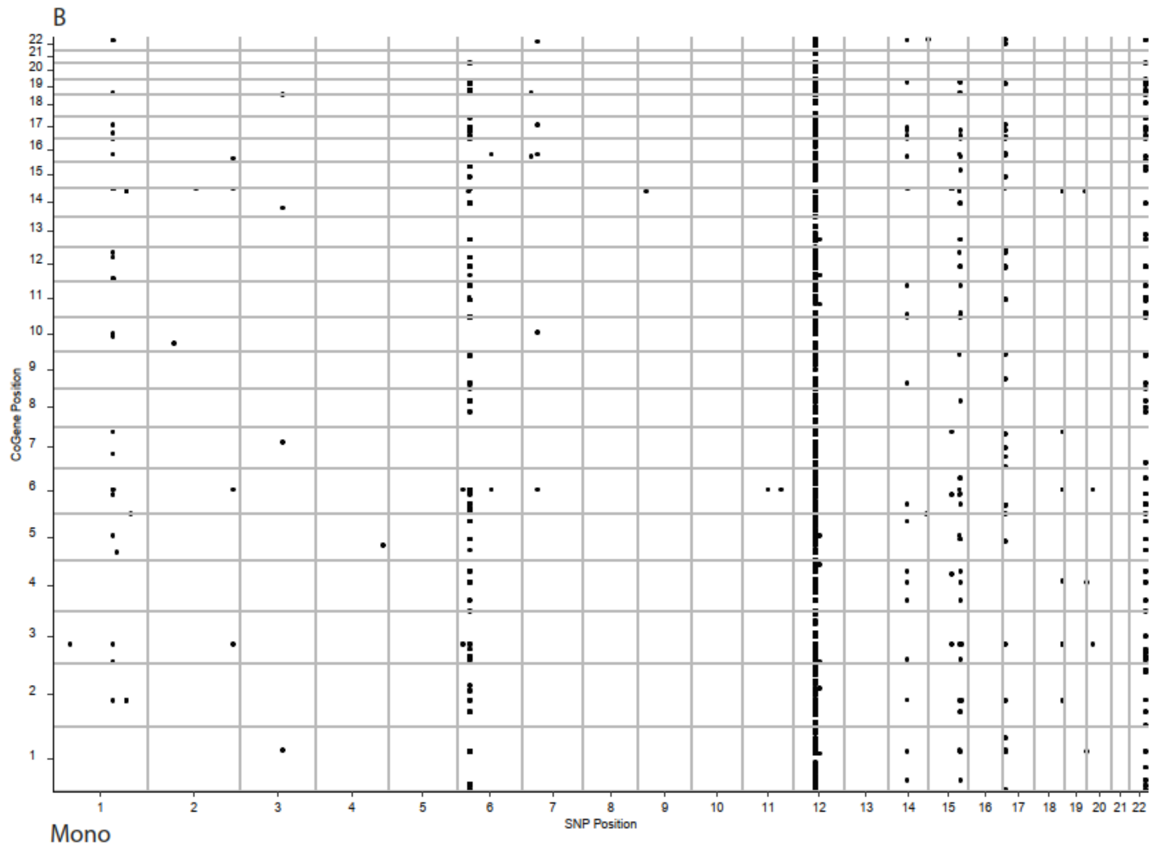


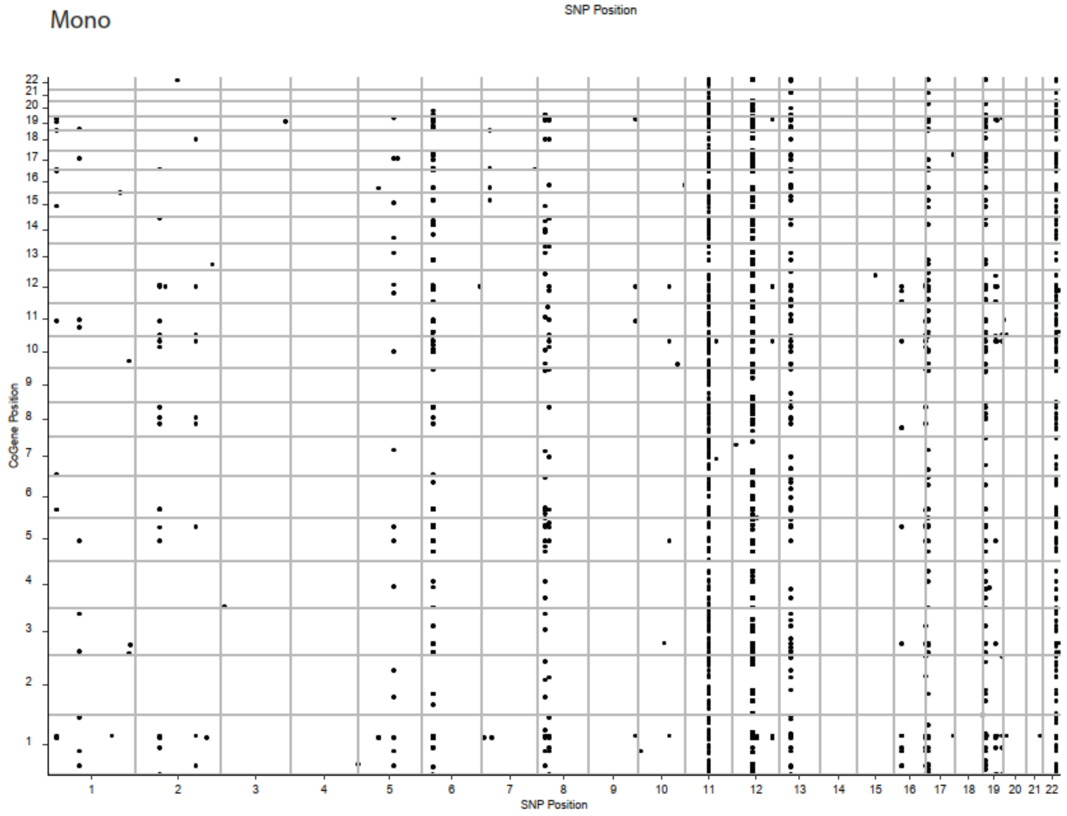


**Suppl. Fig. 5:** Distribution of effects. **a)** Histograms showing how many eGenes have a certain number of co-eGenes. **b)** Scatter plot for significant co-eQTLs showing position of co-eGenes compared to variants. Each point reflects a significant gene pair. In case of multiple significant variants per gene pair, the entry with the lowest p-value is shown.

Suppl. Fig. 6

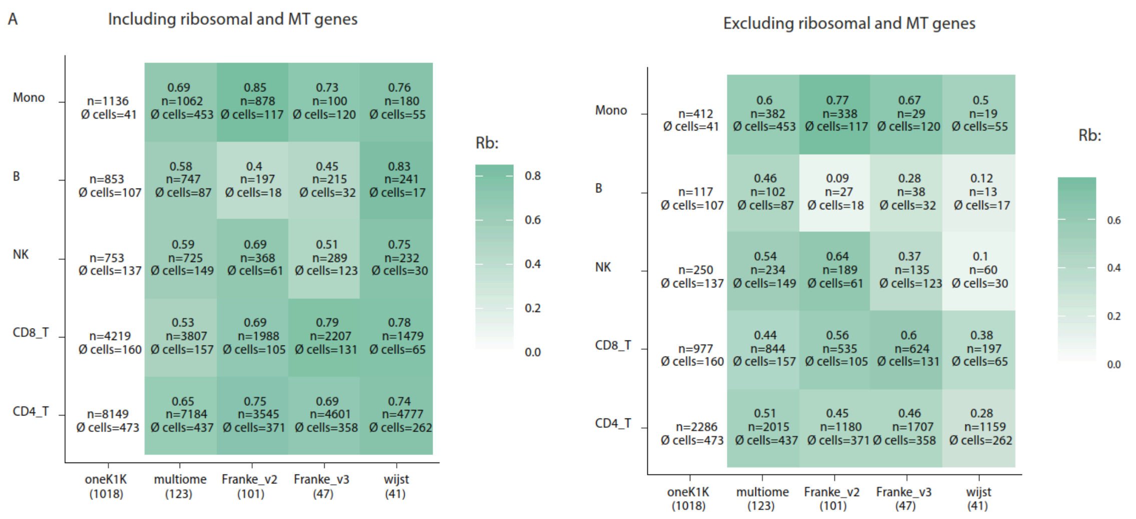


**Suppl. Fig. 6:** Across dataset replication of co-eQTLs discovered in OneK1K dataset and replicated in the other datasets. Replication measured by correlation of z-scores. Text inside the heatmap indicates the correlation value, the number of replicated co-eQTLs (n) and the average number of cells per individual in that cell type and dataset. For OneK1K no correlation values are shown as OneK1K effects were replicated in the other datasets. Effects were correlated on a gene pair level, the strongest effect per gene pair was selected. For the comparison on the right side all gene pairs including ribosomal or mitochondrial genes were excluded.

Suppl. Fig. 7

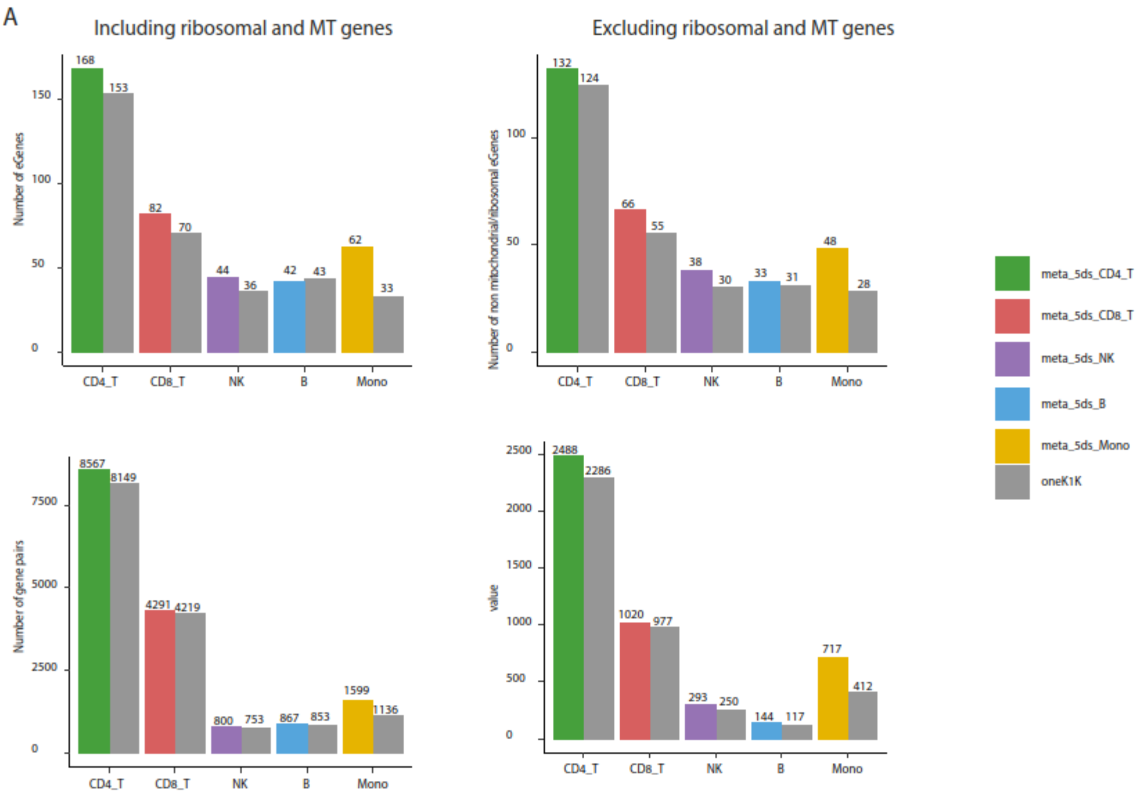


**Suppl. Fig. 7:** Comparison of number of eGenes and gene pairs with significant co-eQTL effect for OneK1K vs. the meta-analysis across all 5 datasets (meta_5ds) and per cell type.

Suppl. Fig. 8

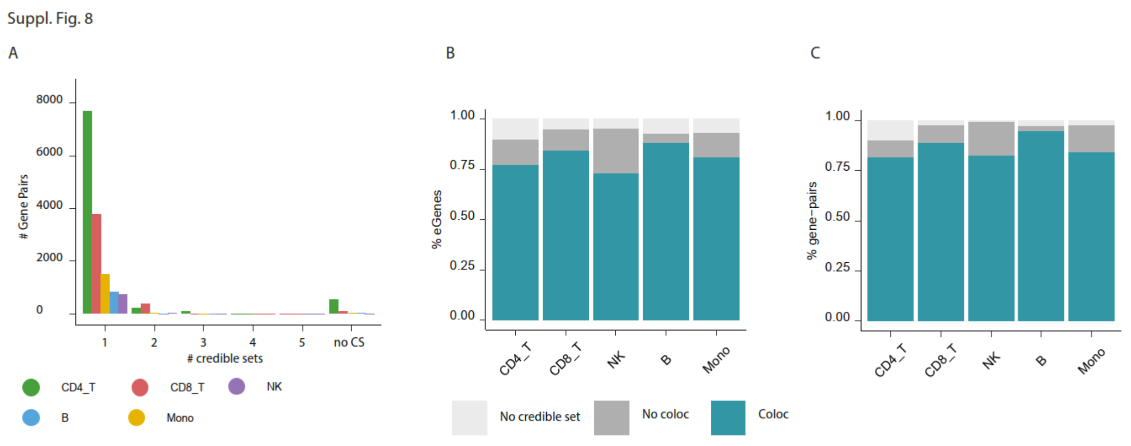


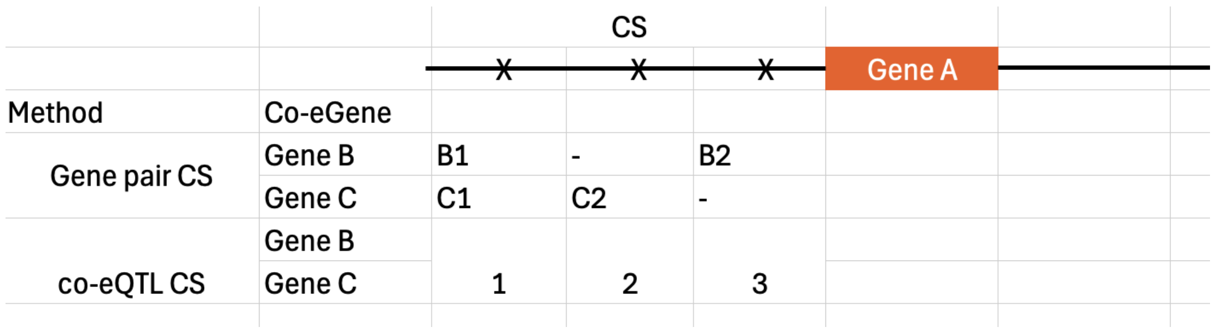


**Suppl. Fig. 8: a)** Number of credible sets per gene pair. **b)** Proportion of eGenes for which at least one eQTL credible set colocalized with at least one co-eQTL credible set. **c)** Proportion of gene pairs for which at least one "gene pair credible set" colocalized with one eQTL credible set. **d)** Difference between “gene pair credible sets” and “co-eQTL credible sets”. For gene A being the eGene, and gene B and C being co-eGenes for gene A. At a gene pair credible set level each co-eGene (gene B and gene C) has two credible sets identified with gene A. At a co-eQTL credible set level there are three credible sets identified in total, this is done by colocalizing the credible sets across gene B and gene C.

Suppl. Fig. 9

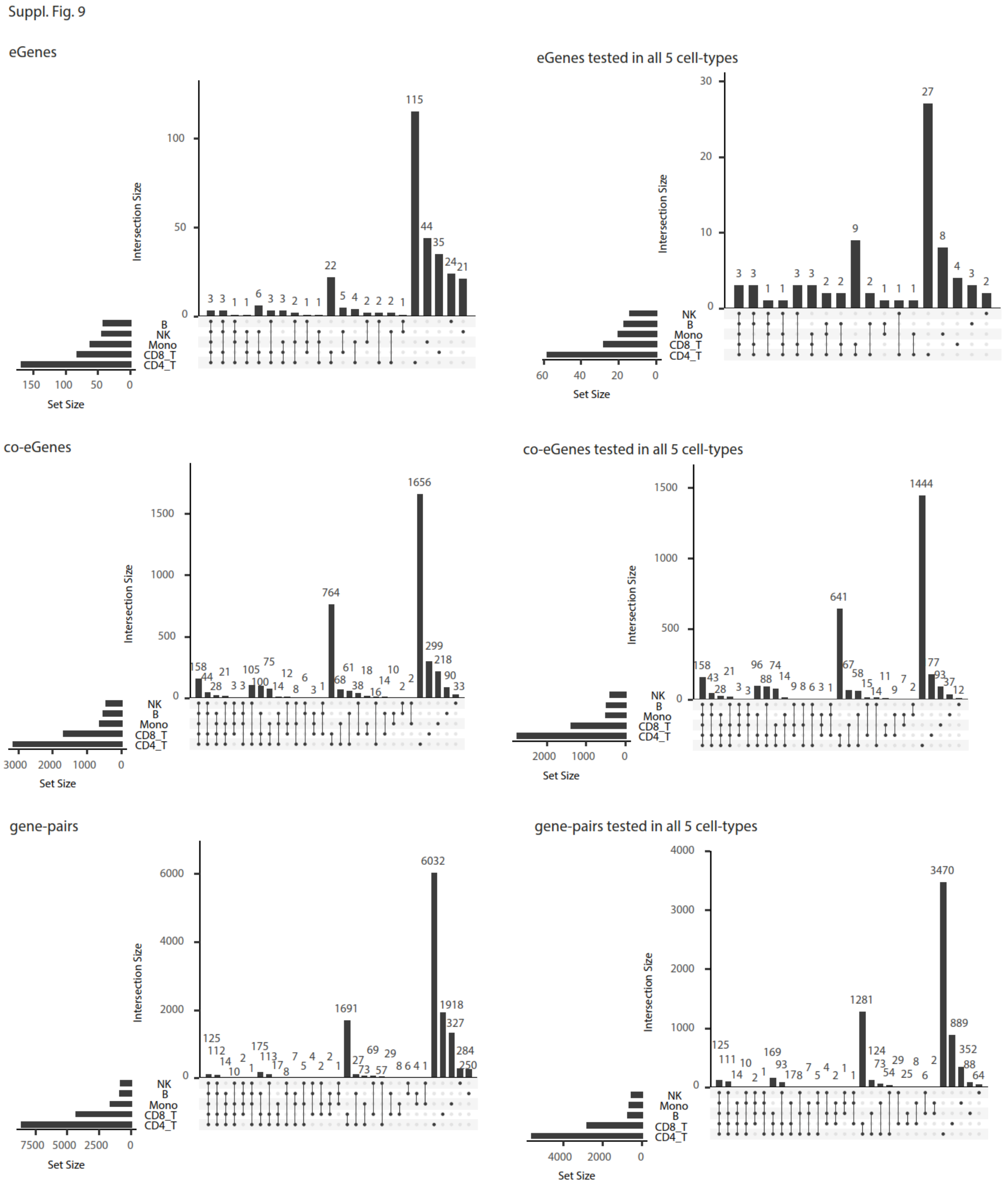


**Suppl. Fig. 9:** Upset plots showing the overlap of significant co-eQTL effects across the different cell types for eGenes, co-eGenes and gene pairs. Plots on the right side were filtered on eGenes, co-eGenes and gene pairs that were tested in all 5 cell types respectively.

Suppl. Fig. 10

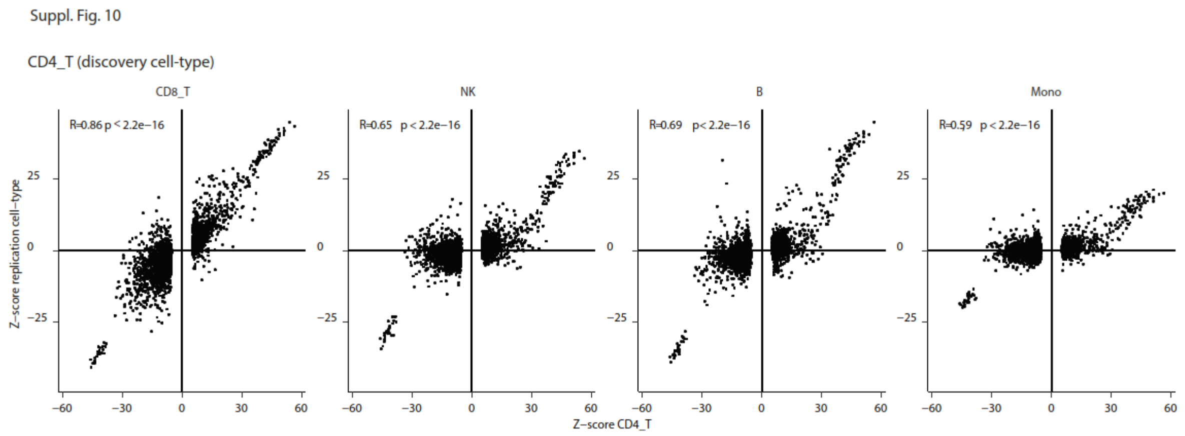


**Suppl. Fig. 10:** Correlation of z-scores for co-eQTLs identified in cell type indicated as discovery cell type and tested in the indicated replication cell types. Scatterplots illustrated correlations shown in Fig. 3 e,h.

#### Suppl. Fig. 11

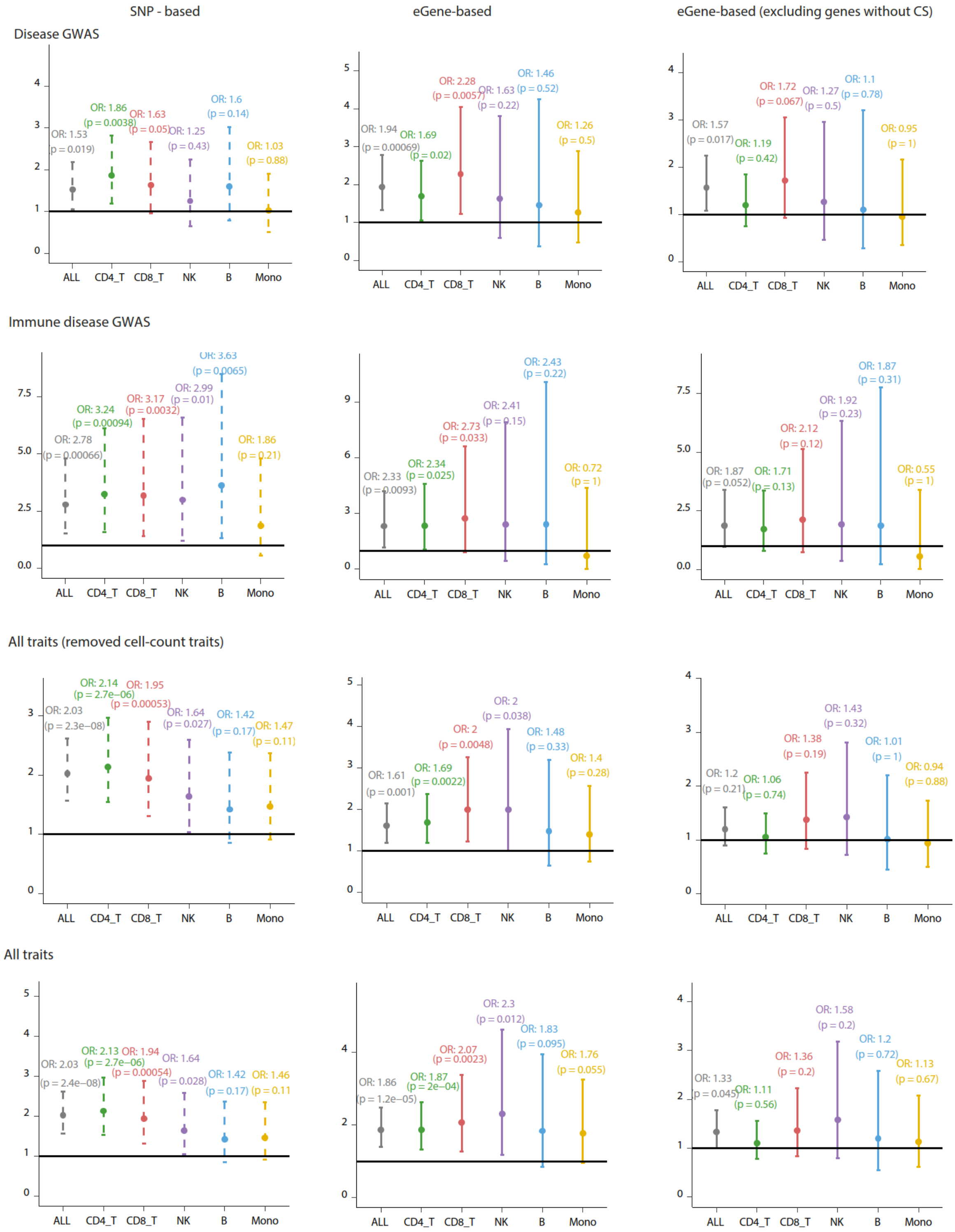


**Suppl. Fig. 11:** Odds ratio of enrichment of GWAS terms shown for different subsets of GWAS categories (based on EFO terms) and different analysis approaches comparing variants (variant-based) and eGenes (eGene-based) with co-eQTL effect to variants and eGenes without co-eQTL effect. GWAS subsets defined as: All traits (no filtering of GWAS terms); All traits (removed cell-count traits); Disease and Immune-Disease (Methods). Different analysis methods included: variant-based and two versions of the eGene-based analysis including and excluding eGenes without credible set (Methods).

Suppl. Fig. 12

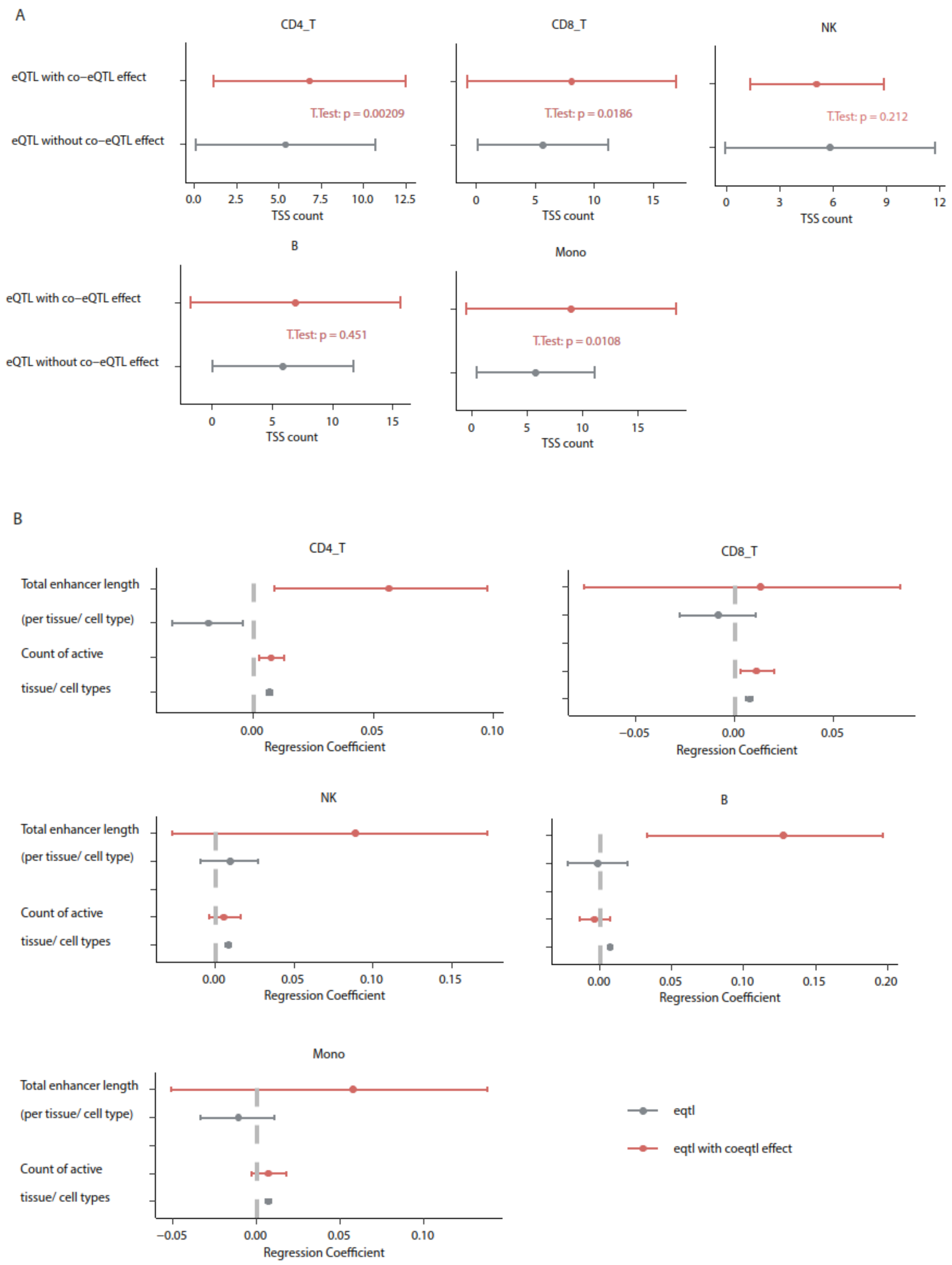


**Suppl. Fig. 12:** TSS differences and logistic regression model coefficients for analysis of Fig. 3b for the different single-cell types. **a)** Mean count of TSSs per gene in the FANTOM project for eGenes with co-eQTL effect compared to eGenes without co-eQTL effect. **b)** Enhancer features: logistic regression coefficients for enhancer features derived from the Roadmap dataset (total enhancer length, count of active tissue/cell types) for predicting co-eQTL hits (red) and eQTL hits (grey) versus random variants (n = 100,000) after adjusting for confounders (Methods).

Suppl. Fig. 13

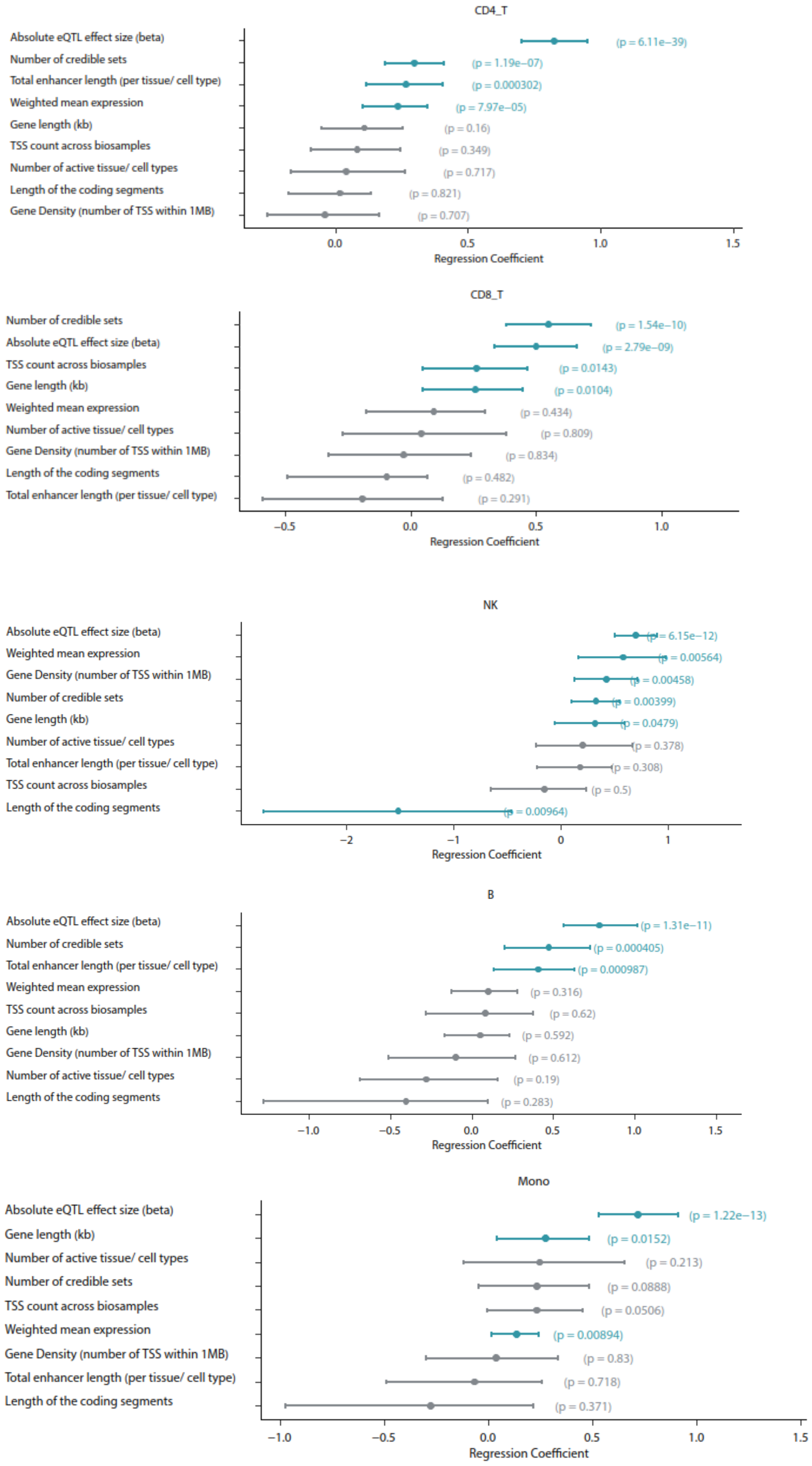


**Suppl. Fig. 13:** Cell-type-specific results for logistic regression model of Fig. 4c. Logistic regression coefficients of model predicting co-eQTL hits vs. classical eQTL hits.

### Suppl. Fig. 14

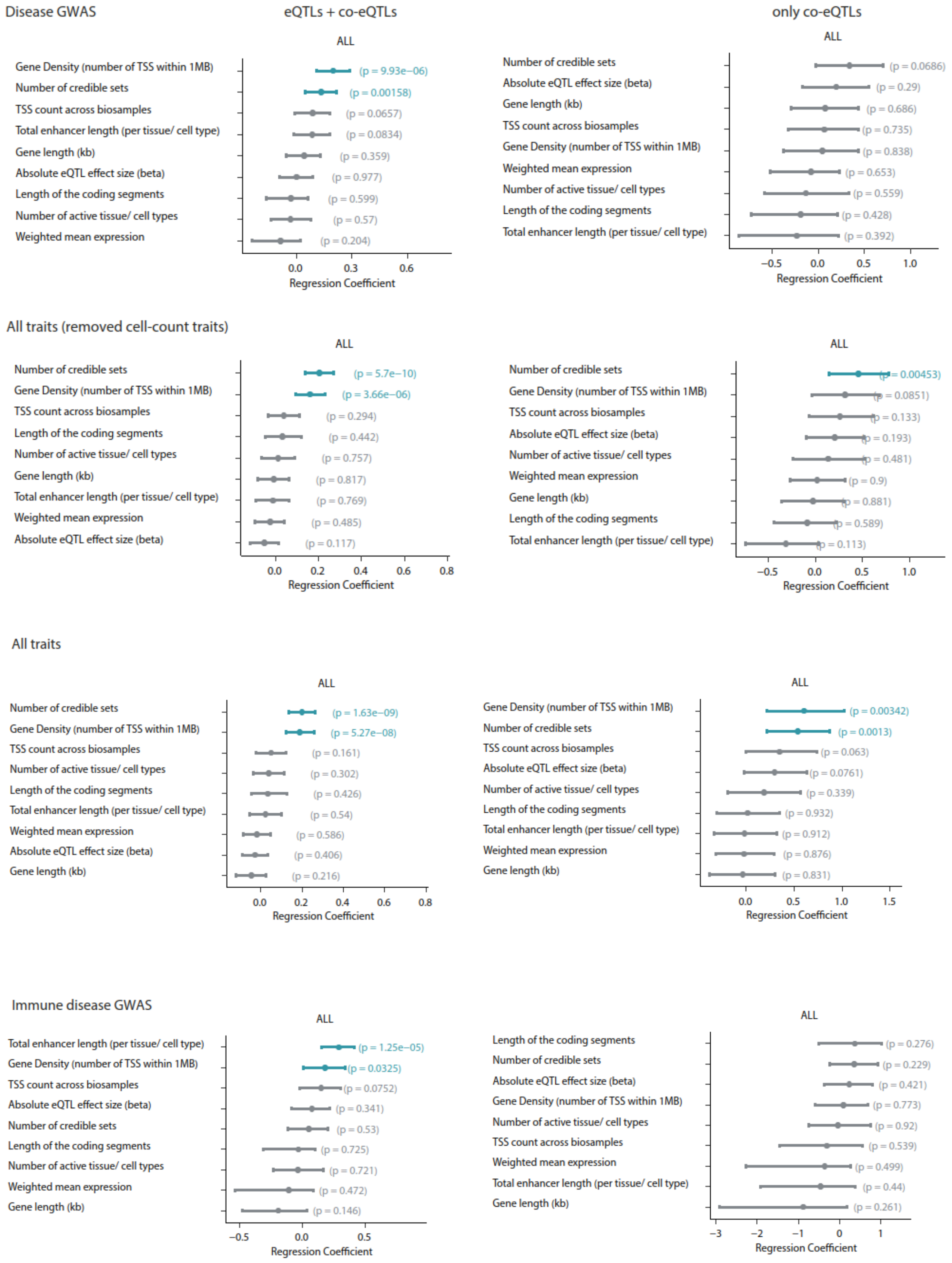


**Suppl. Fig. 14:** Logistic regression models to evaluate the influence of the number of credible sets and other gene features on the GWAS colocalization (predicting GWAS colocalization vs. no GWAS colocalization). The model was run for different subsets of GWAS traits as presented in Suppl. Fig 11 (rows) and including all eGenes tested for co-eQTL effects (left) vs. only eGenes with co-eQTL effect (right).

Suppl. Fig. 15

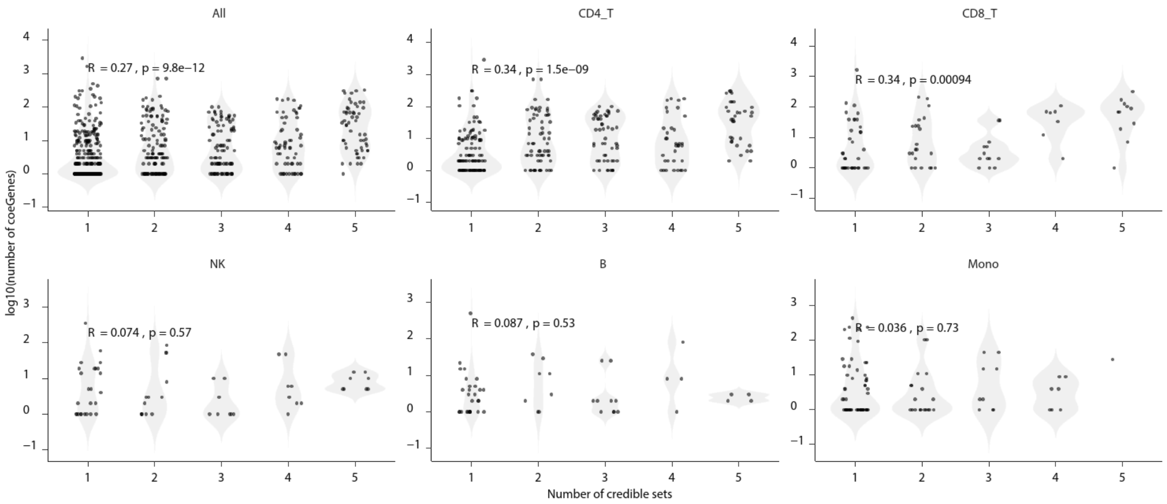


**Suppl. Fig. 15:** Evaluation of the relationship between the number of credible sets of an eGene and the number of associated co-eGenes per cell type and across all cell types (first panel). Spearman correlation coefficients (R) are shown.

### Suppl. Fig. 16

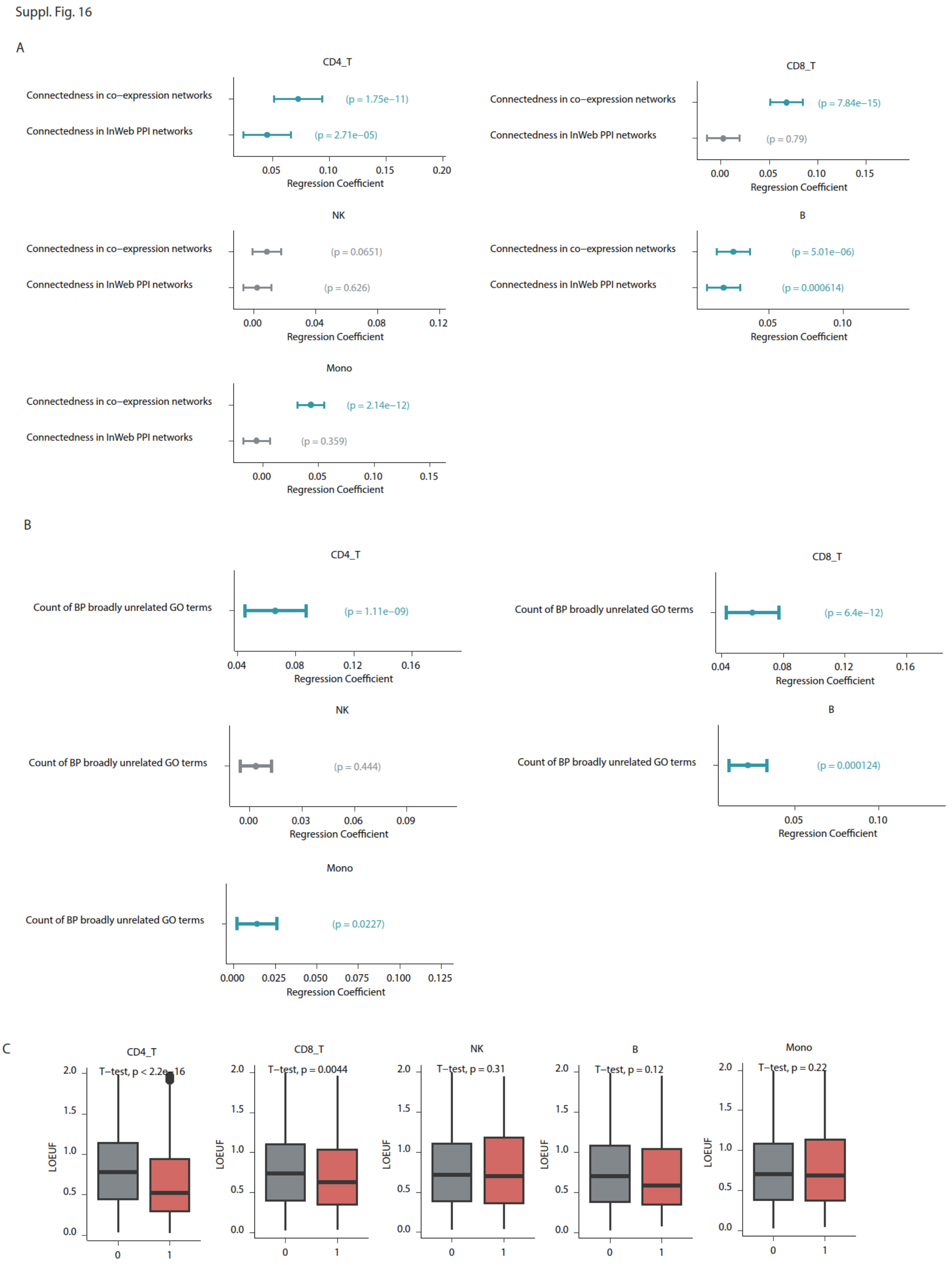


**Suppl. Fig. 16:** Evaluation of co-eGene characteristics comparing co-eGenes to all tested genes per cell type as presented across all cell types in Fig.4. **a, b)** Regression coefficients of linear models on the residuals of predicting significant co-eGenes vs. tested genes after correcting for the mean expression. **c)** LOEUF scores for significant co-eGenes vs. tested genes.

Suppl. Fig. 17

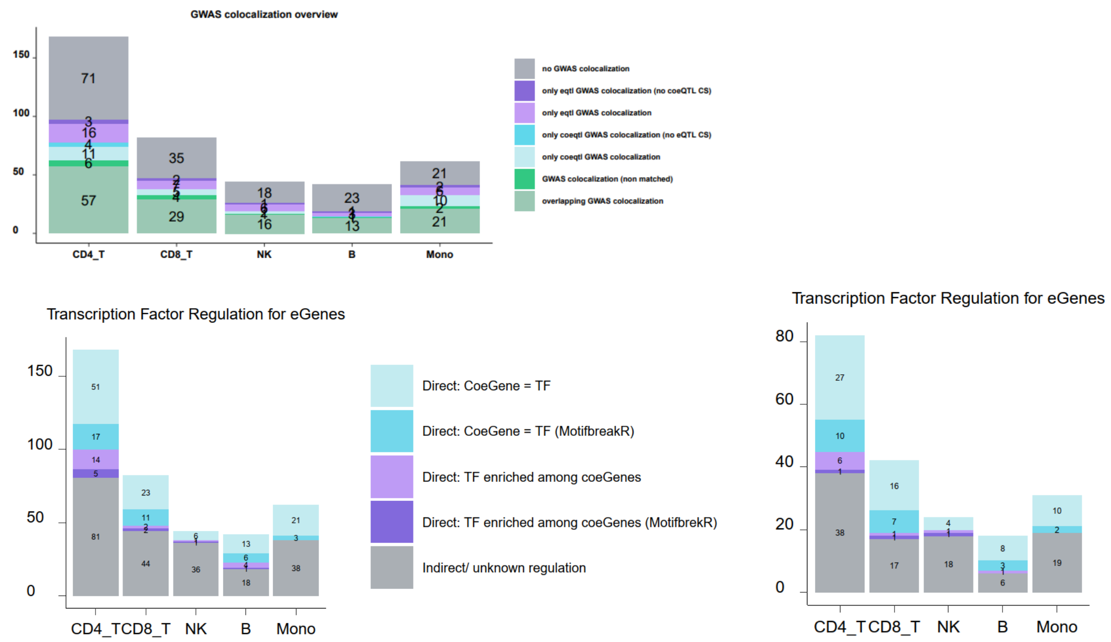


**Suppl. Fig. 17:** Proportion of eGenes with supporting evidence. **a)** Proportion of eGenes (both eQTL and co-eQTL, split by cell type) showing colocalization to GWAS. Split by eQTL (purple), co-eQTL (blue) or both (green). **b)** Proportion of co-eQTL eGenes with evidence for TF. Similar to fig. 5a except split per cell type, separated on whether the TF is part of identified co-eGenes (blue) or identified through enrichment analysis of identified co-eGenes (purple). Left plot shows all eGenes with identified TF, right plot shows identified TF when there was coloc. to disease.

Suppl. Fig. 18

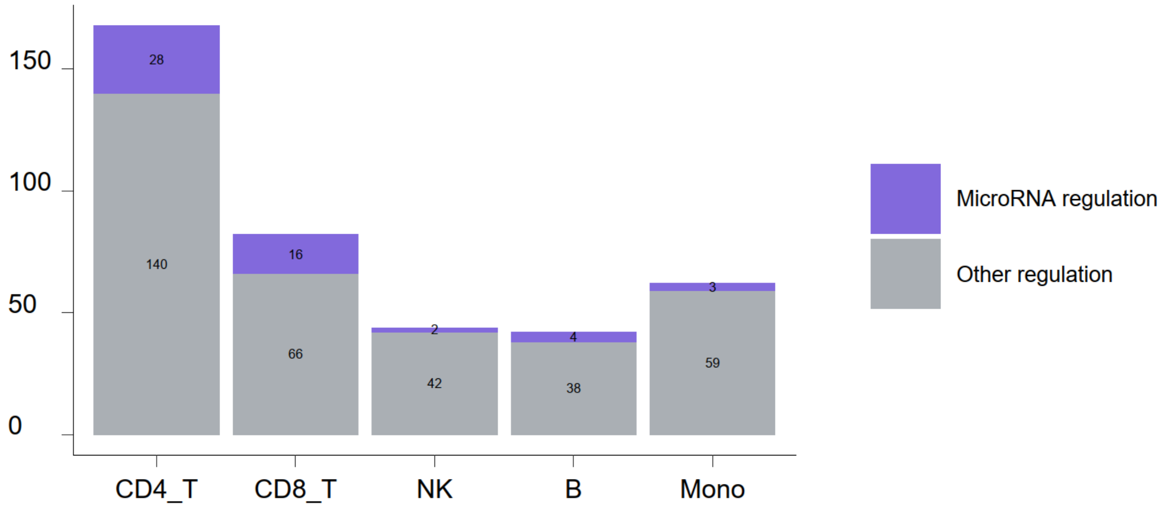


**Suppl. Fig. 18:** Proportion of eGenes per cell type with evidence of being regulated by microRNAs.

Suppl. Fig 19

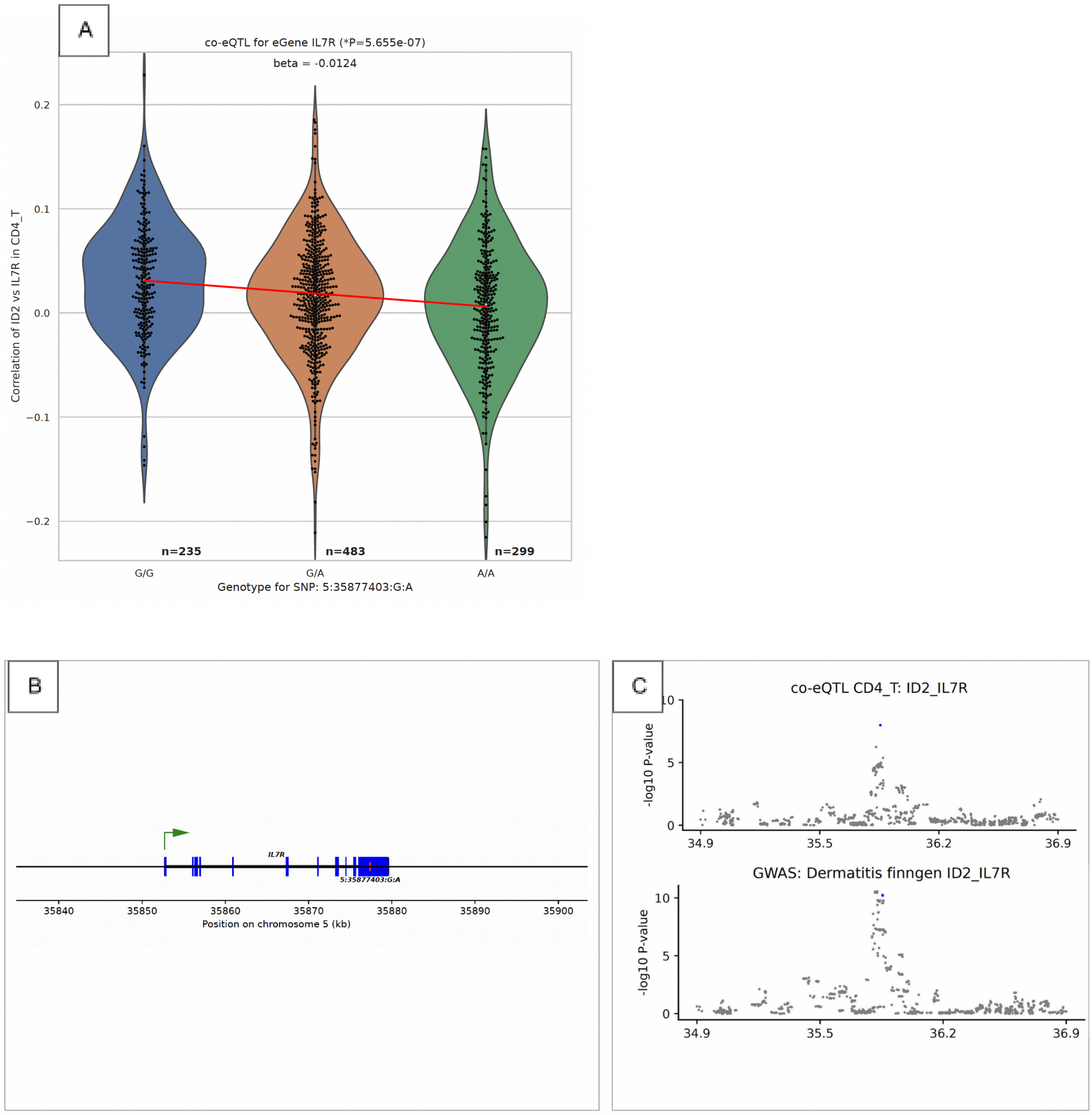


**Suppl. Fig. 19: a)** Correlation co-eQTL plot showing the co-eQTL effect size of 5:35877403:G:A on the correlation between *IL7R* and *ID2* in CD4+ T cells, based on effect sizes in OneK1K. **b)** Location of variant in relation to IL7R, exons shown in blue. **c)** Locus zoom plot showing both the dermatitis GWAS and co-eQTL signals in the locus. The variant 5:35877403:G:A is highlighted in blue. A strong colocalization is observed between these signals (H4 = 0.97). The protein-coding gene landscape is shown below the co-eQTL plot.

Suppl. Fig. 20

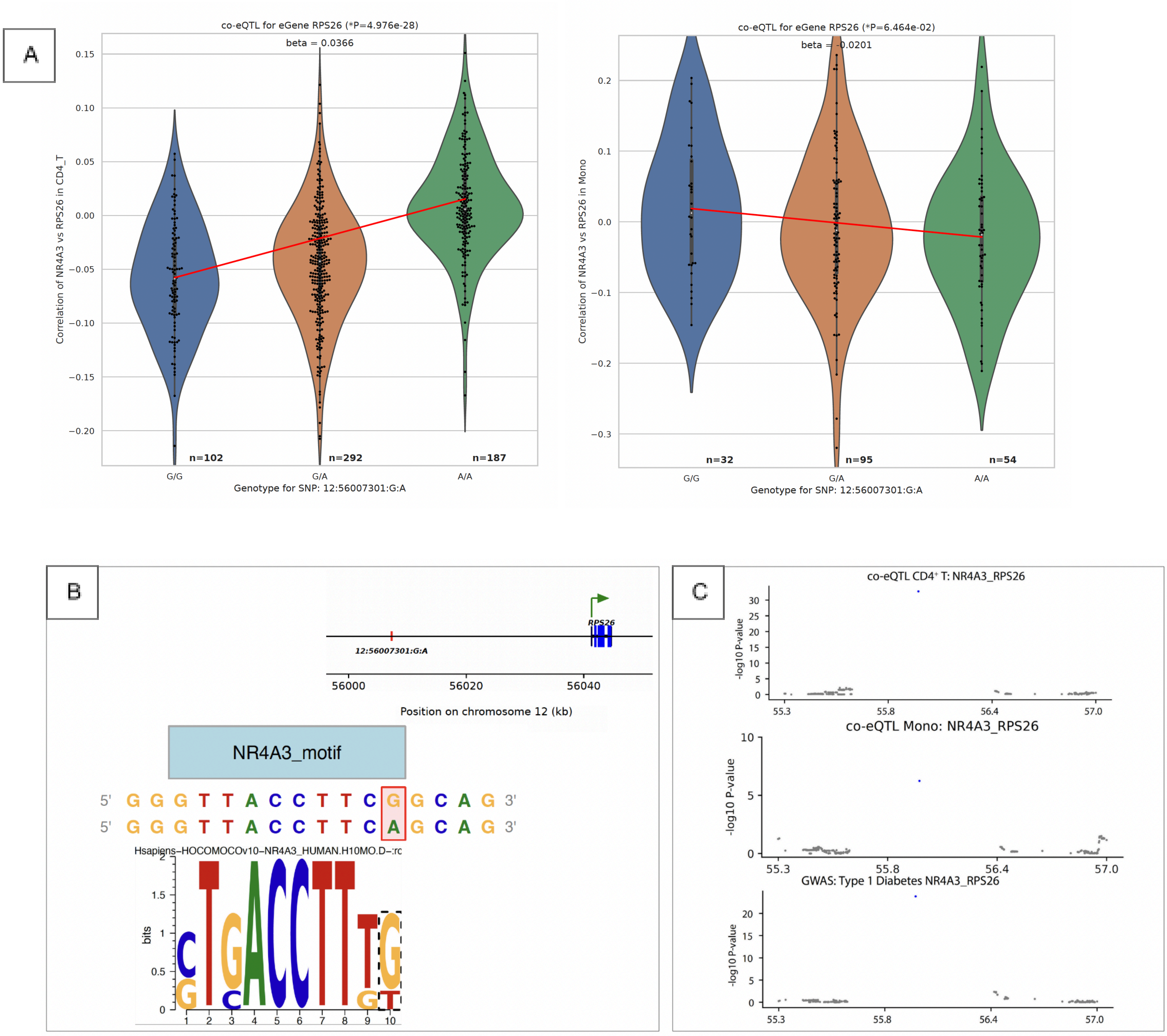


**Suppl. Fig. 20: a)** Correlation co-eQTL plots showing the co-eQTL effect size of 12:56007301:G:A on the correlation between *RPS26* and *NR4A3*, based on effect sizes in OneK1K in CD4+ T (left) and monocyte (right) cells. **b)** Motif plot from MotifbreakR. G is shown to match the preferred binding motif for IKZF2, whereas A breaks the motif. **c)** Locus zoom plot showing both the type 1 diabetes GWAS and co-eQTL signals in the locus. The variant 12:56007301:G:A is highlighted in blue. A strong colocalization is observed between these signals (H4 = 0.996). The protein-coding gene landscape is shown below the co-eQTL plot.

Suppl. Fig. 21

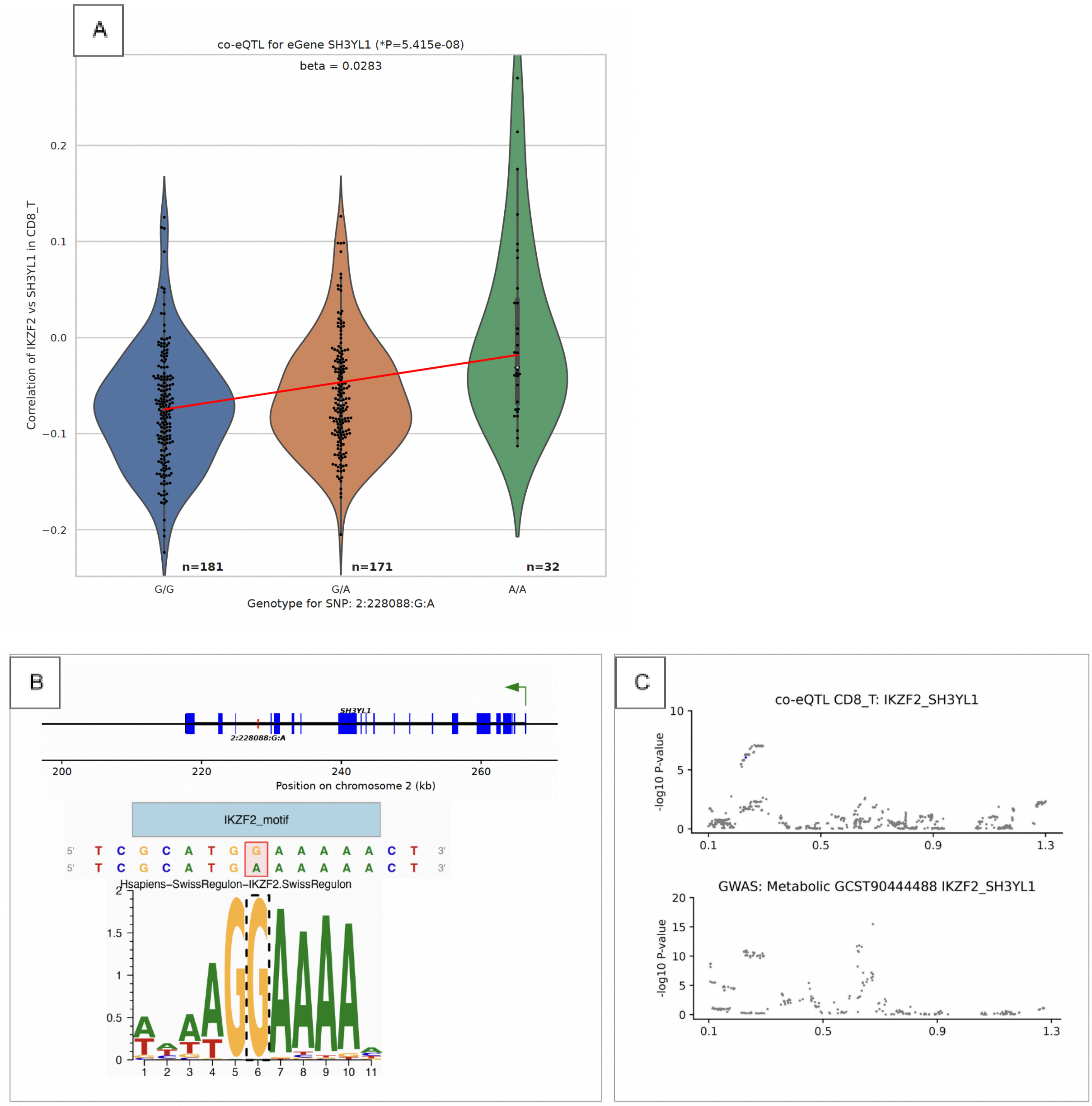


**Suppl. Fig. 21: a)** Correlation co-eQTL plot showing the co-eQTL effect size of 2:228088:G:A on the correlation between *SH3YL1* and *IKZF2* in CD8+ T cells, based on effect sizes in OneK1K. **b)** Motif plot from MotifbreakR. G is shown to match the preferred binding motif for *IKZF2*, whereas A breaks the motif. **c)** Locus zoom plot showing both the Metabolic disease GWAS and co-eQTL signals in the locus. The variant 2:228088:G:A is highlighted in blue (variant not included in GWAS). A strong colocalization is observed between these signals (H4 = 0.96). The protein-coding gene landscape is shown below the co-eQTL plot.

Suppl. Fig. 22

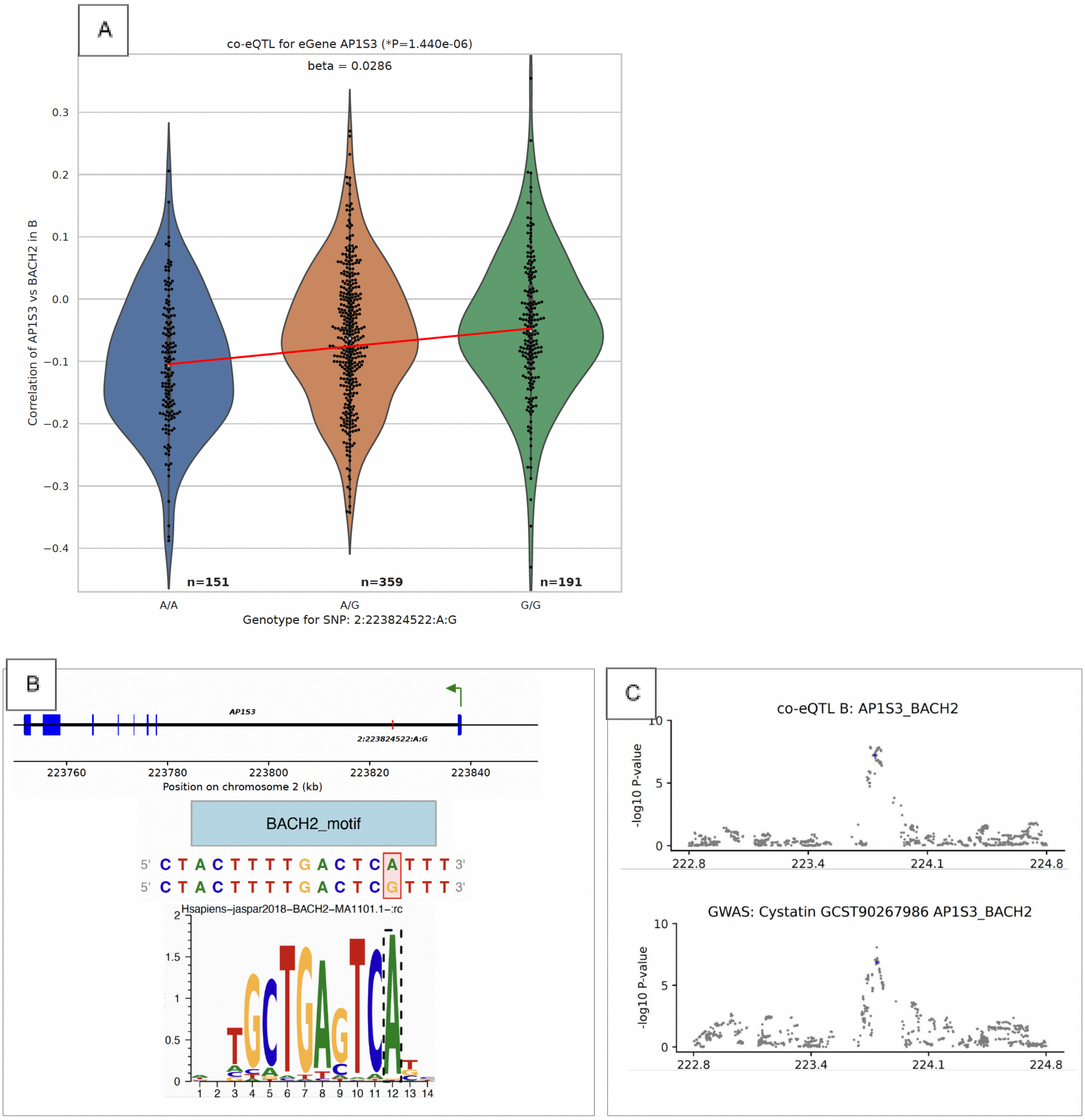


**Suppl. Fig. 22: a)** Correlation co-eQTL plot showing the co-eQTL effect size of 2:223824522:A:G on the correlation between *AP1S3* and *BACH2* in B cells, based on effect sizes in OneK1K. **b)** Motif plot from MotifbreakR. A is shown to match the preferred binding motif for *BACH2*, whereas G breaks the motif. **c)** Locus zoom plot showing both the Cystatin C GWAS and co-eQTL signals in the locus. The variant 2:223824522:A:G is highlighted in blue. A strong colocalization is observed between these signals (H4 = 0.85). The protein-coding gene landscape is shown below the co-eQTL plot.
